# The psychophysics of visual search with heterogeneous distractors

**DOI:** 10.1101/2020.08.10.244707

**Authors:** Andra Mihali, Wei Ji Ma

**Affiliations:** Center for Neural Science, New York University, New York, NY 10003, USA; Department of Psychology, New York University, New York, NY 10003, USA; Research Foundation for Mental Hygiene, New York State Psychiatric Institute, New York, NY 10032, USA; Department of Psychiatry, Columbia University, New York, NY, 10032, USA

## Abstract

Visual search is one of the most ecologically important perceptual task domains. One research tradition has studied visual search using simple, parametric stimuli and a signal detection theory or Bayesian modeling framework. However, this tradition has mostly focused on homogeneous distractors (identical to each other), which are not very realistic. In a different tradition, Duncan and Humphreys (1989) conducted a landmark study on visual search with heterogeneous distractors. However, they used complex stimuli, making modeling and dissociation of component processes difficult. Here, we attempt to unify these research traditions by systematically examining visual search with heterogeneous distractors using simple, parametric stimuli and Bayesian modeling. Our experiment varied multiple factors that could influence performance: set size, task (N-AFC localization vs detection), whether the target was revealed before or after the search array (perception versus memory), and stimulus spacing. We found that performance robustly decreased with increasing set size. When examining within-trial summary statistics, we found that the minimum target-to-distractor feature difference was a stronger predictor of behavior than the mean target-to-distractor difference and than distractor variance. To obtain process-level understanding, we formulated a Bayesian optimal-observer model. This model accounted for all summary statistics, including when fitted jointly to localization and detection. We replicated these results in a separate experiment with reduced stimulus spacing. Together, our results represent a critique of Duncan and Humphrey’s descriptive approach, bring visual search with heterogeneous distractors firmly within the reach of quantitative process models, and affirm the “unreasonable effectiveness” of Bayesian models in explaining visual search.

## Introduction

People often perform visual search in daily life, such as when detecting the face of a friend in a group or locating the keys on a table among various other objects. A topic of broad interest in visual search research is what factors influence performance, which is usually measured through reaction times or accuracy. Visual search performance can vary with the number and complexity of objects, their similarity to neighboring objects, the type of information that is needed from the search (i.e - whether an item is present or not, or where an item is located), as well as whether one searches for an object by looking at the environment, or by recalling the visual environment from memory (for reviews, see Wolfe (2010); Eckstein (2011)).

In the laboratory, early visual search studies typically used simplified stimuli such as letters or shapes, and addressed the question of how reaction time depends on the number of items (*set size*). Perhaps the best-known line of visual search work started with Treisman and Gelade (1980). Treisman and Gelade (1980) formulated the feature integration theory (FIT) of attention to explain the result that reaction times were approximately constant with set size for feature searches (for example, a T in a display of letters) but increased with set size for conjunction searches (for example, the green T). FIT posits that in a pre-attentive stage, all features are processed in parallel and in a second stage, selective attention is engaged serially to bind these loose features into joint objects, so that the observer can ultimately determine whether the target is present.

### Visual search with heterogeneous distractors

Duncan and Humphreys (1989) questioned the distinction between feature and conjunction search and between parallel and serial search and instead proposed that search falls on a continuum of efficiency, which depends not on the features of the stimuli per se, but on their similarity to one another. To obtain variation in stimulus similarity, they employed displays with heterogeneous distractors. At the time, there was some earlier evidence that increasing distractor heterogeneity increased search times (Gordon, 1968; Gordon, Dulewicz, & Winwood, 1971; Farmer & Taylor, 1980). Duncan and Humphreys (1989) performed 6 search experiments with letter displays and used the results to build a general proposal in which search performance: 1) decreases as target-distractor similarity increases; 2) decreases as distractor-distractor similarity within a display decreases (or, in other words, distractor heterogeneity increases). These two factors interact: specifically, higher distractor heterogeneity decreases search performance mainly when the target-distractor similarity is also high. They captured the factors and their interactions visually in a “search surface” (Figure 3 of Duncan and Humphreys (1989)).

Still, the question of how exactly to best quantify target-distractor similarity and distractor heterogeneity remains open. The empirical results of Duncan and Humphreys (1989) only had a coarse divide into homogeneous and heterogeneous; granted heterogeneity in letter displays is not straightforward to quantify. In a related experiment, Duncan (1989) found a similar pattern of results in search with color arrays, which lend themselves more naturally to quantification of distractor heterogeneity; however their measures of heterogeneity were still coarse (distractor set having 0, 1 or 2 neighbors of the target in a 6 point color set). With variable targets, Nagy, Neriani, and Young (2005) found that in color search, performance was impaired with heterogeneous distractors relative to homogeneous distractors, result in line with the proposal of (Duncan & Humphreys, 1989). However, other studies found either an increase, or no effect on performance with increasing distractor heterogeneity (Nagy & Thomas, 2003; Vincent, Baddeley, Troscianko, & Gilchrist, 2009). These inconsistencies in quantification of distractor heterogeneity as well as conclusions justify further investigation. Here, we will revisit the question of how distractor heterogeneity influences visual search performance and reframe this question with a quantitative modeling framework.

### Modeling approaches to visual search

To achieve even greater experimental control of visual search in the laboratory, a group of researchers introduced further simplifications in experimental design: unidimensional stimuli that can be easily parametrized, brief stimulus presentation times to prevent eye movements, placement of stimuli at equal eccentricities from the center of eye fixation, and wide spacing between stimuli (reviewed in Palmer, Verghese, and Pavel (2000)). In these designs, the effect of set size on performance is not usually characterized through reaction times, but through accuracy, or related measures, such as stimulus difference threshold (Palmer, 1990). A large advantage of these simplifications is that they make the data amenable to quantitative modeling, in particular within the signal detection theory (SDT) framework which allows for the dissociation of perceptual encoding and decision processes (Peterson, Birdsall, & Fox, 1954; Green & Swets, 1966). SDT has been routinely applied in psychophysics to the detection or categorization of a stimulus in noise (Green & Swets, 1966). SDT was extended to the more complex task of visual search by specifying ways to combine the information from the noisy representation of the stimuli, such as using a maximum-of-outputs rule (Green & Swets, 1966; Nolte & Jaarsma, 1967; Palmer, Ames, & Lindsey, 1993; Palmer et al., 2000; Eckstein, Thomas, Palmer, & Shimozaki, 2000; Shaw, 1982; Cameron, Tai, Eckstein, & Carrasco, 2004; Verghese & Nakayama, 1994; Verghese, 2001), a heuristic-probabilistic rule (Vincent et al., 2009) or a probabilistically-optimal (Bayesian) rule (Ma, Navalpakkam, Beck, van den Berg, & Pouget, 2011; Ma, Shen, Dziugaite, & van den Berg, 2015; Shen & Ma, 2019), the latter being the optimal-observer model.

In most studies using SDT modeling of visual search, the distractors were *homogeneous,* meaning identical to each other within a display, and their identity was kept constant across trials (Bergen & Julesz, 1983; Palmer, 1990; Palmer et al., 1993; Palmer, 1994; Palmer et al., 2000; Eckstein et al., 2000; Shaw, 1982; Cameron et al., 2004; Verghese & Nakayama, 1994). When the distractors are homogeneous, the difference between the target and distractors can be represented by a single number, which has the advantage of simplifying calculations. The experimental result that accuracy decreases with set size can be captured with SDT modeling. The intuition is that the higher the number of distractors, the higher the chance that the noisy representation of one of them might get mistaken for the target.

SDT was additionally able to capture how accuracy (or related measures) degrades with target-distractor similarity (Shaw, 1982; Palmer et al., 1993; Eckstein et al., 2000; Verghese, 2001). This has been demonstrated across multiple tasks (target detection, localization and identification (Palmer et al., 1993; Palmer, 1994; Palmer et al., 2000; Eckstein et al., 2000; Shaw, 1982; Cameron et al., 2004)), memory demands (both perception and memory conditions Palmer (1990)), and stimulus types (line length, orientation, luminance, shape (Palmer et al., 1993; Palmer, 1994), speed (Verghese & Stone, 1995); but see Verghese and Nakayama (1994)). Moreover, the application of SDT models to visual search has refined ideas about the role of attention in visual search (Palmer, 1994; Verghese, 2001; Cameron et al., 2004).

SDT modeling of visual search has mainly focused on homogeneous distractors and with this approach something important has been lost, namely ecological relevance. In the real world, distractors are largely heterogeneous. For instance, when searching for a particular key on a key chain or one’s car in a parking lot, the non-target keys or cars will likely not be all identical, but have various different shapes and/or colors, akin to a set of heterogeneous distractors. Of course, many objects in the real world are even more complex, but distractor heterogeneity is one naturalistic aspect that can be feasibly incorporated in models in the signal-detection theory framework.

One of the first attempts to bridge the study of visual search with heterogeneous distractors with SDT quantitative modeling was made by Rosenholtz (2001). SDT predicts that distractor heterogeneity would impact visual search as follows: if the heterogeneity of the distractors would make them more similar to the target, performance would worsen, but if it would make them more discriminable than the target, performance would improve (Rosenholtz, 2001; Vincent et al., 2009). Only the SDT prediction in the first case is in line with (Duncan & Humphreys, 1989). Rosenholtz (2001) extended the usual form of the signal detection theory model to account for more possibilities of target and distractor values. Even with ideal experimental conditions for an SDT model (sparse displays etc), the performance decrement with heterogeneous distractors was worse in the human observers than in the SDT model predictions. A limitation of this study was that the heterogeneity of the distractors could not be quantified unambiguously; the distractor distributions were non-smooth (discrete) distributions with up to 3 parameters, making them potentially hard to learn (Mazyar, van den Berg, Seilheimer, & Ma, 2013).

Standard SDT is a particular case of Bayesian decision theory, which can accommodate a wider range of assumptions about how stimuli yield measurements beyond the Gaussian assumption from standard SDT. In the Bayesian decision theory framework, one can compute trial-by-trial predictions of responses for given stimuli. Optimal-observer models are instantiations of Bayesian decision theory for given tasks. In another sense, optimal-observer models are more particular than SDT models as they have a unique way to combine information across all the stimuli; this unique way is the Bayesian decision rule, derived by taking into account the statistical distribution of the stimuli and making plausible explicit assumptions, for instance about task operations, and the noise distribution on encoding (see Ma (2012) for clarifications on terminology). Thus, if such assumptions are justified, these models become good candidates to capture visual search performance with heterogeneous distractors. Using a task of target detection in search arrays with oriented patches, Ma et al. (2011) found that an optimal-observer model captured human behavior. This was the case both for homogeneous distractors and heterogeneous distractors drawn from a uniform distribution; however, this work did not vary set size. In a similar task, Mazyar, van den Berg, and Ma (2012) used heterogeneous distractors drawn from a uniform distribution (maximally unpredictable). They varied set size in a target detection task and found that an optimal-observer model only fitted well if the additional allowance was made that the mean encoding precision of a given item varied with set size in a decreasing fashion. This pattern of results held true regardless of whether the identity of the target was revealed before or after the search array, meaning for both visual search in perception and in memory. Calder-Travis and Ma (2020) extended the study of target detection in visual search with heterogeneous distractors by employing two types of distractor distributions (uniform and von Mises) and also found that an optimal-observer model captured the data well.

## Goals of the present study

Here, we further extended the optimal-observer modeling approach to visual search with heterogeneous distractors with four goals: 1) to assess in a comprehensive fashion which factors related to distractor heterogeneity influence performance, 2) to put this optimal-observer model to a thorough test and 3) to revisit questions that have previously been asked outside of a modeling context, such as search patterns across the tasks of localization and detection and 4) search patterns in perception and memory.

To approach the first goal of achieving a comprehensive characterization of visual search with heterogeneous distractors, we took advantage of the rich variation that naturally occurs in displays consisting of independently drawn distractors, here from a uniform distribution. While in visual search with homogeneous distractors target-distractor similarity can be captured with a single number, in the case of heterogeneous distractors we have more flexibility in measuring trial difficulty or confusability. Intuitively, on a given trial, performance might be impaired if: a distractor is highly similar to the target, the average of the distractors is similar to the target, or the distractors are highly variable. Therefore, we characterized search behavior as a function of the following summary statistics which naturally emerge from the uniform distribution, computed on a trial-by-trial basis: the difference between the target and the most similar distractor (Mazyar et al., 2012, 2013) the difference between the target and the mean of the distractors, and distractor variance. Only one study so far has investigated all these summary statistics (Calder-Travis & Ma, 2020). Because we use a unidimensional, continuous stimulus variable (orientation), these quantities can be defined exactly and unambiguously. Each of these summary statistics gave rise to a set of psychometric curves (one for each set size).

Beyond addressing the first goal, these rich summary statistics also served towards our second goal of providing a thorough test for the optimal-observer model. For the application of the optimal-observer model to be successful in capturing the data, the observers need to have knowledge of the distribution of stimuli; this is a plausible assumption in the case of a simple distribution easy to learn such as the uniform (vs, for instance, a less common distribution with several parameters). Additionally, it is possible for a model to fit well to coarse summary statistics (such as performance as a function of set size) but fail to fit a further breakdown of the data. Thus we set to subject the optimal-observer model to a more thorough test.

The third goal consisted of revisiting a question from more qualitative work in visual search, namely the relation between search behavior in target localization versus target detection. In the real world, localization might be a more frequent task than detection (Dukewich & Klein, 2009). For instance, you might need to not just detect, but locate your desired key on a keychain or your car in a parking lot several times per day. Detection and localization have been studied together, sometimes even in the same observers, but only in the context of homogeneous distractors, (Liu, Healey, & Enns, 2003; Cameron et al., 2004; Busey & Palmer, 2008; Dukewich & Klein, 2009). As pointed out by Liu et al. (2003), visual search theories largely took for granted that localization and detection are linked, with some suggesting that detection is solved through localization, and others the other way around. Liu et al. (2003) emphasized a dual-systems approach (Goodale & Milner, 1992) to localization and detection and showed some evidence that direct localization might rely on the “how” dorsal stream, involved in perception for action and with higher spatial resolution, while detection might rely on the “what” ventral stream; still there is substantial overlap between these two streams, especially in early encoding. Here, we took advantage of the fact that the optimal-observer models for detection and localization are highly similar in the early encoding stages (Ma et al., 2011). As the same observers performed both tasks, we tested a version of the model assuming shared encoding precision for localization and detection, in a sense probing whether they are reliant on shared early representations.

The fourth goal concerned potential differences between *perception-based* and *memory-based* visual search. Studies have suggested that searching in the visual environment shares similarities with searching in one’s memorized internal representations of the objects in the visual environment (Kuo, Rao, Lepsien, & Nobre, 2009; Kong & Fougnie, 2019). There is evidence that the orienting of attention *externally* (in perception) works through similar mechanisms as the orienting of attention *internally,* within visual short-term memory (Baddeley & Hitch, 1974; Kiyonaga & Egner, 2012). Such orienting of attention in memory can be spatially driven, but could also be based on a feature dimension value (i.e. “Search the array of colors in your memory and determine whether there is a blue one”). Kuo et al. (2009) employed such a task and found that the behavior and neural correlates were largely similar between perception and memory-based visual search. Other behavioral studies also found that patterns such as the set size effect and the target-nontarget similarity effect were shared between perception and memory-based search (Mazyar et al., 2012; Kong & Fougnie, 2019). Studies found a decrease of accuracy (Kuo et al., 2009) and precision (Mazyar et al., 2012) with set size in both perception and memory. Accuracy (Kuo et al., 2009; Mazyar et al., 2012), as well as precision (Mazyar et al., 2012) were higher in perception than in memory. Here, we revisit these questions in target detection with heterogeneous distractors and across more detailed performance metrics, as well as extend them to another search task, target localization.

The paper is organized as follows. We first explored how visual search performance varies with the summary statistics above. We report the fits of an optimal-observer model to the localization and detection datasets, first separately and then jointly, allowing us to test whether the precisions with which items are encoded can be shared across localization and detection. Next, we conducted a control experiment to examine whether our conclusions hold at a lower stimulus spacing. Finally, we compared the precisions of visual representations across our experimental conditions and experiments.

## Experimental methods

The task was either *N*-alternative forced-choice (*N*-AFC) localization or detection in different blocks. *N*-AFC localization, which we refer to as localization, allowed for more informative responses than 2-AFC detection. We considered two possible temporal orders of target and search array, such that the search was either based on perception (by having the target orientation revealed before the search array) or short-term memory (by having the target orientation revealed after the search array). Every participant performed both tasks and both temporal order conditions.

### Experiment 1

#### Participants

11 participants (9 female, 2 male) performed the task upon providing informed consent. The study adhered to the Declaration of Helsinki and was approved by the NYU Institutional Review Board.

#### Apparatus

We displayed stimuli on a Dell 1907FPC LCD monitor with resolution 1280 × 960 pixels and width 19” (with 16.1” viewable) and 75 Hz refresh rate. A Windows computer running Matlab 8.2 (MathWorks, Massachusetts, USA) with Psychtoolbox 3 (Brainard, 1997; Pelli, 1997; Kleiner et al., 2007) displayed the stimuli. Subjects were located at approximately 60 cm from the screen. The screen background was mid-level gray.

#### Stimuli

On each trial, target and distractor orientations were drawn independently from a uniform distribution on 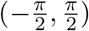. Each stimulus was an oriented Gabor patch of spatial frequency 2.85 cycles/o with standard deviation 0.26 degrees of visual angle (dva) subextending an image size of height and width both equal to 1.3 dva. Participants viewed search displays in order to detect or locate the item that matched the orientation of the centrally presented target stimulus. The number of stimuli in the search display was 2, 3, 4 or 6 and they were randomly interleaved. Since the search display stimuli were placed adjacently at 60° angular intervals on an imaginary circle and eccentricity was 5 dva, the center-to-center distance between two adjacent Gabors was 5 dva.

### Procedure

#### Localization

Each trial in the Localization-Perception condition began with a central fixation cross (diameter 0.15 dva) for 500 ms, followed by a centrally located target Gabor for 100 ms, a blank screen for 1000 ms, and then a search display for 100 ms (Figure 1A). The short stimuli presentation times served to minimize the effect of eye movements. In localization, the target was always present in the search display. The location of the target in the display was chosen randomly from the stimulus locations. Once the response screen appeared, participants had to use the mouse to click on the location where the stimulus orientation matched the target orientation. The possible location options were *N* white circles corresponding to the original locations of the stimuli. Thus, the localization task was *N*-AFC and the probability of responding correctly by chance was 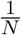.

**Figure 1:**
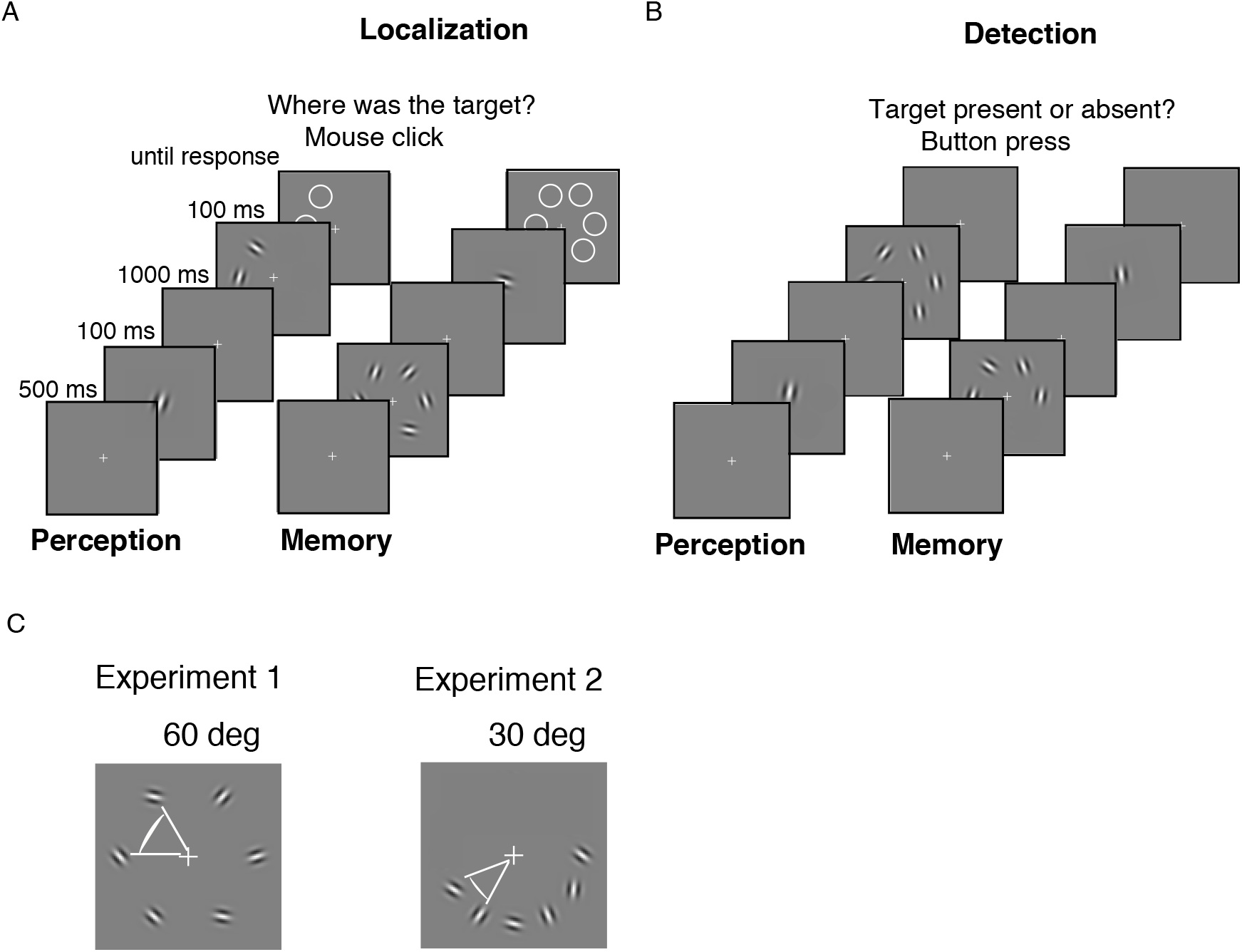
Task design. Trial sequence for **(A)** Localization and **(B)** Detection. Across both Localization and Detection, observers performed both a Perception condition, in which the search target was revealed before the search array, and a Memory condition, with the target revealed after the search array. In **(A)** and **(B)**, the stimuli are spaced as in Experiment 1, 60 deg apart. **(C)** The spacing of the stimuli in the search array in the Experiment 1 vs Experiment 2, depicted here for the highest set size 6. Experiment 2 is almost the same as Experiment 1, with the only difference that stimuli are spaced 30 deg apart.

The Localization-Memory condition was identical to the Localization-Perception condition except that the temporal order of the target and the search display was reversed. Both perception and memory conditions required observers to engage memory to some extent, but while in the perception conditions observers only had to remember the orientation of one Gabor over the delay period, in the memory conditions observers had to remember all *N* Gabors over the delay period.

##### Detection

While in localization the target was always present in the search display, in detection the target was present in the display with a probability of 0.5. On a target-present trial, the orientation of one of the items in the search display matched the orientation of the target Gabor. Subjects reported whether or not the target orientation matched the orientation of any item in the search display. Subjects indicated their responses by pressing one of two possible keys, located nearby on a small keyboard (thus the task could be performed with 2 different fingers of their dominant hand). Thus, the probability of responding correctly by chance in Detection was 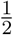 regardless of set size. The trials in Detection-Perception and Detection-Memory conditions followed the same sequence as in the analogous localization conditions.

##### Blocks and sessions

The full experiment consisted of four sessions of 800 trials each. Each session was divided into 8 blocks. At the end of each block, the participants were informed of their performance scores so far. There were four conditions: Detection-Perception, Localization-Perception, and respectively Detection-Memory and Localization-Memory. Thus, we obtained in total 800 trials per each condition, (200 trials per each set size). In the first four blocks, we randomly chose the order of the conditions for each participant; we mirrored that order in the second half (palindrome construction). This order of blocks was maintained across the four sessions for that participant.

##### Instructions

Before the start of the first block on the first session, participants were informed about the procedure. In particular, we emphasized to them that while in Detection there is a 0.5 probability that the target is present in the search display, in localization the target is always present in the search display. Before starting the experiment, participants performed a total of 40 training trials, 10 for each of the 4 conditions we detail below: Localization-Perception, Localization-Memory, Detection-Perception and Detection-Memory.

##### Data and code

Experiment, analysis and modeling code are available at the following github repository: https://github.com/lianaan/Vis_Search.

## Experiment 2

Experiment 2 was almost identical to experiment 1 with just one key difference: the search display stimuli were placed adjacently at 30 ° angular intervals, and thus two adjacent Gabors were spaced at 2.6 dva center-to-center distance (Figure 1C). The rest of the methods are identical to what we described for Experiment 1, with the additional differences listed below.

### Participants

6 participants (3 female, 3 male) performed the task.

### Apparatus

In Experiment 2, we displayed stimuli on a ViewSonic VX2475Smhl-4K LED monitor with resolution 3840 × 2160 pixels and width 24” (with 23.1” viewable) and 60 Hz refresh rate. For this monitor, the screen background was also mid-level gray.

## Models

We can see that the models for localization and detection share a similar architecture (Figure 2). The generative model for localization is presented in Figure 2A.

**Figure 2:**
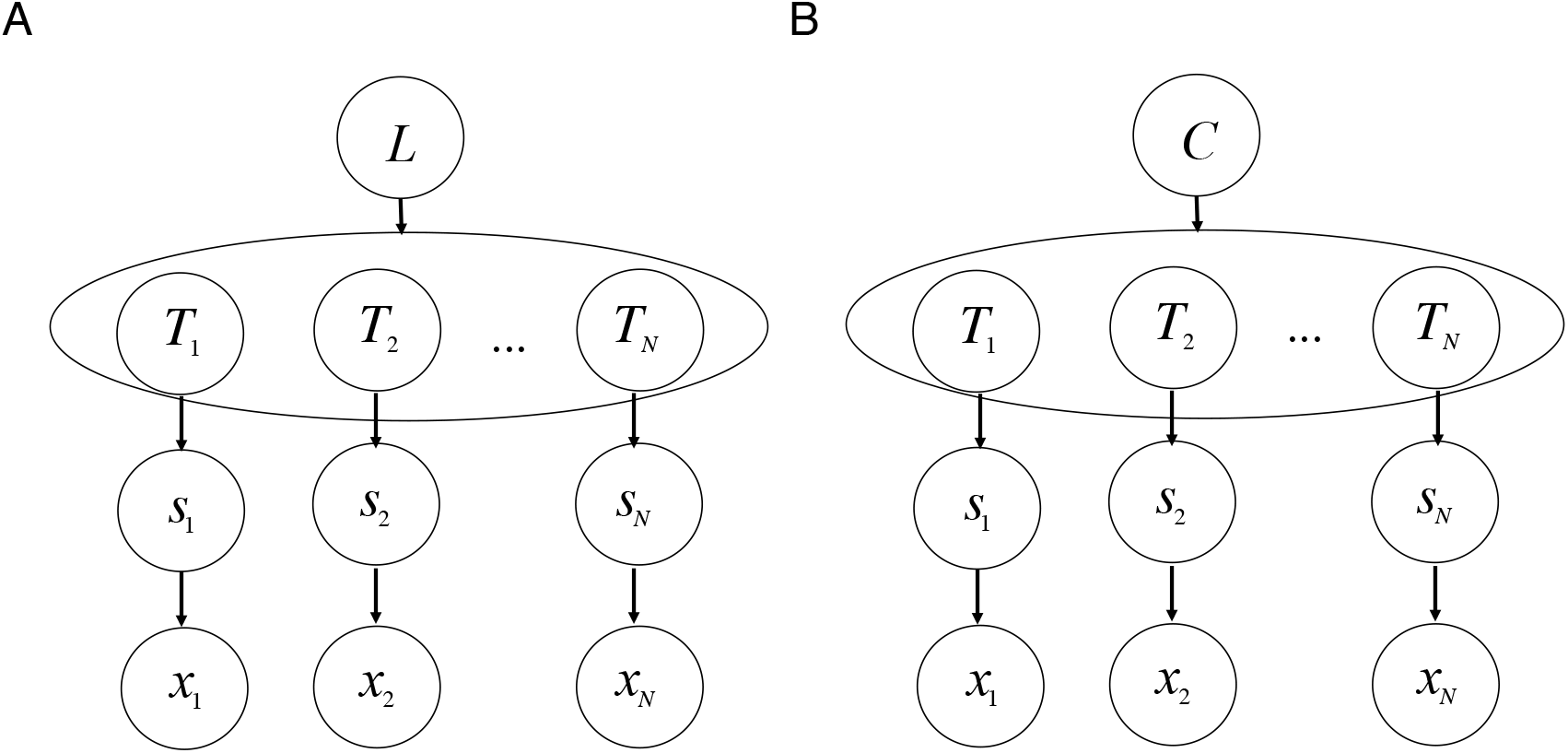
The generative models for **A)** Localization and **B)** Detection share a common structure. Each node represents a variable and each arrow a link between the variables. These links are explicitly presented in the text. For localization, *L* can take values 1,2,… *N* and *T_L_* = 1, while the rest of the *T_i=L_* = 0. For detection, as *L* can take values 0 and 1, we refer to it by *C*. If *C* = 1, *T_L_* = 1 and *T_i≠L_* = 0. However, if *C* = 0, all *T_i_* = 0.

**Figure 3:**
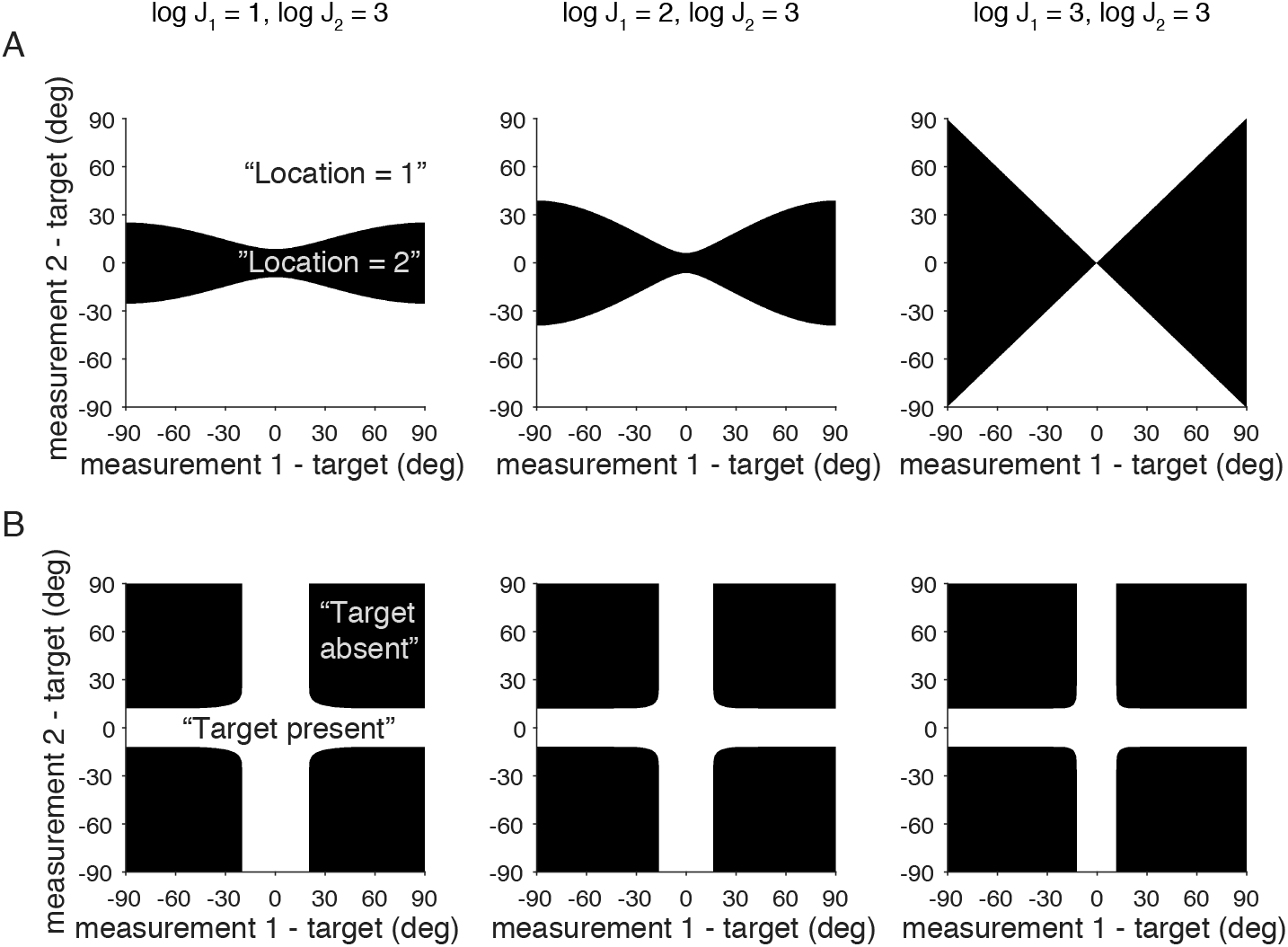
Visualization of the optimal decision rules for the case of 2 items. The decision rule is depicted for each possible combination of measurements *x*_1_ and *x*_2_ relative to to the target value separately for: **A)** Localization : if *d*_1_ > *d*_2_, respond “location 1” (white) versus “location 2” (black) and **B)** Detection: if global *d* > 0 respond “target present” (white) vs “target absent” (black). The columns differ in the combinations of precision values associated with measurements 1 and 2. Intuitively, for localization, the higher the precision of one item, the narrower the region around the target in which this item will be chosen. Intuitively, for detection, the higher the precision of one item, the narrower the region around the target for which the observer reports “target present”. For this visualization we used the range [—180, 180) deg for the measurement - target difference, which we converted back to [–90, 90) deg for visualization.

### Optimal-observer model for Localization

#### Step 1. Generative model / encoding

We extend the optimal-observer framework (Ma et al., 2011; Mazyar et al., 2012) to the *N*-AFC localization task here. Here we fit the optimal-observer model with variable encoding precision as we consider it the most canonical model (van den Berg, Shin, Chou, George, & Ma, 2012; Mazyar et al., 2012).

The generative model is presented in Figure 2A. We denote the location of the target with *L*, which can take one of the values 1,2,… *N*, with *N* being set size (2, 3, 4 or 6). Any location is equally likely such that the prior over the locations for a given set size *N* is: 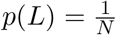. Then, the target will be present at location *L*, but nowhere else: *T_L_* = 1, *T_i≠L_*, = 0. We denote **T** = (*T*_1_,*T*_2_, …*T_N_*) and can write formally *p*(**T**|*L*) = *δ*(**T** – 1_*L*_). Each orientation stimulus *s_i_* (with *i* = 1, 2,…*N*) follows a uniform distribution 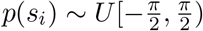; same does *p*(*s_L_*|*L*). We assume that the stimuli *s_i_* are each independently encoded as the noisy observations *x_i_* and thus:

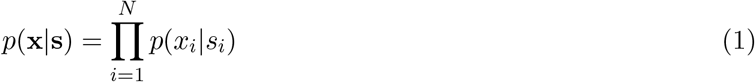

In particular, we assume the noise follows a von Mises distribution with concentration parameter *κ_i_* such that *p*(*x_i_*) ~ *VM*(*s_i_,κ_i_*). The support of the von Mises distribution is [–*π,π*); we thus remap all orientations *s_i_* from 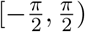 to [–*π,π*) in our models and analyses. Then, we can write:

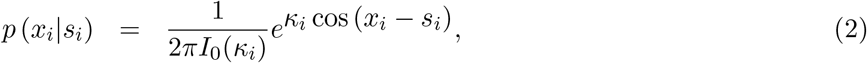

where *I*_0_ is the modified Bessel function of the first kind of order 0. To see how the concentration parameter *κ* is related to precision *J*, see Appendix A.1.

An additional assumption in the variable-precision encoding model (van den Berg et al., 2012) is that the precision *J* varies across trials and items, specifically according to a Gamma distribution with shape parameter 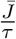 and scale parameter *τ*, and thus with mean 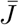.

#### Step 2. Decision rule

In this step we show the decision rules that we assume the optimal-observer employs to produce their responses. To eventually report their inferred location, 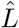, the observer has to invert the generation model and compute the posterior over the possible locations: *p*(*L*|**x**). This entails applying Bayes’ rule to the prior *p*(*L*) and the likelihood *p*(**x**|*L*):

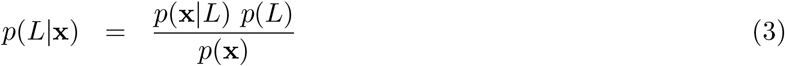

The computation of *p*(**x**|*L*) is guided by the generative model in Figure 2A. We first write out *p*(**x**|*L*) by marginalizing over the stimuli, factorizing across items and segregating the target or otherwise the item at location *L* from the distractors:

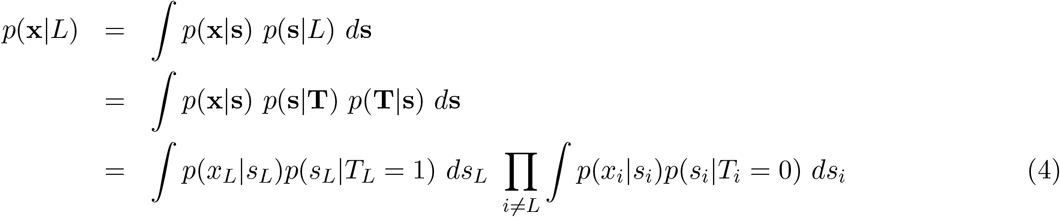

For further simplification, we re-organize the factors, multiply and divide by a factor for a possible distractor located at location *L* and then make use of the facts that *p*(*s_i_*) is uniform on 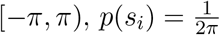 and 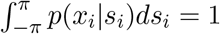.

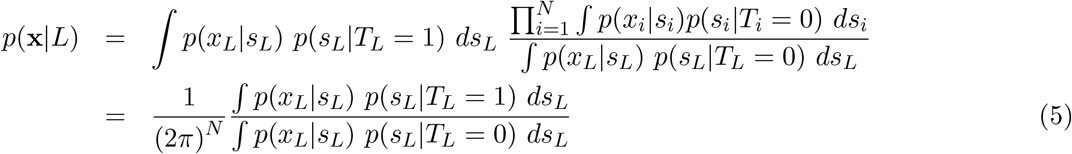

Here, we recognize the log ratio of the probability of the local measurement at position *L*, *x_L_*, coming from the target *s_T_* relative to the probability of *x_L_* coming from any other distractor, *s_L_*. We refer to this term as the log decision variable *d_L_*:

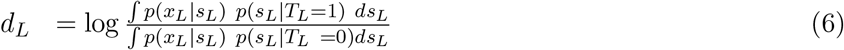

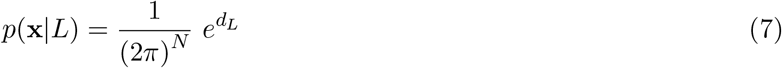

We unpack Equation 6 (See Appendix A.2) and get:

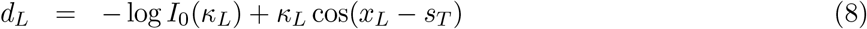

We assume the optimal observer reports their response 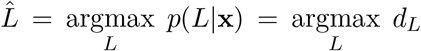. For full derivations, see Appendix A.2.

Below we also consider a model in which the observer does not reliable respond with the location corresponding to the maximum of the posterior *p*(*L*|**x**), but instead their responses are affected by decision noise (Model 2).

We emphasize a particular case in which an item at location *i* is encoded with 0 precision such that = 0 and thus *d_i_* = 0. If the set size is *N* = 2, the decision rule for choosing the other item *j* becomes a threshold: *j* will be chosen if *d_j_* > 0.

#### Step 3. Model predictions

As we saw above, the decision rule provides a prescription to compute observer’s responses 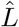 when their internal measurements **x** are known. But on every trial, it is only known that the observer sees a set of stimuli **s**. Thus, to find the predicted responses we need to compute the integral 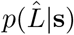. To do this, we have to marginalize over all the possible measurements **x**:

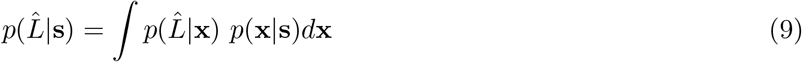

We computed the probability of response 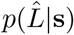 by estimating the above integral by sampling: we simulated several samples (here *N*_samples_ = 1200) of **x** from **s** and averaged the predictions over the corresponding outcomes 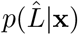.

### Optimal-observer model for Localization with decision noise : Model 2

In the form outlined above, the optimal-observer model assumes that the observer calculates and utilizes in a deterministic way the decision variable from Step 2 of the generative model. However, some studies considered the possibility that the decision criterion in signal detection theory (or the decision variable in the optimal-observer model) varies itself on a trial-by-trial basis (Mueller & Weidemann, 2008; Drugowitsch, Wyart, Devauchelle, & Koechlin, 2016; Shen & Ma, 2019; Stengård & van den Berg, 2019). The decision noise is thus the parameter that captures the variability in response for a given decision variable.

Such decision or inference noise would lead the observer to no longer respond systematically with the MAP estimate. Various ways of modeling the decision noise have been proposed. Some ways of adding decision noise are specific to binary tasks, such as softmax. Here we use a very general way proposed by Acerbi, Vijayakumar, and Wolpert (2014), referred to as the “power posterior”. We prefer this more general way as the localization data is not binary and we wanted to model the decision noise in the same way for both localization and detection datasets. While the power posterior is not the only reasonable way to formalize decision noise, Acerbi et al. (2014) showed more generally that this implementation can be used to approximate well the formation of decisions based on various types of noisy (Poisson-like or Weber-like) or sample-based posteriors.

Instead of reporting the 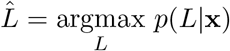, in this model we assume that the observer picks a sample from a noisy response distribution 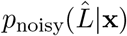 that is implemented via the “power posterior” computed as below:

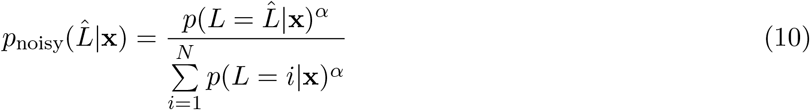

We see that setting the decision noise parameter *α* to 0 would give a uniform distribution, which corresponds to the observer responding randomly. Taking the limit *α* → ∞ would give back the MAP estimate, as in Model 1 above.

Lastly, to compute the model predictions conditioned on the stimuli **s**, 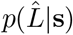, we used sampling as described for the previous model to estimate the integral:

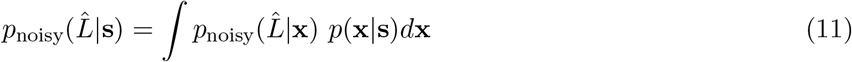

### Optimal-observer model for Detection

The optimal-observer model for detection is highly similar to the localization one and has been previously derived in (Ma et al., 2011; Mazyar et al., 2012) (also see (Ma, 2019) for a primer on Bayesian decision models). We briefly present it below.

#### Step 1. Encoding

The encoding stage is presented here as well in Figure 2B. For detection, the target present variable *C* can take values 0 or 1. If *C* = 1, *T_L_* = 1 and *T*_*i*≠*L*_ = 0. However, if *C* = 0, all *T_i_* = 0. The rest of the encoding stage is the same as in localization.

#### Step 2. Decision rule

For target detection, the optimal observer will also deterministically perform MAP estimation and report “target present” (*Ĉ* = 1) if the log posterior ratio *d* defined as below is greater than 0. We denote it with *d*, or the global decision variable:

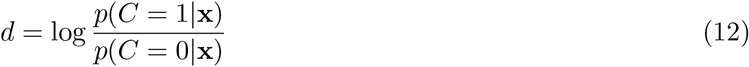

The observer might have a prior over the probability of target presence which is different than 0.5; alternatively the observer might have a higher motor cost for pressing a button with one hand as opposed to the other. Another possibility is that the observer might have an asymmetric utility for correct detection vs correct rejection. We incorporate all these possible contributions to bias into a single parameter, *p*(*C* = 1) = *p*_present_ and write *d* in terms of the prior and the log likelihood ratio:

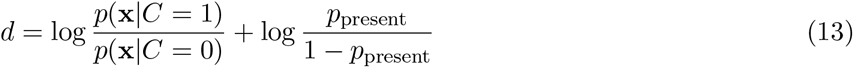

We unpack the likelihood ratio by marginalizing as follows and make use of the same properties as above, as well as of the fact that any location is equally likely among the *N* items:

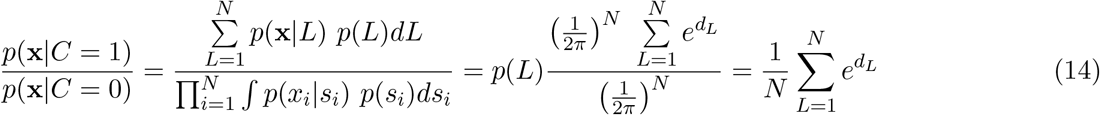

Thus, luckily, the global decision variable *d* can be written as a function of the local decision variables:

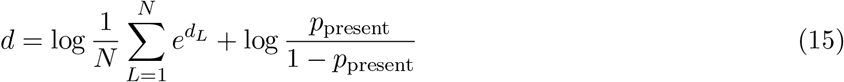

We assume the optimal observer reports their response *Ĉ* as follows:

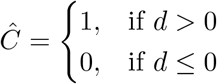

#### Step 3. Model predictions

Again, similarly as for localization, to eventually infer *Ĉ* knowing the stimuli the observer saw, we need to marginalize over all the possible measurements. Knowing *p*(*Ĉ*|**x**) from above, we compute the probability of response *p*(*Ĉ*|**s**) by estimating the integral below by sampling; specifically taking 1200 measurements calculated and averaging over the corresponding outcomes.

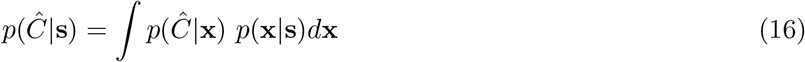

### Optimal-observer model for Detection : addition of decision noise, Model 2

In detection, we also formalize the decision noise parameter with *α*. We assume that observer picks a sample from a noisy response distribution *p*_noisy_(*Ĉ*|**x**) that is implemented with the power posterior. Instead of *N* values as in localization, the distribution of the power posterior can only take two values. Then the probability of choosing the response “target present” ends up following a logistic function of the decision variable *d*:

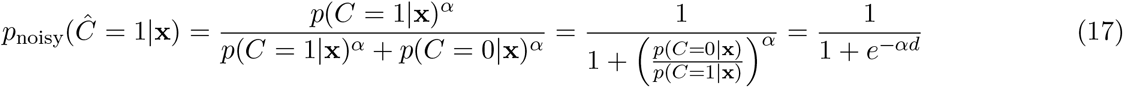

As in localization, a value of *α* → 0 corresponds to a flattening of the decision variable as if there was no information; the observer will respond close to chance. Taking the limit *α* → ∞ corresponds to a sharpening of the decision variable as if there was no noise, giving back the previous model.

The decision rule and the model prediction calculations are modified similarly to the way we showed for localization. The decision rule to respond “target present” or *Ĉ* = 1 vs “target absent” or *Ĉ* = 0 becomes:

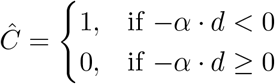

Lastly, we used *p*_noisy_(*Ĉ*|**x**) as defined above and sampling as described for localization to compute the model predictions:

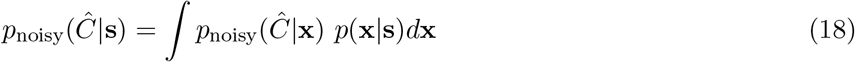

The generalization of the logistic function to several alternatives - the softmax function - has been widely used to model decision noise (Daw, O’Doherty, Dayan, Seymour, & Dolan, 2006). Other ways of formalizing the decision noise have been put forward: Shen and Ma (2019) formalized decision noise as Gaussian noise with standard deviation *σ_d_* to the global decision variable *d.* The logistic function and Gaussian noise actually represent very similar formalizations of decision noise as the logistic function resembles the Gaussian cumulative density function (cdf); we chose the logistic function to have the same implementation as in the localization task.

#### Model intuition

Figure 3 shows the decision landscapes for set size *N* = 2, for both localization (A) and detection (B), for three combinations of possible precision values associated with the two measurements (relative to the target).

In the case of localization (Figure 3A), the observer chooses the item with the highest local decision variable. If one item is encoded with higher precision, it will be chosen when its measurement is narrowly close to the target; the other item will be chosen the rest of the time. If the precisions associated with the 2 measurements are equal as in the third subplot, the decision landscape will present a bow-tie pattern and each item will be chosen half the time, based on which measurement is closer to the target. The bow-tie pattern takes this exact shape for any precision values as long as they are equal among the two items.

In the case of detection (Figure 3B), the higher the precision associated with a measurement, the more narrow the region around the target for which the observer reports “target present”. This makes sense in light of the model decision rules presented above, as the model assumes that the observers have knowledge of the precision associated with their measurements and factor them into the computations of the decision rules.

Figure 3 only shows the decision rules of the optimal-observer model and does not contain additional information about the generative model. We will see later how that informs the distributions of the measurements relative to the target and subsequently the response choices predicted by the optimal-observer model.

#### Model fitting

We performed maximum-likelihood estimation (MLE) of the parameters in the optimal-observer model variants. The parameters, denoted as *θ*, are as below:

**Table 1:**
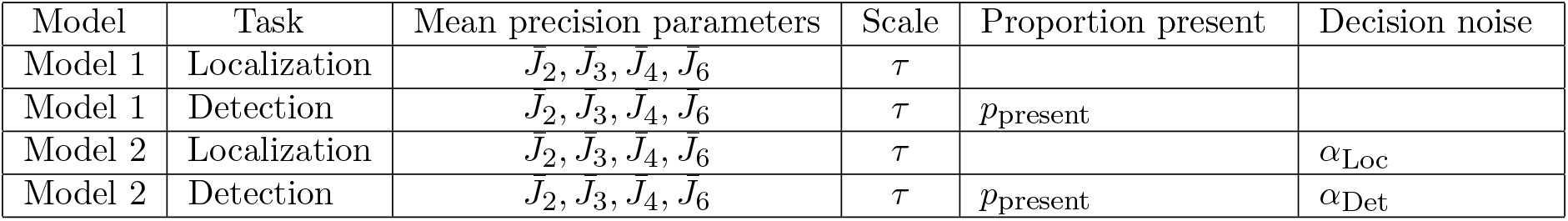
Models and their parameters for each task. Both model 1 and model 2 are optimal-observer models, but model 2 additionally includes a decision noise parameter.

For a particular model, the likelihood of a set of parameters *θ* is the probability of the data given those parameters, *p*(data|*θ*). We denote the log likelihood with LL. We assumed that trials are independent of each other and thus we could sum the log likelihoods across all trials:

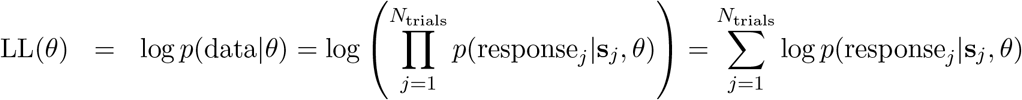

We denote the subject’s response on the *j*^th^ trial with response_*j*_ above. For the localization task this is the estimated location 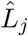 and for the detection task it is either “target present” or “target absent”, variable referred to as *Ĉ_j_* = 1 or respectively *Ĉ_j_* = 0.

We replaced extreme values of 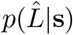 as follows: for 0 with 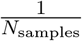 and for 1 with 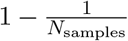. While in this case fixed sampling seemed to work well, it is worth considering computing the probability of response 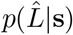 with a newly developed method, inverse binomial sampling (van Opheusden, Acerbi, & Ma, 2020).

Just as in “Model predictions”, we approximated the *p*(response_*j*_|*s_j_,θ*) through sampling. Even with 1200 samples, the log likelihood can be considered noisy and we have found standard deviations of around 0.3 across several function evaluations for a few parameter sets. To find the parameters θ that maximize LL(*θ*) we used an optimization method called Bayesian adaptive direct search (BADS) (Acerbi & Ma, 2017) that is especially suited for noisy functions. For each dataset and model, we ran BADS with 20 starting points and chose the best fitting parameters among those. Within BADS, we set the estimated noise size to 1 (`options.NoiseSize=1`). While we used 1200 samples and had a noise size below 1, Acerbi and Ma (2017) showed good performance for our localization data also with 800 samples and a standard deviation of the log LL(*θ*) of up to 3.5, updated accordingly in options.NoiseSize. Acerbi and Ma (2017) also showed that more widely used optimization functions such as Matlab’s `fmincon` or `fminsearch` are substantially worse at navigating noisy landscapes to find the global optima.

We searched the parameter space in the log range for almost all the variables to ensure higher proximity of values on each parameter dimension and thus an easier problem for the optimization algorithm. The plausible parameter ranges were [log(1.1), log(200)] for the mean of the gamma distribution 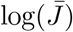 from which the precision *J* is drawn, [log(1.1), log(200)] for the scale parameter of the gamma distribution *τ*. For these parameters, we set the hard bounds equal to these plausible bounds. For *p*_present_, we set the plausible bounds to be [0.48, 0.63] while the hard bounds were [0, 1]. For the models with decision noise, the soft bounds were [log(0.001), log(100)], and the hard bounds were set equal to these soft bounds.

#### Model comparison

To assess whether the addition of the decision noise parameter in model 2 substantially improved the fit to the data after penalizing for the extra parameter, we performed model comparison based on the following commonly used metrics: the Akaike Information Criterion (AIC) (Akaike, 1974) and the Bayesian Information criterion (BIC) (Schwarz, 1978). These metrics are defined as AIC = −2LL* + 2*n*_pars_ and BIC = −2LL* + *n*_pars_ log*n*_trials_, respectively, where LL* is the maximum log likelihood and n_pars_ and nt_rials_ the number of free parameters and respectively the number of trials.

#### Statistical analyses

We analyzed accuracy as a function of set size with repeated-measures ANOVA. We quantified the summary statistics we defined by making use of the circular mean and circular variance functions from the CircStat toolbox (Berens, 2009). We used Spearman correlations to assess the correlations across these summary statistics. We performed logistic regressions to understand the observers’ responses (correct or incorrect) as a function of task metrics using Matlab’s `fitglm`. The output of Matlab’s `fitglm` also provided the regression model’s information criteria; we used AIC and BIC for the subsequent analyses. Prior to the logistic regression, all the predictors were Z-scored.

To identify whether adding task metrics to the logistic regression helps explain responses better, we used model comparison, in particular the strategy developed by Shen and Ma (2019), inspired by stepwise regression. We started by defining our *base model*, in which the only regressor was set size and the *full model*, in which added all the task metrics: min T-D difference, T-D mean and distractor variance. For each task metric/ regressor/ factor (F), we measured *factor usefulness* as the knock-in difference (KID) for both AIC and BIC as the difference between the respective metric of the base model and the one of the base model with the added factor: KID_AIC_(*F*) = AIC(Base) - AIC(Base+F) and same for BIC. For each task metric, we also measured *factor necessity* as the knock-out difference (KOD) for both AIC and BIC as the difference between the respective metric of the full model without the factor and the one of the full model: KOD_AIC_ = AIC(Full-F) - AIC(Full) and same for BIC. More positive KID values represent more evidence that the factor is useful; more positive KOD values represent more evidence that the factor is necessary.

## Results

The goal of Experiment 1 was to characterize in detail visual search with heterogeneous distractors by analyzing the effects of set size, task, and temporal order of target and distractor (perception and memory conditions). We analyzed all effects both in a model-free and in a model-based way. For the model-free approach, we characterized the data with psychometric curves for accuracy as a function of the minimum orientation difference between the target and the distractors, and respectively the circular mean of the distractors. Additionally, we calculated distractor circular variance within a trial. All our summary statistics are presented in Table 2. For the model-based approach, we applied and extended the general structure of the optimal-observer process model from (Ma et al., 2011; Mazyar et al., 2012) and tested its generality in capturing performance in visual search with heterogeneous distractors across more detailed metrics and experimental conditions.

**Table 2:** *N*-AFC target localization and target detection summary statistics. We used summary statistics A) - D) for both localization and detection, while E) was only applicable for localization. Additionally, for detection, we split the dependent variable, proportion correct, into hit rate and false-alarm rate.

|  | Dependent variable | Independent variable | Figures |
| --- | --- | --- | --- |
| A) | Proportion correct | set size ($N$) | 4A, A5A, A7A |
| B) | Proportion correct | the minimum absolute orientation difference from the target to the distractors, denoted by “min T-D difference” | 4B, 6B, 7, 8, 9 |
| C) | Proportion correct | the difference in orientation from the target to the circular mean of the distractors within a trial, denoted by “T-D mean” | 4C, 6C, A5C, A7C |
| D) | Proportion correct | distractor heterogeneity measured as the circular variance of the orientations of the distractors within a trial, denoted by “distractor variance” | 4D, 6D, A5D, A7D |
| E) | Frequency of occurrence | the rank of the absolute orientation difference between the chosen item and target (when the chosen item is the target itself, this rank is 0) | 4E, A5E |

### Summary statistics

We asked what influences performance in the target localization and target detection tasks. We defined a rich set of five summary statistics (Table 2). Summary statistics A - D overlap with the ones from Calder-Travis and Ma (2020). We characterized performance in localization according to all five summary statistics, and in detection according to A-D, as E was only applicable to localization. First, we focused on the localization data and model fits in isolation; next, we showed the detection results, followed by the joint fits to both.

We asked to what extent the summary statistics were correlated with each other. To address this, we present the distributions of each of these summary statistics and their correlations in Appendix B. We calculated the Spearman correlations by simulating 10000 trials for the localization condition (in which the target is always in the search array). When targets and distractors follow uniform distributions (Table A1A), we found positive correlations between min T-D difference and T-D mean and negative correlations between min T-D difference and distractor variance. These correlations tended to decrease in magnitude as set size increased.

For target and distractors distributions different than the uniform, we would see different patterns of these correlations. To illustrate this, Table A1B shows the case when the target and distractors follow a von Mises distribution with mean 0 and concentration parameter 1.5, as in Calder-Travis and Ma (2020). The Spearman correlations between min T-D difference and T-D mean are positive and only mildly decrease with increasing set sizes, while the min T-D difference and distractor variance are not significantly correlated.

### Target localization: data and optimal-observer model fits

We first explored the patterns in the data according to the summary statistics in Table 2. We saw that proportion correct decreases with set size *N* (Figure 4A). Speed accuracy trade-offs were not at play here as reaction times increased with set size (see Appendix H). ANOVA revealed significant effects for proportion correct with set size for both perception (*F*(3, 40) = 137.3, p = 4.36 · 10^−21^) and for memory (*F*(3, 40) = 146.2, *p* = 1.37 · 10^−21^). Since the distribution of the circular distances of the target to the most similar distractor (min T-D difference) varies with set size, we placed the bin values separately for each set size (Figure 4A), such that equal number of data points would go into each bin. Taking into account that stimuli are uniformly distributed, we analytically calculated the positions of these bins (see Appendix D).

**Figure 4:**
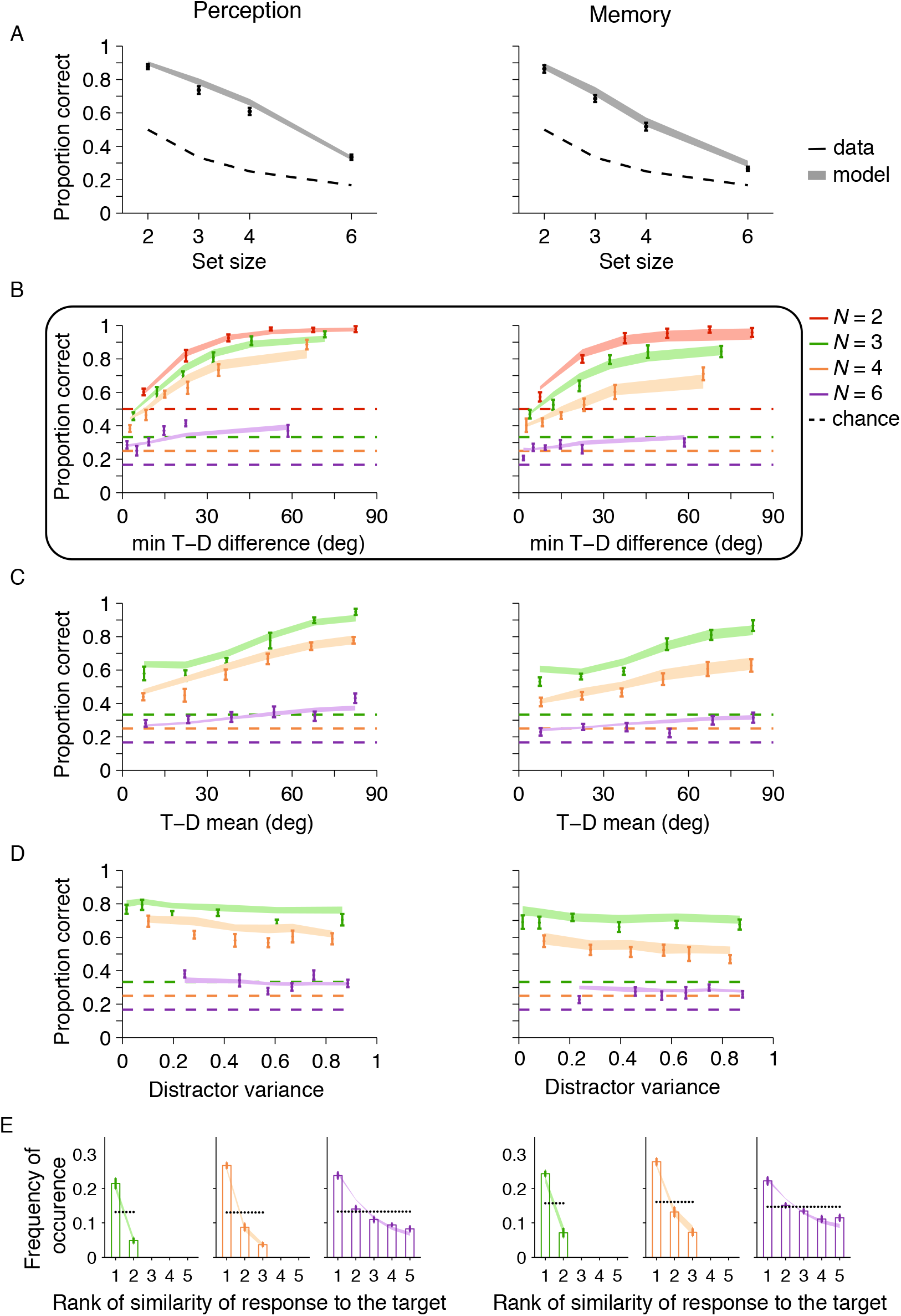
Results from the target localization condition in Experiment 1: data (mean ± sem and model 1 fits. Proportion correct decreases with set size and increases most saliently with min T-D difference. Here and in the upcoming figures, both perception (Left) and memory (Right) data were collected within the same participants. **(A)** Proportion correct with set size. **(B)** Proportion correct with the min T-D difference difference. The box indicates the summary statistic that is the most diagnostic of performance. **(C)** Proportion correct with the T-D mean. **(D)** Proportion correct as a function of circular variance of the distractors. **(E)** Frequency of occurrence of chosen (incorrect) item by rank of similarity to the target.

Visually, proportion correct increased with the min T-D difference (Figure 4B). Each one of the subplots in Figure 4B-D presents proportion correct split by set size *N* and the corresponding summary statistic: min T-D difference, T-D mean or distractor variance.

We applied several logistic regression models (Table A2) to evaluate the importance of each of these factors. Note that for these logistic regressions we only incorporated trials with set sizes greater or equal than 3, such that all summary statistics are meaningful. Regression model 1 represents the base model with just set size as the factor and regression model 5 represents the full model, with all 4 factors: set size, min T-D difference, T-D mean and distractor variance. Regression models 2, 3 and 4 are built as knock-in from the base model and are used to assess the usefulness of each factor. Models 6, 7 and 8 are built as knock-out from the full model and are used to assess the necessity of each factor. Positive KID shows factor usefulness and positive KOD signals factor necessity. A factor is more useful if it has a more positive KID and more necessary if it has a more positive KOD.

We found that the min T-D difference regressor was the most *useful* factor when added to the base model with the knock-in method as it improved the model criteria the most (Figure 5A). T-D mean was also useful, while distractor variance was not. With the knock-out method, there is no robust evidence that either regressor is *necessary*; knocking out any factor from the full model leads to decreases in information criteria and thus either negative KOD or inconsistent (Figure 5B). It is notable that the min T-D difference factor is necessary according to AIC, but not to BIC. The fact that neither factor is *necessary* is likely due to numerous possible trade-offs. Keep in mind that a knock-down of a factor from the full model also eliminates 4 interactions in addition to the factor itself. As presented in Table A1A, there are correlations among min T-D difference and T-D mean and distractor variance, so it is perhaps not surprising that a knock-down of this factor (or either factor) does not hurt the model fit. In addition to visual observation, these results further consolidate the status of set size and min T-D difference as the most diagnostic of performance.

**Figure 5:**
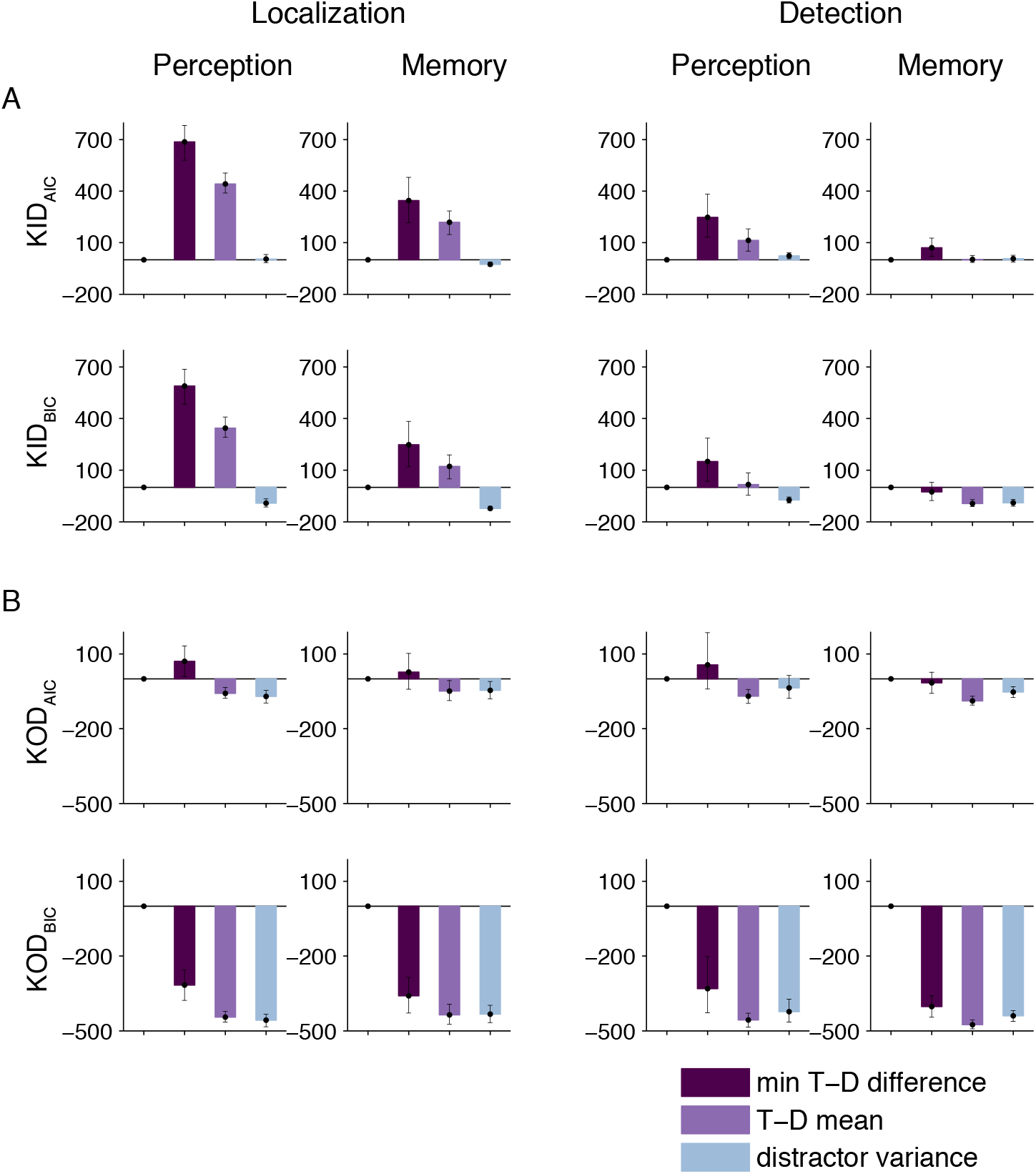
Experiment 1: Factor importance for the regressors: min T-D difference, T-D mean and distractor variance. min T-D difference was the most useful factor; T-D mean was useful as well, but distractor variance was not. Neither factor was necessary. **A)** Factor usefulness : sum across subjects and bootstrapped 95 % confidence intervals of the knock-in difference (KID), for both AIC (top) and BIC (bottom). As a reminder, KID_AIC_(*F*) = AIC(Base) - AIC(Base+F), F being the factor. The equation is analogous for BIC. **B)** Factor necessity: sum across subjects and bootstrapped confidence intervals of the knock-out difference (KOD) of the full model from the model without the factor, for both AIC (top) and BIC (bottom). As a reminder, KOD_AIC_ = AIC(Full-F) - AIC(Full).

Lastly, we unpacked the incorrect response choices. Specifically, we broke down the error responses by the rank of similarity to the target T. Proportion response decreased with the increase of the rank of (orientation) similarity to target (Figure 4E). Chi-squared goodness-of-fit tests confirmed that the distributions of the similarity rank of the response were different from uniform for every participant and every set size higher than 2.

While the logistic regression models employed above gave us a first sense of the factors that influence performance in localization with heterogeneous distractors, here our focus is also to employ an optimal-observer model to better understand the mechanisms of target localization in visual search.

As the optimal-observer model, we chose the variant model which usually provided the best fit in previous work (Mazyar et al., 2012, 2013). Therefore, we fit the optimal-observer model version with a variable-precision encoding stage and Bayesian decision rule. We overlayed the model 1 (without added noise) predictions for the best fitting parameters on the localization performance data and saw that it visually captured the data well (Figure 4). The optimal-observer model predictions increase with increasing min T-D difference and T-D mean, but seem flat with increasing distractor variance. Lastly, we emphasize that Figure 4E depicts the optimal-observer model fitting a fine grained break-down of the data: when observers were incorrect, their response choices tended to be the ones more similar to target.

We further explored whether model 2 with an additional decision noise parameter could capture the data better, but we found this was not the case according to model comparison metrics AIC and BIC (Figure A10).

### Target detection: data and optimal-observer model fits

Performance decreased with set size (Figure 6A) as follows: for perception, ANOVA revealed that hit rate varies with set size (*F*(3,40) = 2.66, *p* = 0.061) and so does false-alarm rate (*F*(3,40) = 9.88, *p* = 5.26o10^−5^) ; for memory, ANOVA revealed significant effects of hit rate with set size (*F*(3,40) = 3.7, *p* = 0.019) and similarly for false-alarm rate (*F*(3, 40) = 8.62, *p* = 1.55 o 10^−4^).

**Figure 6:**
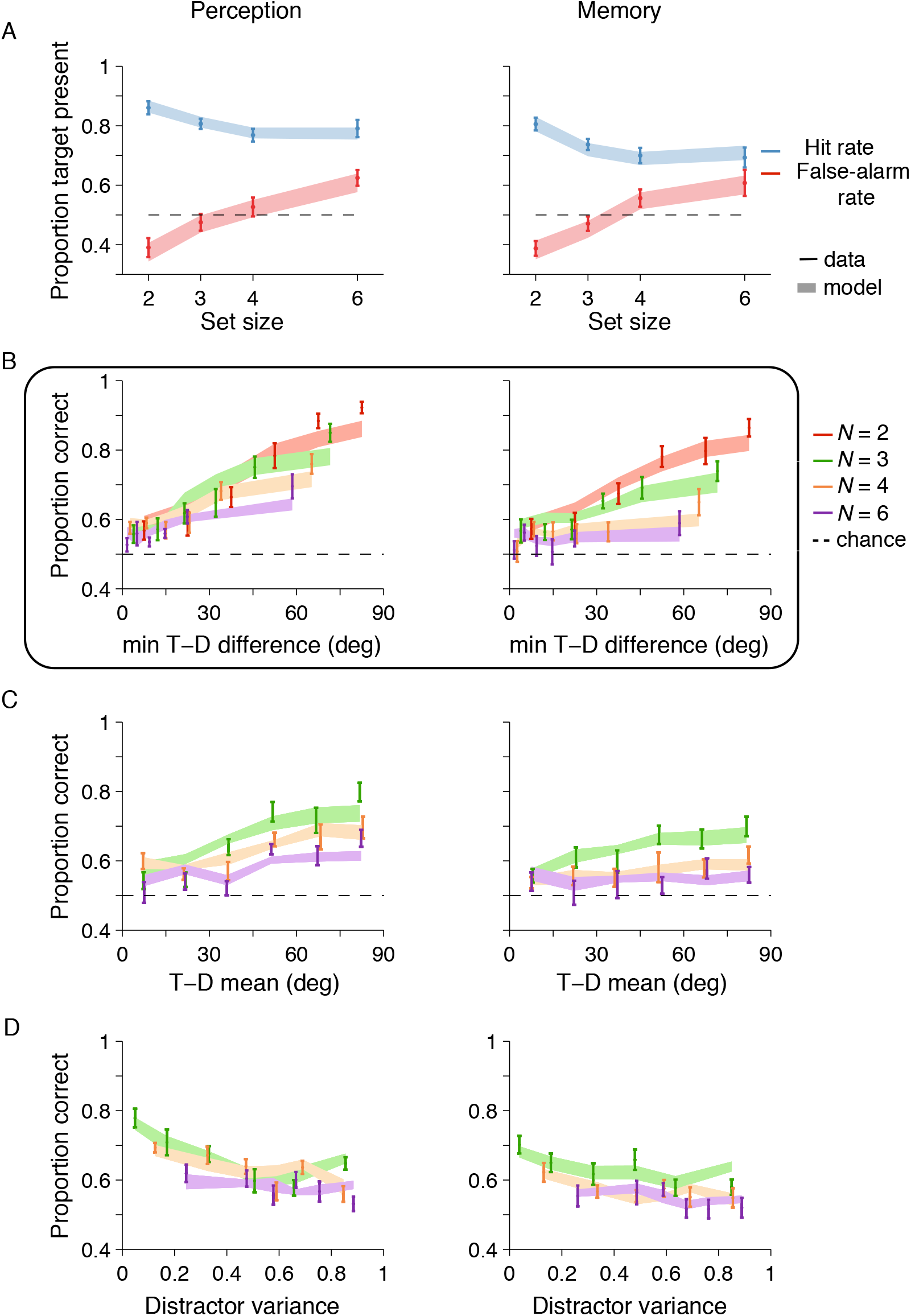
Results from the target detection condition in Experiment 1: data (mean ± sem) and model 1 fits. Proportion correct increases most saliently with min T-D difference. **(A)** Proportion correct as a function of set size. **(B)** Proportion correct as a function of the min T-D difference. As in Figure 4, here also we put a box around this summary statistic as it is the most diagnostic of performance and will be used in future plots. **(C)** Proportion correct with T-D mean. **(D)** Proportion correct as a function of the circular variance of the distractors.

As in localization, we performed logistic regressions analyses corresponding to each subplot (Figure 6B-D). We also found that the min T-D difference regressor was the most *useful* factor when added to the base model with the knock-in method as it improved the model criteria the most (Figure 5B) across the majority of conditions (with the exception of Detection-Memory, BIC metric). With the knock-out method, there is no robust evidence that either regressor is *necessary*; knocking out any factor from the full model lead to decreases in information criteria (Figure 5B).

As we said before, the optimal-observer model only differed from the localization one in the decision rule stage. We were also able to capture the detection data relatively well with the optimal-observer model (Figure 6), in line with Mazyar et al. (2012) and Calder-Travis and Ma (2020). These fits are satisfactory; the data was captured even better by including a decision noise parameter (Figure 7). For brevity, for this figure and for most of the paper from now on, we show proportion correct with the most diagnostic min T-D difference metric. We validated this visual observation that decision noise improves the optimal-observer model fits in Detection-Perception with the model comparison metrics AIC and BIC (Figure A10).

**Figure 7:**
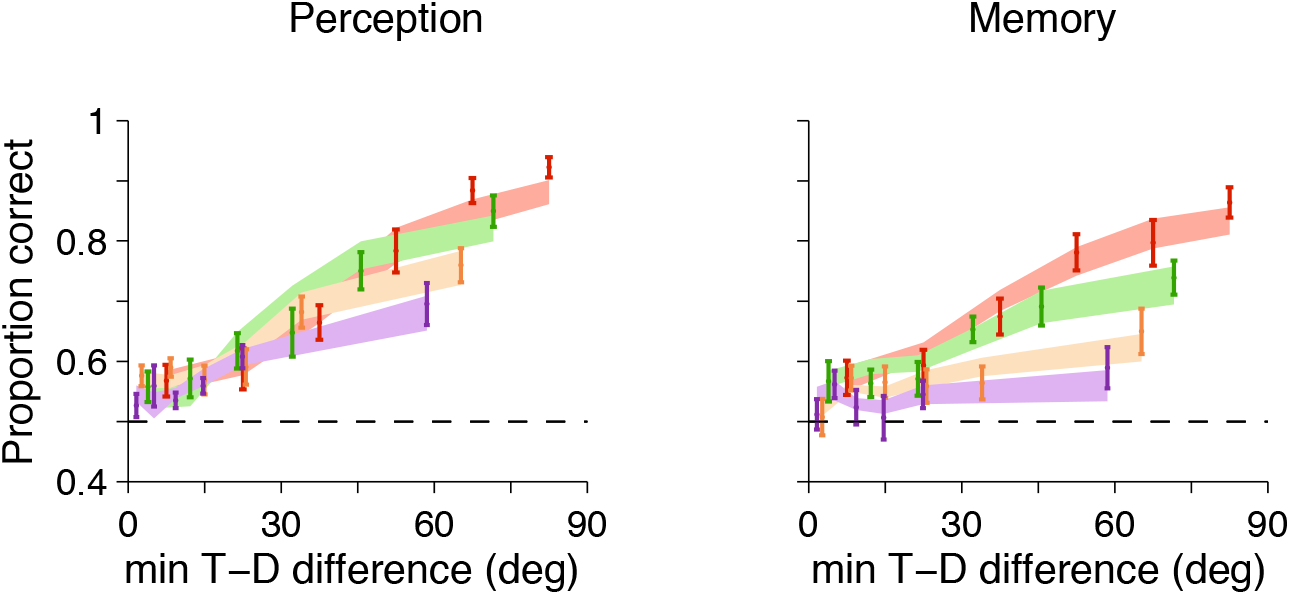
Results from the target detection condition in Experiment 1: part of the data (mean ± sem) and model 2 fits. Model 2 which includes decision noise provides a better fit to the detection data. Here and in the upcoming figures, we only show part of the data: performance with the most diagnostic summary statistic, min T-D difference. Notice that the addition of decision noise to model 2 makes these fits better than for model 1 (compare with Figure 6B, especially Perception). This visual observation is validated by model comparison (Figure A10).

### Target localization and target detection: joint fitting

Since the same observers performed both the detection and the localization tasks and the generative models are highly similar at the encoding stage, we can jointly fit the data from both tasks per observer. We did this separately for the perception and memory conditions. This approach might shed some new light to the question whether localization and detection are underlined by shared early encoding processes (Liu et al., 2003). In the joint fits depicted here, we specifically assumed that the mean precisions across set sizes and the scale parameter *τ* are shared across localization and detection, and that there are two decision noise parameters, *α*_loc_ and *α*_det_. A model in which we assume that the decision noise parameter was also shared across localization and detection provided a worse fit to the data. The joint model fits were able to capture the data well (Figure 8). While Figure 8 shows only performance with min T-D difference, the Appendix E shows performance with all the summary statistics (Figures A3 and A4). For Experiment 1, model 2 (with the addition of the decision noise parameters) did not provide a substantially better fit to the data, as confirmed with competitive model comparison (Figure A10).

**Figure 8:**
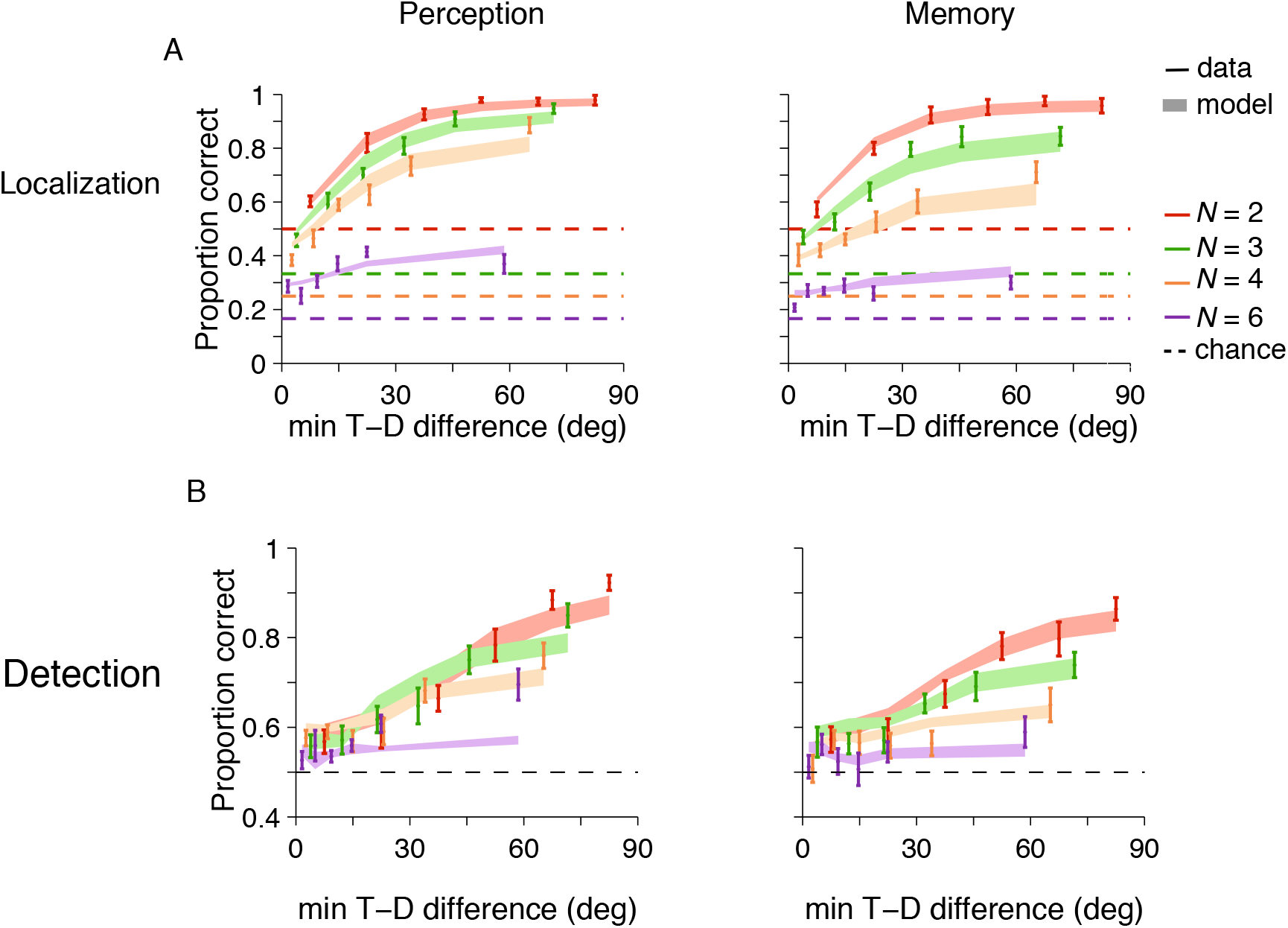
Jointly fitting all data of experiment 1: data (mean ± sem) and joint model 1 fits. The joint model fits were able to capture the data well. Results from **A)** localization and **B)** detection conditions with the optimal-observer model with shared precision parameters across the 2 tasks. Again, we only show here the most diagnostic summary statistic, min T-D difference. We see that localization joint model fits are comparably good with the ones in localization alone (compare with Figure 4B); however, in detection, we do see some deviations (compare with Figure 6B). For performance with respect to all the summary statistics, see Appendix E: Figure A3 for localization and Figure A4 for detection.

### Experiment 2: smaller stimulus spacing

The patterns of results we described for Experiment 1 largely hold for Experiment 2 as well. We see a strong effect on performance of min T-D difference, pattern which is captured by the optimal observer model well, both when fitted separately and jointly. We depict this in Figure 9 (see Appendix F, Figures A5 and A7 for full summary statistics and separate model fits for localization and respectively detection). Again, as in experiment 1, model 2 (with the addition of the decision noise parameter) was able to capture the data even better in Detection-Perception, but not in localization, and additionally in the Joint case (Figure A9). These visual observations were confirmed with competitive model comparison (Figure A10). The logistic regressions analyses for experiment 2 are largely in accordance with the ones from experiment 1 and are presented in full in Appendix F: the min T-D difference was the most useful factor in almost all conditions, though it was not necessary. We do note one exception, in the case of Experiment 2, the Detection-Memory condition, when distractor variance was actually the most useful factor (Figure A6).

**Figure 9:**
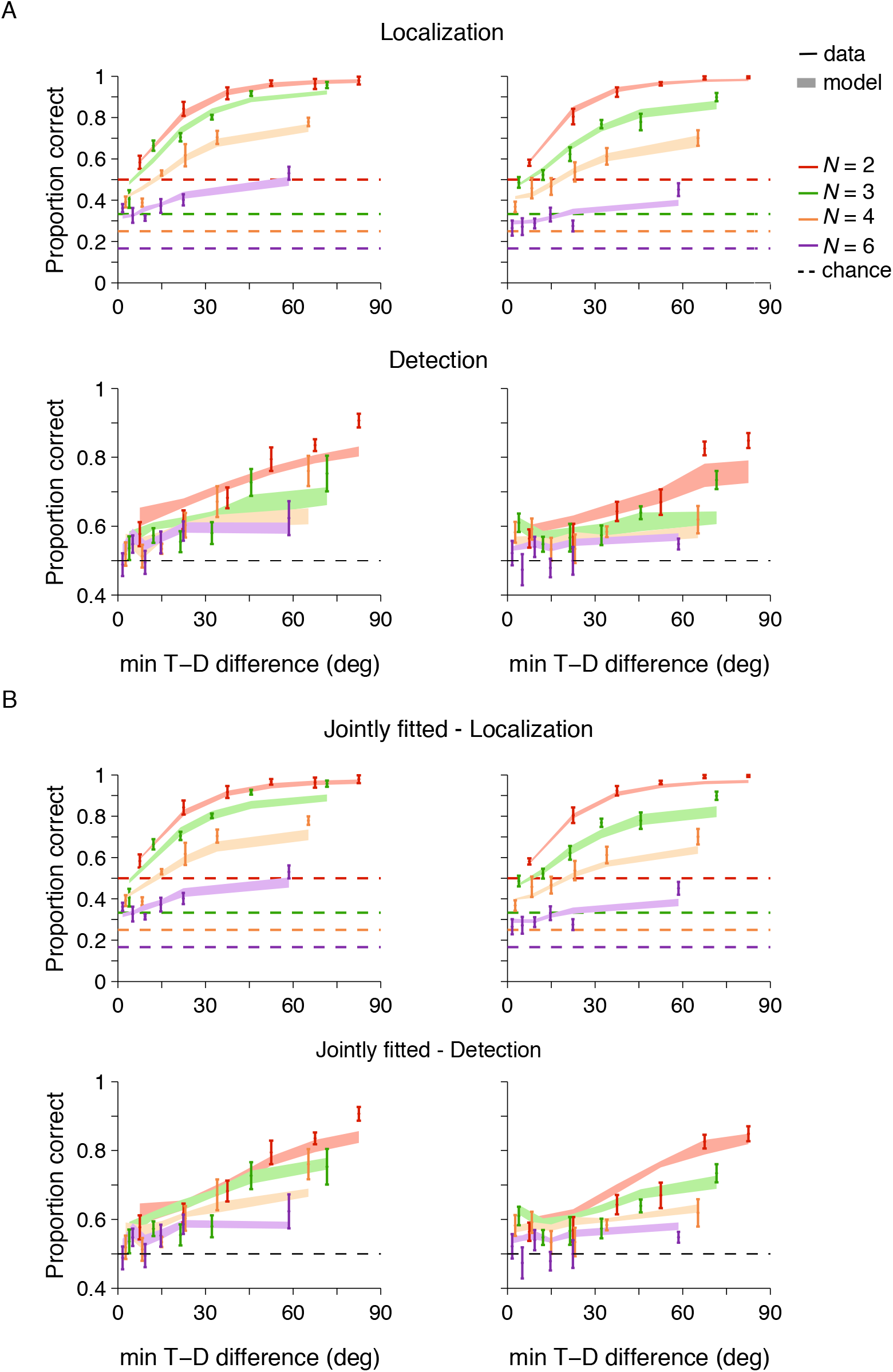
Results from the localization and detection conditions in Experiment 2: data (mean ± sem) and model 1 fits, both A) separately and B) jointly fitted. Experiment 2 replicates the main results from Experiment 1: proportion correct increases saliently with min T-D difference across both localization and detection and this pattern is captured well by the optimal-observer model. The separate model fits recapitulate the patterns seen in Experiment 1: the optimal-observer model for localization fits the data very well, while the model for detection fits the data relatively well. As is also the case for Experiment 1, the joint models fits are comparably good relative to the separate model fits.

### Min T-D difference and the optimal-observer model

Why would the min T-D difference be the most useful factor in predicting the observer’s responses? To see how the optimal-observer model provides insight into how the min T-D difference factor influences performance more than others, we revisit Figure 3 which depicts the decision rules for set size *N* = 2, for both localization and detection. We caution however that min T-D difference is calculated based on the stimuli, while the intuitions discussed below are based on the measurements of the stimuli.

The decision rules are presented in measurement space in Figure 3, but the distributions of measurements are not uniform. To better understand how the visualization of the decision rules informs the observer’s response choices, we make use of additional information from the generative model. The target and distractors are uniformly distributed, and each of them are assumed to be encoded by the observer into measurements with noise that follows a von Mises distribution. The target is always present in localization and thus there the distribution of measurements will be von Mises around the target for the target and uniform for the other distractor items. The target is present on half of the trials in detection; thus the distribution of measurements will be a mix of von Mises and uniform for the target and uniform for the other distractors.

For localization (Figure 3A), we get an intuitive sense of how min T-D difference influences performance. The observer chooses the item with the highest local decision variable. In the case of the third subplot with two measurements of equal precisions, the bow-tie pattern corresponds to picking the min T-D difference. For the first two subplots in (Figure 3A), the choice of the min T-D difference is distorted a bit, most saliently towards the center band. Even with a slightly higher proximity of measurement 1 to the target, the observer will still pick the item at location 2. As the variable precision model assumes that precision varies across trials and items according to a Gamma distribution with fitted parameters (see next section), large discrepancies between the precisions with which two items are encoded will be rare. Thus, it is likely that the center band in the decision variable landscapes across trials will not be very thick and thus the distortion from picking the item with min T-D difference not very frequent.

If we imagine extending this decision rule visualization to higher dimensions corresponding to search with several items, we can additionally imagine the influence of distractor variance. At a first intuition, higher distractor variance should not have an effect on performance: the observer should still just pick the item with the highest decision variable. However, with higher distractor variance, there will be a higher chance that the measurement of one of the distractors would fall closer to the target (or comparatively close to the measurement of the target itself as in the first two subplots) and thus be reported instead. This explanation of the influence of distractor variance through min T-D difference has been put forward in the standard SDT accounts.

For detection, Figure 3B shows that the higher the precision associated with a measurement, the more narrow the region around the target for which the observer reports “target present”. Intuitively, we can see how the min T-D difference influences performance: if a distractor is close to the target, chances are its measurement will also be and fall within the white arms of the cross-shape. However, given that the “target present” response regions are with respect to all axes (here 2, as set size is 2), we can see how with more variable distractors, by chance, the min T-D difference of one might lead to a measurement - target value small enough for the observer to report “target present”.

### Model parameters

Lastly, we present and analyze the optimal-observer model parameters. Figure 10 presents the parameters from model 1 for both experiments: the mean precisions log 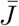 and the scale parameter *τ* that capture the Gamma distributions describing how encoding precisions vary across trials and items.

**Figure 10:**
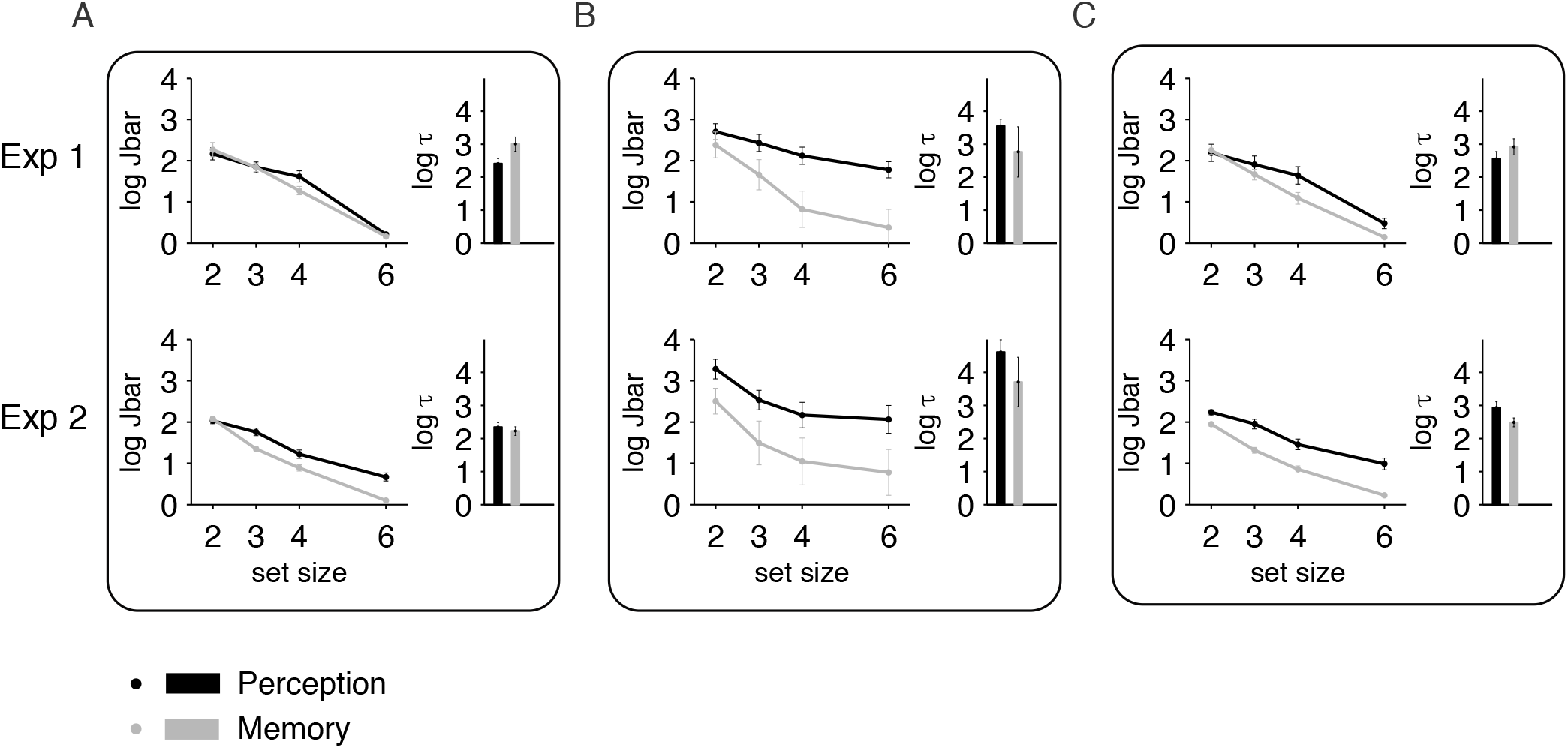
Estimates of precision parameters in model 1 across conditions and experiments: mean precision decreases with set size and is higher in perception than in memory. **(A)** Localization. **(B)** Detection. **(C)** Jointly fitted localization and detection.

Mean precision log 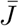 seems to decrease with set size across all experiments and conditions (Figure 10). In Experiment 1 Localization-Perception, ANOVA revealed significant effects with set size (*F*(3,40) = 46.3, *p* = 4.38 · 10^−13^), similarly in detection (*F*(3, 40) = 3.56, *p* = 0.023) and in the joint model fitting condition (*F*(3, 40) = 13.99, *p* = 2.21 · 10^−6^). In memory, ANOVA again revealed significant effects of log 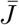 with set size for localization (*F*(3, 40) = 58.78, *p* = 1.02 · 10^−14^), for detection (*F*(3, 40) = 4.66, *p* = 0.0069) and for the joint model precision parameters (*F*(3, 40) = 60.53, *p* = 6.31 · 10^−15^).

In experiment 2, ANOVA revealed that log 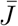 varies significantly with set size in Localization-Perception (*F*(3, 20) = 19.82, *p* = 3.31 · 10^−6^), but not in Detection-Perception (*F*(3, 20) = 1.71, *p* = 0.19) and again for the joint condition (*F*(3, 20) = 10.35, *p* = 0.0003). In memory, we found again that log 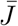 varies with set size for localization (*F*(3, 20) = 108.27, *p* = 1.55 · 10^−12^), but not for detection (*F*(3, 20) = 1.03, *p* = 0.40), and again for the joint fits (*F*(3, 20) = 60.98, *p* = 3.05 · 10^−10^).

Visually, the mean precision parameter 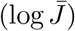 seems higher in perception that in memory. To check whether we could meaningfully compare the mean precision parameter values statistically, we first checked for differences in the *τ* parameters across perception and memory (mean precision and *τ* are known to trade-off in the variable precision model). Paired sample t-tests were used to determine whether there were statistically significant differences between the *τ* parameter estimates across perception and memory for each condition depicted in Figure 10. We found no statistical differences between the *τ* parameters in Perception vs Memory for almost all conditions (all *p* > 0.082), except for Experiment 1 Localization.

Next, we performed two-way repeated measures ANOVA on log 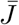 with set size and perception vs memory for the conditions for which *τ* did not differ across perception and memory. For Experiment 1 Detection, we find a significant effect of set size (*F*(3,80) = 7.69, *p* = 0.0001), a significant effect of perception vs memory, (*F*(1, 80) = 16.7, *p* = 0.0001), and no significant interaction (*F*(3, 80) = 1.17, *p* = 0.32). For Experiment 1 Joint, we find a significant effect of set size (*F*(3,80) = 49.8, *p* = 2.84 · 10^−18^), a significant effect of perception vs memory, (*F*(1, 80) = 5.35, *p* = 0.023), and no significant interaction (*F*(3, 80) = 1.22, *p* = 0.31). For Experiment 2 Localization, we find a significant effect of set size (*F*(3,80) = 82.23, *p* = 3.74 · 10^−17^), a significant effect of perception vs memory, (*F*(1, 80) = 16.35, *p* = 0.00020), and no significant interaction (*F*(3, 80) = 2.71, *p* = 0.057). For Experiment 2 Detection, we find no significant effect of set size (*F*(3, 80) = 2.33, *p* = 0.088), a significant effect of perception vs memory, (*F*(1, 80) = 6.06, p = 0.018), and no significant interaction (*F*(3, 80) = 0.060, *p* = 0.98). For Experiment 2 Joint, we find a significant effect of set size (*F*(3, 80) = 42.7, *p* = 1.49 · 10^−12^), a significant effect of perception vs memory, (*F*(1, 80) = 34.4, *p* = 7.32 · 10^−7^), and no significant interaction (*F*(3, 80) = 1.06, *p* = 0.38).

Overall, these results help address more thoroughly our fourth goal of comparing visual search in perception vs memory. So far, we saw similarities across search in perception and memory, as our summary statistics had similar influences on search in perception versus memory (Figures 4 and 6). Figure 10 and the above statistics emphasize a difference, as precision in visual search varies with set size and with the perception vs memory condition, mean precision being higher in the perception condition. This is as intuitively expected and in line with Mazyar et al. (2012), study which also measured encoding precision according to this model. More broadly, this result joins previous studies which found reduced accuracy in visual search in memory relative to perception (Kuo et al., 2009).

In the Appendix, we also present the precision parameters from model 2 (Figure A11), as well as the decision noise parameters from model 2 (Figure A12). Where applicable, the trends are largely similar: precision decreases with set size across all conditions and tends to be higher in perception than in memory. However, the presence additional parameters (*τ* as before but also decision noise) decreases the interpretability of the precision parameters from model 2 as they might have larger trade-offs with the other parameters.

The observers who performed experiment 2 were different than the ones who performed experiment 1. Thus, we were not able to directly address the question of how the resolutions of perceptual versus memory representations differ across stimulus spacing conditions. Still, that experiment 2 produced results highly similar to experiment 1 is consistent with the study of Tamber-Rosenau, Fintzi, and Marois (2015), which showed that within Bouma’s limit, perception and memory are impaired at comparable extents with reducing stimuli spacing, albeit through different mechanisms. Relatedly but reaching a different conclusion, Ahmad et al. (2017) found that a measure of working memory precision was significantly lower for simultaneously presented near items relative to far items. Pertzov and Husain (2013) found a similar pattern, while Harrison and Bays (2018) found that the precision of working memory is independent of the cortical spacing between items. If the same observers would have performed both experiments 1 and 2, we could have addressed this question with a direct comparison between our mean precision parameters across experiments.

## Discussion

Visual search is frequent in the daily life of agents (people, animals and artificial intelligence agents), who have to localize and detect target objects among various other objects as steps towards actions and goals. We revisit the question of how target-distractor similarity and distractor heterogeneity influence performance, as the previous literature has not come to definitive conclusions based on unambiguous quantifications of these measures. Duncan and Humphreys (1989); Duncan (1989)‘s account of how stimulus similarity influences visual search performance allowed researchers to go beyond thinking in terms of parallel and serial search and provided an essential jump-start to the study of visual search with heterogeneous distractors. However, quantifying stimulus similarity with words such as “high” or “high enough” or “sufficiently distinct” or “neighbors” inherently reaches a limit in explanatory power.

Thus, here we studied visual search with heterogeneous distractors with simple parametric stimuli within a quantitative modeling framework based on SDT, in order to get more insight into the component processes. Only a handful of studies so far have applied STD or related models to visual search with heterogeneous distractors (Rosenholtz, 2001; Mazyar et al., 2012; Bhardwaj, van den Berg, Ma, & Josić, 2016; Calder-Travis & Ma, 2020). Moreover, we used two tasks, target detection as in previous work, but also target localization, in which observers reported the spatial location of a target among several alternatives. Using two tasks allows for a more in depth study of visual search in two different ways: first it permits to account for a larger range of visual search behaviors and second it provides a stronger test of the optimal-observer models. Additionally, as the second task was *N*-AFC target localization, we were likely able to provide a more ecologically relevant testbed for visual search theories and models (Dukewich & Klein, 2009).

Our first goal was to comprehensively assess what properties of the visual search array influence performance in visual search with heterogeneous distractors. Our second goal was to employ a normative model to better understand visual search. Optimal-observer models have been able to capture visual search datasets before (Ma et al., 2011; Mazyar et al., 2012; Calder-Travis & Ma, 2020); here we additionally attempted to subject this model to a more robust test, partly through employing the task of *N*-AFC target localization. Our third goal was to revisit within this optimal-observer modeling framework a question from more qualitative work in visual search: the relation between search behavior in target localization versus target detection. Our fourth goal was to compare differences between perception-based and memory-based visual search. We will now discuss our findings for each individual goal.

### Goal 1: Effect of distractor statistics on visual search performance

We first analyzed how performance is affected by the summary statistics describing the set of distractors on each trial. We found that performance was most strongly influenced by the most similar distractor (min T-D difference, Figures 4B, 6B, and Figure 5 for Experiment 1 and Figure 9 for Experiment 2). This is in line with the first part of the Duncan and Humphreys (1989) proposal: performance indeed decreases as target-distractor similarity increases, pattern widely replicated in other studies. It is important to note that target-distractor similarity can be quantified in several ways; here we used two such ways, min T-D difference and the mean of the distractors, T-D mean (these two measures being correlated with each other). As we used simple stimuli which varied parametrically and could be to quantified by one number, we had an unambiguous quantification of both target-distractor similarity and distractor heterogeneity, which was not the case in prior work (i.e, Duncan and Humphreys (1989); Duncan (1989) used letters and respectively a set of 6 colors).

We did not find evidence that performance decreased when distractor heterogeneity increased, which is in contrast to the second part of the Duncan and Humphreys (1989) proposal. Duncan and Humphreys (1989) and Duncan (1989) indeed emphasized that when targets are sufficiently distinct from non-targets, search is always very easy, and distractor heterogeneity should not have an effect. This is visually depicted in the asymmetry of their proposed “search surface”. However, this claim allows room for ambiguity, as it implies that some arbitrary criterion will need to be defined to determine when targets are “sufficiently distinct” from non-targets. Here, we considered a range of min T-D differences between 0 and 90o and found a more general pattern, as distractor heterogeneity was not a useful factor to explain performance, with the exception of Experiment 2, the Detection-Memory condition (Figure A6). Overall, with the logistic regression models and comparisons based on the factor knock-in and knock-out, we find that min T-D difference was the most *useful* regressor to explain performance, though there was no evidence that it was *necessary*.

The analyses described above are based on logistic regression models capture trends in the data. However, they provide insufficient insight into mechanism; for instance, they cannot address thoroughly questions such as: why is min T-D difference a useful factor for predicting performance while distractor variance is not? By contrast, the optimal-observer model breaks down the brain computations involved in visual search into two stages: an encoding stage of the search array into measurements in the brain and a sequential decision stage. The decrease in performance with increased min T-D difference is predicted by the optimal-observer model. In this model, with uniformly distributed targets and distractors there should be no additional decreases in performance with increasing distractor variance. Any effects that may seem driven by distractor heterogeneity are secondary to the influence of the min T-D difference. This is visible in the approximately flat model predictions in Figures 4D, 6D, and can be understood based on the decision landscapes from Figure 3 and their interpretation in relation to the min T-D difference.

A different pattern of the influence of distractor heterogeneity on visual search performance comes from Bhardwaj et al. (2016). They utilized a target detection task similar to ours and found a decrease in proportion correct with the sample standard deviation of orientations. They found that performance decreased with increasing distractor heterogeneity (Bhardwaj et al., 2016). Inconsistencies between their study and ours could be due to the differences in the distractor distributions employed. It is important to note that their distractor distribution was narrower and that their circular distractor standard deviation values were largely between 0 and 15o. Our corresponding circular standard deviations would go from 0 up to close to 90o, as the distractors follow a uniform distribution (though we report instead distractor variance, which is between 0 and 1). Additionally, they utilized a vertical target always the same across trials (like in the original Duncan and Humphreys (1989); Duncan (1989) experiments), while our target varied randomly across trials. Differences between tasks with the same target across trials vs variable targets can be reasonably expected (for instance, Treisman and Gormican (1988) found that search times were longer with variable targets than when targets were the same across trials).

A pattern similar to ours of how distractor variance influences performance in visual search comes from the recent study of Calder-Travis and Ma (2020). They investigated the effect of the same distractor statistics as ours on target detection performance with heterogeneous distractors across two different distributions of distractors: uniform and concentrated von Mises. In line with our results, they also found that min T-D difference and set size were the regressors that best predicted accuracy. They also explained this pattern with the optimal-observer model. Their decision rule landscapes depict the same pattern as ours in the uniform case, however in the concentrated case a different decision landscape emerged. Unlike in the case of uniform distractors, in the case of concentrated distractors they found that false alarms decreased as distractor variance increased. The shape of the decision landscapes depict how increasing min T-D difference and increasing distractor variance provide overlapping contributions to performance (cross-shape), while this is less so the case for concentrated distractors (diamond shape). Overall, their main result is in line with ours : min T-D difference is the main distractor statistic that influences performance in the case of uniform stimuli, without substantial contribution from distractor variance.

Some other work has used the term distractor heterogeneity to refer to variability of the distractor set across trials, but here we confined our discussion to this term as meaning variability within a trial. As such, here we did not consider distractor displays that are homogeneous (within a trial) but that vary across trials (Bravo & Nakayama, 1992; Nagy et al., 2005; Shen & Ma, 2016; Mazyar et al., 2013). We also did not consider situations in which the distractor distribution itself varied across blocks, as was the case in (Mazyar et al., 2013).

### Goal 2: Optimal-observer modeling of visual search with heterogeneous distractors

While there have been extensive applications of signal detection theory models to visual search, only a small proportion of this work used heterogeneous distractors (Rosenholtz, 2001). One way in which optimal-observer models differ from signal detection theory models is that they assume a unique Bayesian decision rule for the combination of information across items. There is some robust evidence that optimal-observer models can capture to a good extent visual search behavior with heterogeneous distractors in target detection tasks (Ma et al., 2011; Mazyar et al., 2012; Calder-Travis & Ma, 2020). As in Calder-Travis and Ma (2020), our comprehensive set of summary statistics constitute an even more robust test of optimality in visual search. Additionally, by using two search tasks, detection and localization, and the additional summary statistic from localization, we subjected the optimal-observer model to a more general and thorough test.

Specifically, we found that the optimal-observer model (Ma et al., 2011) was able to capture behavior in target localization (Figure 4). Crucially, the optimal-observer model also captured well the detailed breakdown of error types afforded by *N*-AFC target localization. The model also fit well the detection data (Figure 6); thus we recapitulated the patterns from Mazyar et al. (2012) and Calder-Travis and Ma (2020). Moreover, we also explored the addition of a decision noise parameter, which improved the optimal-observer model fits only in Detection-Perception (Figure 7).

We found that precision decreased with set size for both localization and detection, as well as for their joint fits (Figure 10). As emphasized by Mazyar et al. (2012), this is not an obvious result, as some pattern of decrease in performance with set size could have been accomplished with mean precisions that are constant with set size; the decrease in performance would have been due to the fact that the more distractors there are, the higher the chance that one of them would be confused with the target, as in the signal detection theory account (Palmer, 1995).

Overall, the optimal-observer models tested here captured the data relatively well for both localization and detection in visual search, both within perception and memory.

#### Sources of deviations from optimality: decision noise

We further discuss the possibility that the deviations of the optimal model from the data could come from suboptimalities in inference, in the form of decision noise. Decision noise could be simply what its name suggests - noise on the decision variable in the form that we assumed -, but could also account for a broad range of computational imperfections and erroneous inference (Beck, Ma, Pitkow, Latham, & Pouget, 2012). These include wrong beliefs, for instance about their own sensory noise used in the computation of the decision rule, or different decision rules or even a different model than assumed (model mismatch).

We found no benefit of an additional decision noise parameter to capturing the localization dataset (Figure A10). However, the model fits to the detection dataset were substantially improved by allowing for the decision noise parameter (Figure A10). Our finding of substantial decision noise in target detection is in contrast to the study of Shen and Ma (2019), which found little improvements in the model fits by including a decision noise parameter. A possible explanation for this discrepancy might come from an experimental difference: the target there was always vertical (Experiment 11 in Shen and Ma (2019), data originally from Experiment 1 in Mazyar et al. (2013)).

Is this distinction between the role of decision noise in localization (none) and detection (substantial) a real pattern, or an artifact of differential parameter trade-offs? Stengård and van den Berg (2019) and others cautioned that optimal models, due to their flexibility, could allow for decision noise or other suboptimalities in inference to masquerade as estimates of high sensory noise parameters. Relatedly, Shen and Ma (2019) found strong evidence for variable precision in perception, but this was no longer the case once they included the oblique effect and decision noise. As we only tested an optimal-observer model with variable precision, we cannot rule out the possibility that variable precision absorbs behavioral variability which could be decision noise in localization. Nevertheless, in the case of detection, decision noise provided an improvement in the model fit beyond what variability in precision accounted for. Thus, we believe that decision noise in detection captures a distinct imperfection in inference, separate from high or variable sensory noise.

Overall, within our study it is puzzling why within the same observers decision noise would impair the computation of the target detection decision, but not the maximum operation entailed in localizing the target. A simple explanation might be that observers might have experienced higher task engagement and thus less decision noise during the more nuanced decisions required by the *N*-AFC localization task. It is also possible that the more downstream decision processes in localization and detection could be subjected to different amounts of noise, in line with their possible segregation into the dorsal and respectively ventral streams. Such a segregation might account for the influence of decision noise in detection but not localization.

### Goal 3: Possible shared encoding processes in localization and detection

To test the possibility of shared encoding processes in localization and detection, we made use of our optimal-observer modeling framework, under which localization and detection are not fundamentally different tasks, but have a largely shared structure, and they only differ in the last decision stage.

Since the same observers performed both sets of tasks, we were able to see that fitting a model jointly to localization and detection with shared parameters was able to account relatively well for the data (Figure 8). As the optimal-observer model with shared parameters fitted jointly to localization and detection was able to capture the data well, we find it plausible that localization and detection might be reliant on similar early encoding processes. Our work connects to some extent to joint investigations of detection and localization from visual search with homogeneous distractors. Cameron et al. (2004) showed that signal detection theory can explain performance as a function of set size for identification, detection and localization. In our data, while the joint detection and localization fits could be better, the fact that they capture the data as well as they do seems to add further weight to the assumption of shared precision in the early visual encoding processes in target detection and target localization.

### Goal 4: Visual search in perception vs memory

We found that patterns from perception-based visual search such as the set size effect and the target-nontarget similarity effect (here min T-D difference) were also found in memory-based search (Figures 4A,B and 6A,B). These results are in line with prior work detailing such similarities across visual search in perception and memory (Kuo et al., 2009; Kong & Fougnie, 2019). All other distractor statistics had qualitatively similar influences on performance in both perception and memory-based search (Figures 4C,D,E and 6C,D). Differences across perception and memory arose as accuracy as well as the mean precision parameters were higher in perception than in memory (Figure 10), also in line with prior work (Kuo et al., 2009; Mazyar et al., 2012).

In the optimal-observer models we used, the precision parameters correspond to a quantification of the amount of sensory noise with which items are encoded. Such precision parameters are thought to have a neural correspondent in the amplitude or gain of a neural population representing the stimuli (Ma, Beck, Latham, & Pouget, 2006). As could be intuitively expected, as well as in line with Mazyar et al. (2012), mean precision decreased with set size and was higher in the perception condition relative to the memory condition. Thus, we conclude alongside the aforementioned studies that searching the external world in perception works through similar mechanisms as searching within one’s internal, memorized representation of the world.

### Replication with reduced stimulus spacing

We largely replicated these patterns of results in Experiment 2, with some minor differences, overall supporting the answers to our 4 goals presented above. The reduced stimulus spacing employed in Experiment 2 was meant to bring the stimuli together while keeping them still wide apart such that they are outside Bouma’s limit for crowding (Bouma, 1970). In this way, the assumption of the model that the stimuli from the search array are encoded independently of each other should still be justified (Ma et al., 2011); indeed we found that the optimal-observer still captured the data well.

Even apart from crowding, we considered the possibility that reducing the stimulus spacing might impair performance, especially so in the localization task. For instance, observers might make “swap” errors, which entail reporting the location of another item in the array (Bays (2016), but see Pratte (2018)). If swaps would have happened, they would have affected performance in localization but not in detection. However, even our reduced stimulus spacing condition kept the stimuli at 30 dva, sufficiently wide apart such that a hypothetical proportion of swaps would have been low. In theory, the decision noise parameter could have accounted for this additional source of errors, but the model with decision noise did not improve the fit substantially in localization in neither experiment.

### Limitations

While we compiled a rich set of summary statistics that allowed us to analyze how performance varies as a function of each statistic separately, we did not have enough trials to split performance across combinations of summary statistics. A finer split might have been able to address more directly whether heterogeneity made the distractors more or less similar to the target and would have constituted a more robust test of the predictions of signal-detection theory for visual search with heterogeneous distractors (Rosenholtz, 2001; Vincent et al., 2009).

#### Alternative models?

Another limitation of our study comes from the fact that we did not test alternative models besides the optimal-observer model. As such, we cannot make claims that the optimal-observer model captures the data better than other models, for instance with heuristic decision rules (as investigated in Calder-Travis and Ma (2020); Ma et al. (2015); Shen and Ma (2016)). We cannot claim that optimality is preferred by observers or even distinguishable from other models. As the optimal-observer model provided a good fit, we did not find a strong incentive to test alternative models. Optimal-observer models have been usually compared competitively against heuristic models with simpler rules, most prominently the max rule (Palmer et al., 1993; Ma et al., 2011). Beyond signal-detection theory and optimal-observer models, a few alternative models of visual search based on summary statistics (i.e, saliency or cover difficulty, inspired by computer science methods) have been shown to account for how behavior varies with distractor heterogeneity (Rosenholtz, 1999; Avraham, Yeshurun, & Lindenbaum, 2008). We did not delve into these alternative models either. Overall, as the optimal-observer models provided relatively good fits, we did not find compelling reasons against the hypothesis that human behavior in visual search can be captured well with an optimal-observer framework.

#### Towards naturalistic search?

We studied visual search with unidimensional oriented patches stimuli; by allowing the distractors to differ within a display, we examined a more naturalistic behavior than in the traditional SDT displays with homogeneous distractors. However, our experiment is still far from real-world search. On the one hand, visual search in real life is in some ways harder, as objects in the real world do not just vary on one dimension, but tend to have multiple features, values (Biggs & Mitroff, 2014) or affordances for actions (Wulff, Stainton, & Rotshtein, 2016) and are not always clearly differentiated from the background (Wolfe, 1994). On the other hand, visual search in the real world is easier, as rather than having to search through objects that are maximally unpredictable and briefly presented, in daily life we can exploit regularities, for instance by making use of templates (Geng, DiQuattro, and Helm (2017); Geng and Witkowski (2019)) or anchor objects (Boettcher, Draschkow, Dienhart, & Võ, 2018), and we can actively direct our eye movements to specific locations in natural scenes towards maximizing the information gathered (Najemnik & Geisler, 2005; Yang, Lengyel, & Wolpert, 2016; Rothkegel, Schutt, Trukenbrod, Wichmann, & Engbert, 2019). Here, we have have incrementally increased the ecological relevance of our visual search task. Challenges for future research entail incorporating even more of this complexity in an efficient way in an experimental design and developing suitable optimal-observer models for naturalistic tasks (for more information on these challenges, see Geisler and Ringach (2009); Geisler (2011)).

### Concluding remarks

Bayesian modelers of behavior have been criticized for making their models overly flexible and not always comparing them against alternative models (Jones & Love, 2011; Bowers & Davis, 2012). Indeed, when such comparisons are made, deviations from optimality frequently come to light (Bowers & Davis, 2012; Rahnev & Denison, 2018). The literature of visual search with a psychophysical methodology (Palmer et al. (2000), i.e. with eye fixation, brief presentation, and a small number of single-feature stimuli) offers an interesting counterpoint to this criticism. In this domain, Bayesian models are typically strictly constrained by the generative model of the stimuli, combined with a rather generic assumption about the form of measurement noise. In addition, these models are often being formally compared against alternatives (e.g. Ma et al. (2011); Mazyar et al. (2012, 2013); Shen and Ma (2016); Ma et al. (2015); Calder-Travis and Ma (2020))^1^. Yet, the Bayes-optimal model typically performs very well, not only compared to other models but also in absolute terms, e.g. according to visual inspection of summary statistics^2^. The latter should not be underestimated; in the present paper, for example, it is actually quite stunning that with “only” 5 parameters per subject, the large set of psychometric curves in the panels of Figure 4 are simultaneously well accounted for. Thus, to illustrate the success of the Bayesian modeling approach, visual search seems to be at least as good an example domain as the traditionally cited domain of cue combination (where, occasionally, large deviations from optimality are observed (e.g. Battaglia, Jacobs, and Aslin (2003)). At least in these two subfields, Bayesian models are as “unreasonably effective” as mathematics is in the natural sciences more generally (Wigner, 1960).

Zooming out, we would like to argue that any quantitative process model of a cognitive or perceptual behavior - Bayesian or not - is an improvement over descriptive theories, specifically ones that are only formulated in words and not in equations. Mathematical formalization implies clear commitments to assumptions and predictions, and avoids the mercurial vagueness associated with verbal descriptions (Lewandowsky & Farrell, 2011; Fitch, 2014; Levenstein et al., 2020). Process models or observer models have a special status because they follow the flow of information as it could plausibly occur in the brain; specifically, they commit to an internal representation of the stimuli that is accessible to the observer. The theory of Duncan and Humphreys (1989), while pioneering, offered “only” a word conceptualization of visual search with heterogeneous distractors. On the one hand, it may be disconcerting that in the past 30 years, so little has been done to upgrade their model to a quantitative process model. On the other hand, such a state of affairs is not unexpected: almost wherever one looks in psychology, from perceptual illusions (e.g. Weiss, Simoncelli, and Adelson (2002); Goldreich (2007); Howe and Purves (2005)) to pragmatic reasoning (Frank & Goodman, 2012) to action understanding (Baker, Saxe, & Tenenbaum, 2009), it is quite common for decades to elapse between word theories and quantitative process models. If nothing else, this speaks to the rich future of computational approaches in psychology and cognitive science.

## Acknowledgements

This work was supported by NIH grant R01EY020958 to Wei Ji Ma. We are grateful for useful conversations with Marisa Carrasco, as well as members of the Ma lab, present and past, especially Joshua Calder-Travis, Aditi Singh, Luigi Acerbi, Emin Orhan, Bas van Opheusden, Will Adler, Aspen Yoo, Maija Honig, Yanli Zhou, Heiko Schütt, Jenn Lee, Xiang Li, Peiyuan Zhang and Ionatan Kuperwajs. This work has utilized the NYU IT High Performance Computing resources; we acknowledge the support of its staff members, especially Shenglong Wang.

## A Appendix: Details of the generative models

### A.1 Relationship between concentration parameter *κ* and precision *J*

The inverse of the concentration parameter *κ_i_* is related to the noise on item *i*. As described in earlier studies (Ma et al., 2011), resource in this framework is conceptualized as the Fisher information, which represents a way to quantify the amount of information a measurement, *x_i_*, carries about a variable it depends on, here *s_i_* and is defined as the expected value of:

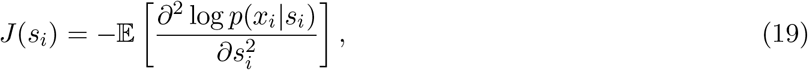

which is evaluated to:

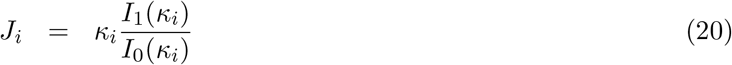

The relationship above between precision *J* and noise *k* is a monotonic function, and thus invertible, allowing for numerical calculation of *k* as a function of *J*. An additional assumption in the variable-precision encoding model (van den Berg et al., 2012) is that the precision *J* varies across trials and items, specifically according to a gamma distribution with shape parameter 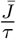 and scale parameter *τ*, and thus with mean 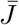.

### A.2 Derivation of the local decision rule

As we said in the main text, in the inference process, the optimal observer inverts the generative model and computes the posterior *p*(*L*|**x**), with *L* = 1, 2… *N*. The observer reports the location *L* for which *p*(*L*|**x**) has the highest value.

Since *p*(*s_L_*) is uniform on 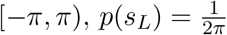 and because 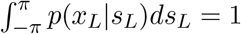, we get:

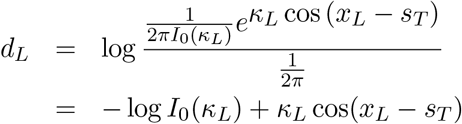

We unpack *p*(**x**) to obtain the normalization term of the posterior *p*(*L*|**x**):

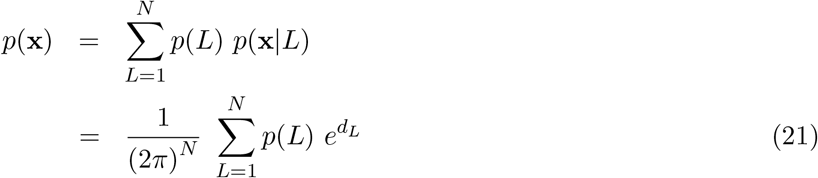

Plugging Equations (7) and (21) into Equation (3), the posterior *p*(*L*|**x**) becomes:

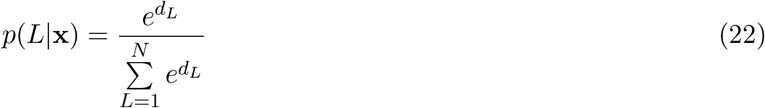

The optimal observer will perform maximum-a-posteriori (MAP) estimation in a deterministic fashion and pick the value 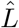 for which the posterior takes the maximum value, which is equivalent to picking the corresponding *d_L_*:

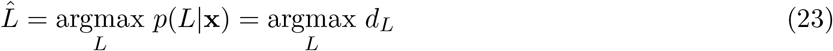

### Detection

In detection as opposed to localization, the target is only half the time present in the display. The observer might have a prior over the probability of target presence which is different than 0.5; alternatively the observer might have a higher motor cost for pressing a button with one hand as opposed to the other. Another possibility is that the observer might have an asymmetric utility for correct detection vs correct rejection. We incorporate all these possible contributions to bias into a single parameter, *p*(*C* = 1) = *p*_present_ and write *d* in terms of the prior and the log likelihood ratio:

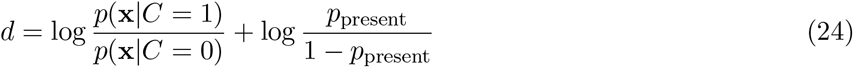

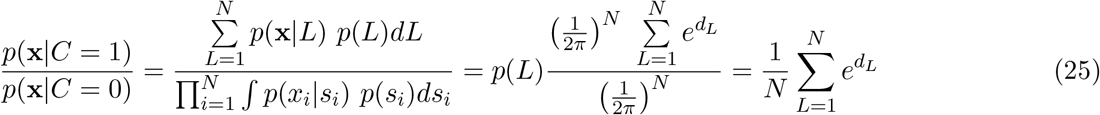

Thus, luckily, the global decision variable d can be written as a function of the local decision variables:

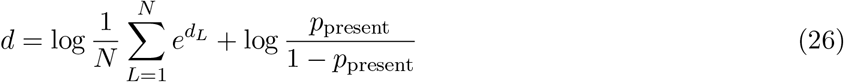

## B Appendix: Distributions of summary statistics and their correlations

The distributions of each one of the three main summary statistics (B-D from Table 2) when targets and distractors are uniformly distributed are presented as below (Figure A1A). To further illustrate the point that other distributions of targets and distractors yield different distributions of summary statistics, we include a von Mises distribution with parameters as in Calder-Travis and Ma (2020) (Figure A1B). Note that the bump for the distractor variance distribution at set size *N* = 6 in Figure A1B captures a pattern in the distribution and is not due to insufficient sampling. For more discussion about the source of this bump, see (Calder-Travis & Ma, 2020).

**Figure A1:**
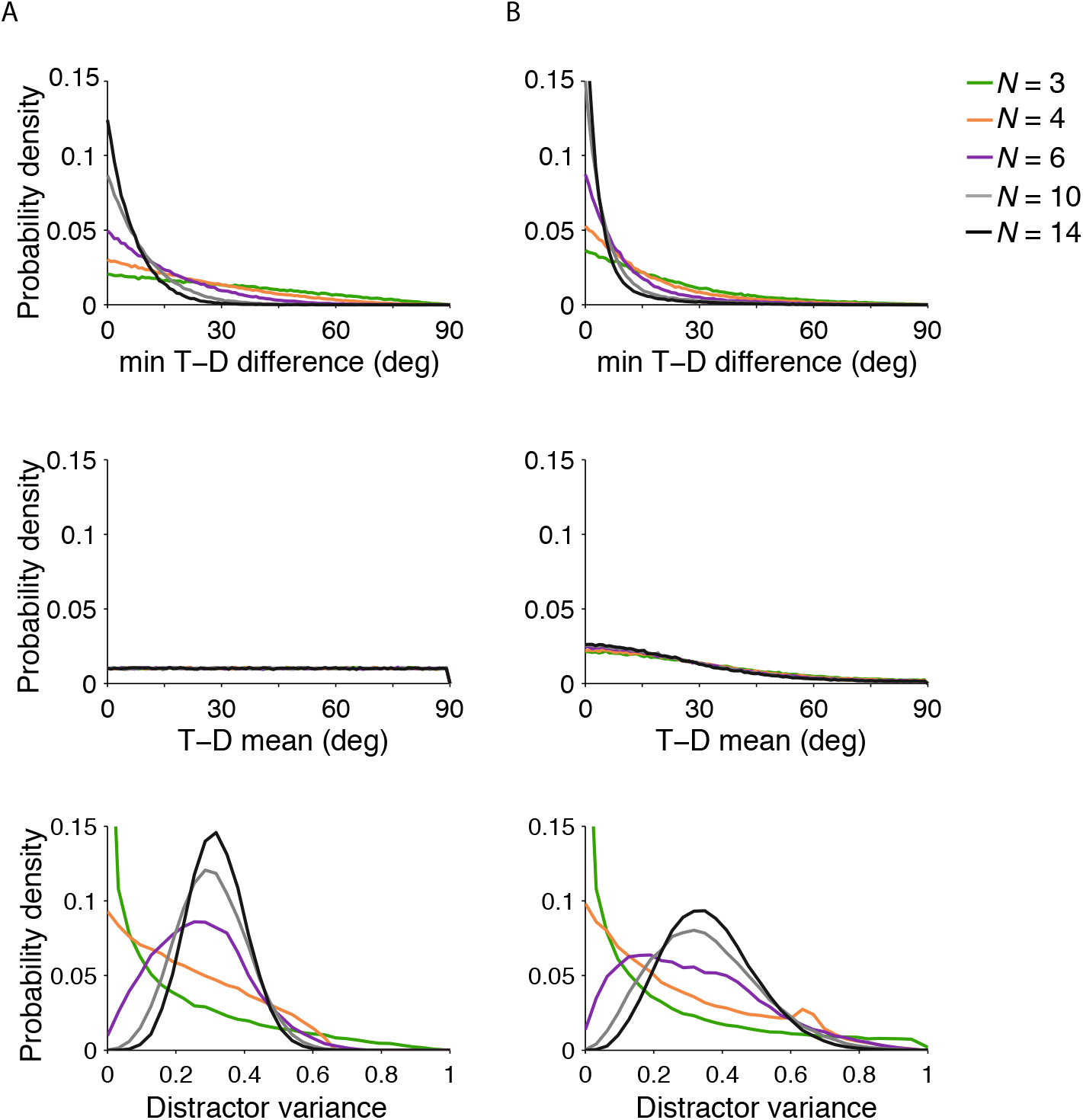
Distributions of the summary statistics (B, C and D, introduced in Table 2) in the localization task, with **A)** uniform distributions of targets and distractors and **B)** von Mises distribution with mean 0 and concentration parameter 1.5. We simulated 100,000 trials and calculated these distributions and correlations across set sizes. Note that in the actual experiment observers were presented with only with set sizes up to 6.

Subsequently, the two types of distributions from above also yield different patterns of correlations (Table A1A vs Table A1B). The correlations between min T-D difference and T-D mean start high and decrease with set size for the uniform distribution, while they stay high even as set size increases for this von Mises distribution. The correlations between min T-D difference and distractor variance decrease with set size for the uniform but are all near 0 for this von Mises distribution.

**Table A1:** Spearman correlations between summary statistics for 100,000 simulated localization displays with distractors following **A) uniform** and **B) von Mises** distributions.

|  |  |  |  |  |  |  |
| --- | --- | --- | --- | --- | --- | --- |
| A) | Correlation between | N = 3 | N = 4 | N = 6 | N = 10 | N = 14 |
|  | min T-D difference and T-D mean | 0.469 | 0.336 | 0.217 | 0.129 | 0.095 |
|  | min T-D difference and distractor variance | -0.171 | -0.138 | -0.082 | -0.038 | -0.019 |
|  | T-D mean and distractor variance | -0.003 | 0.001 | -0.002 | -0.003 | -0.001 |
| B) | Correlation between | N = 3 | N = 4 | N = 6 | N = 10 | N = 14 |
|  | min T-D difference and T-D mean | 0.605 | 0.553 | 0.510 | 0.484 | 0.472 |
|  | min T-D difference and distractor variance | -0.035 | 0.008 | -0.025 | 0.025 | 0.020 |
|  | T-D mean and distractor variance | 0.049 | 0.035 | 0.023 | 0.002 | 0.006 |

## C Appendix: Logistic regression models : accuracy with several possible combinations of summary statistics

**Table A2:** The 8 logistic regression models and the summary statistics included as factors in each. We employed these models in the model comparison method from Shen and Ma (2019) to determine factor *usefulness* and *necessity*.

| Regression model name | Regressor | Regressor | Regressor | Regressor |
| --- | --- | --- | --- | --- |
| Base model | $N$ | | | |
| Knock-in min T-D difference | $N$ | min T-D difference | | |
| Knock-in T-D mean | $N$ | T-D mean | | |
| Knock-in distractor variance | $N$ | distractor variance | | |
| Full model | $N$ | min T-D difference | T-D mean | distractor variance |
| Knock-out min T-D difference | $N$ | min T-D difference | T-D mean | |
| Knock-out T-D mean | $N$ | min T-D difference | | distractor variance |
| Knock-out distractor variance | $N$ | | T-D mean | distractor variance |

## D Appendix: Calculation of quantile bins for the psychometric curve of proportion correct with min T-D difference

We plot the psychometric curves metrics as a function of the target - most similar distractor orientation difference (min T-D difference). We denote the min T-D difference distribution by *p*(*θ*). Its cumulative density function (cdf) is *P*(min *a_i_* < *θ*). Quantile binning of this distribution is important to ensure that there will be enough data points in each bin and thus the psychometric curve is reliable. The quantile function is the inverse of the cdf, which we can analytically compute.

The cdf will vary with condition and set size. The search arrays have set sizes *N* = 2, 3, 4, 6, and with conditions “target present” and “target absent”, the absolute orientation differences *a_i_* (*i* = 1 — N) could have the length anywhere between *N* = 1 — 6. Both foveally presented targets and peripherally presented search array stimuli are independently drawn from uniform distributions on (–*π,π*), so *a_i_* ∈ (0, *π*).

We will decompose the cdf P(*min a_i_* < *θ*) as below. min *a_i_* < *θ* is contradicted when ∀*a_i_* > *θ*, for *i* = 1 – *N*. Taking into account that the *a_i_* values are independently drawn allows for the expression below to be written as a product over *i*. Since the search arrays have set sizes *N* = 2, 3, 4, 6 with both “target present” and “target absent” conditions, we could have anywhere between *N* = 1 – 6 angle differences *a_i_*.

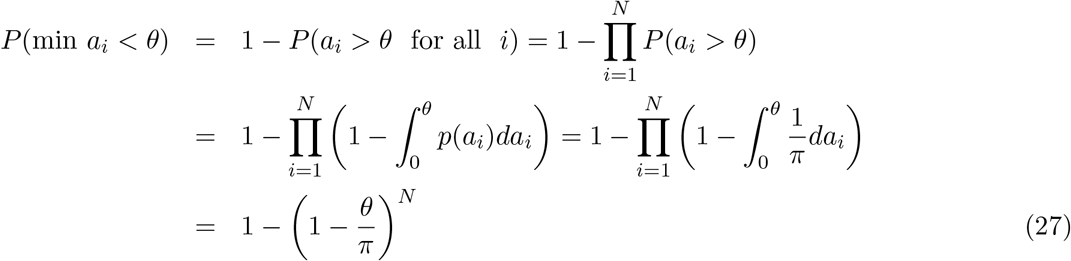

Here we show 5 quantiles *p* = [0, 1/5, 2/5 …1] in Figure A2. More generally, for quantiles *p* ∈ (0,1), we solve for the quantile orientation values *θ_p_* that satisfy *P*(min *a_i_* < *θ_p_*) = *p* to get:

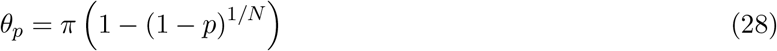

**Figure A2:**
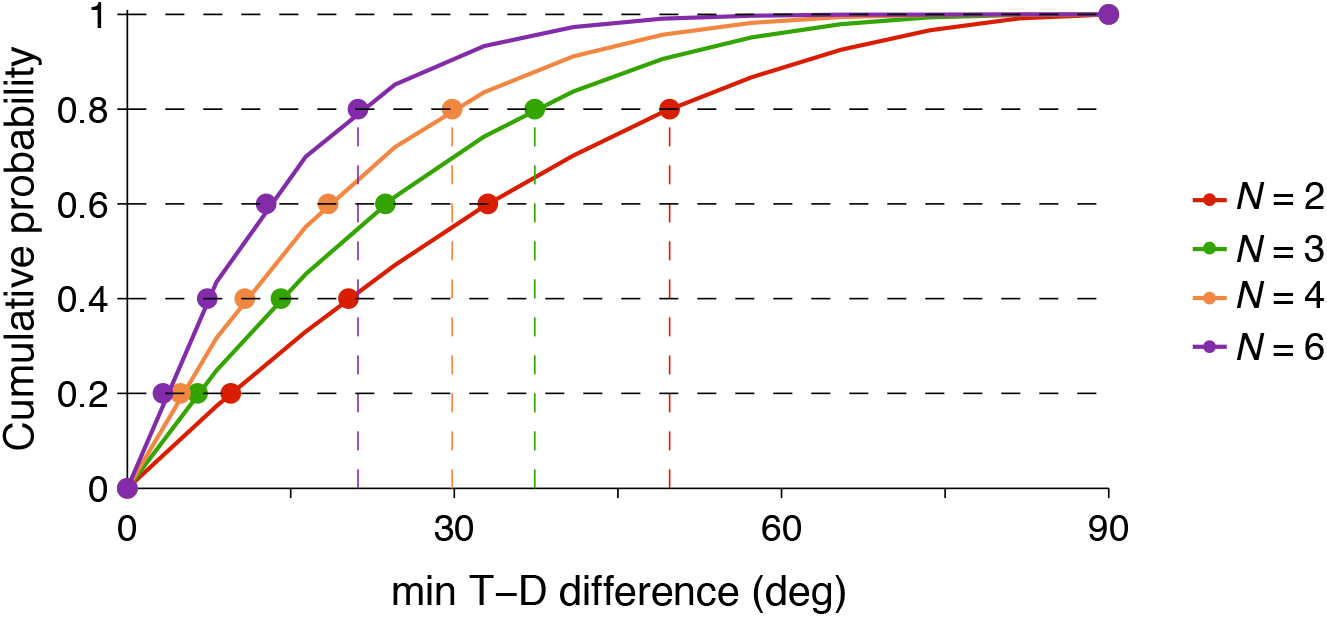
min T-D difference cumulative distributions and visualization of the quantile placement of the bins for each set size. Projecting each dot onto the x-axis yields the min T-D difference value of each quantile bin. Here we illustrate with the colored dashed lines how the placement of quantile bin number 5 falls onto the x axis for each set size.

For the other metrics, target-distractor mean (T-D mean) and circular variance of the distractors, we calculated empirically the quantile bins for each individual subject and condition as analytical expressions were not straightforward to calculate.

## E Appendix 3: Joint fits of Model 1 across localization and detection for Experiment 1

Figure 8 in main already include Figure A3B and A4B; here we also present data and the joint model 1 fits with respect to all the summary statistics.

**Figure A3:**
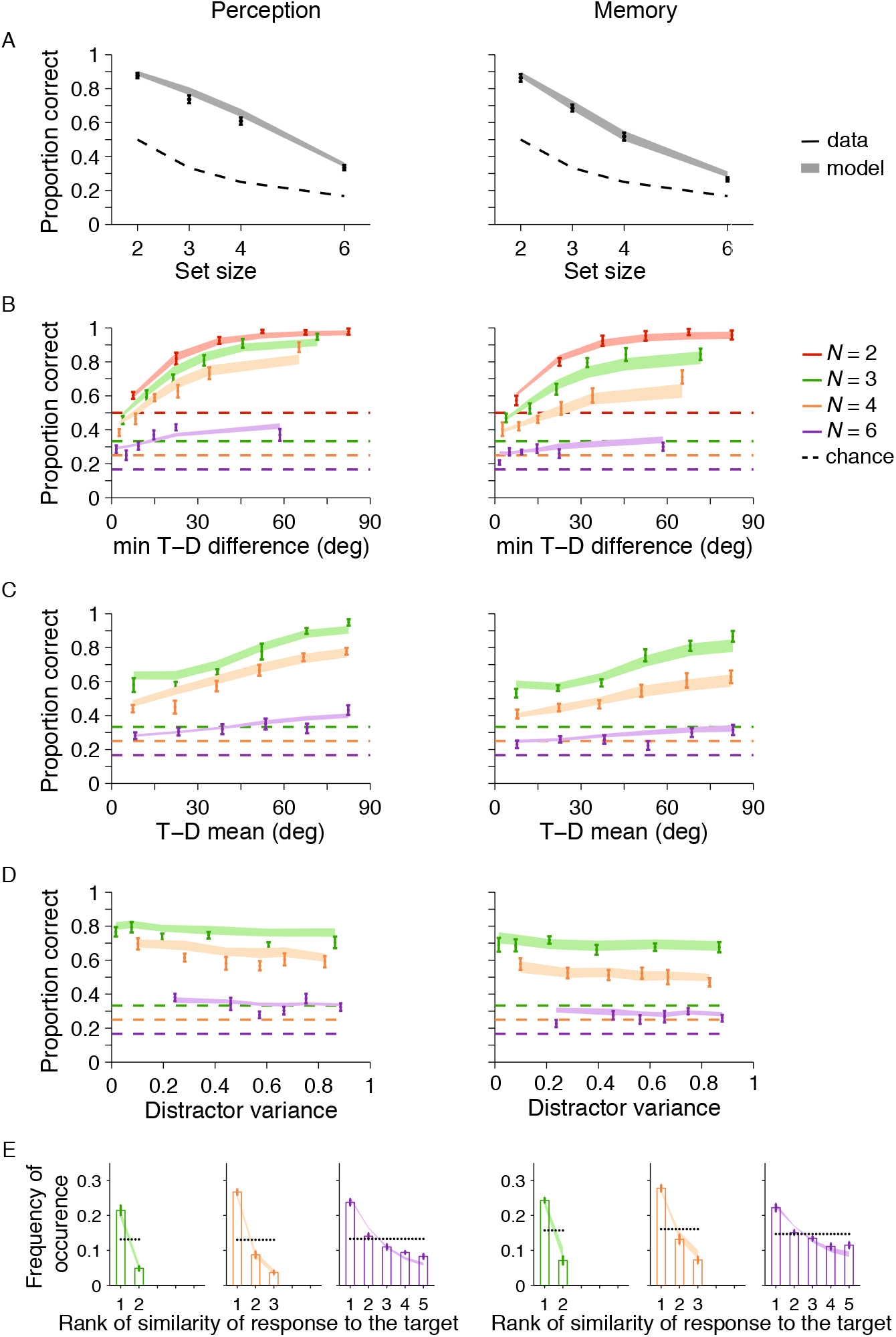
Target localization condition in Experiment 1: data (mean sem and joint model fits. The joint model fits the data very well. Perception (Left) and memory (Right). **(A)** Proportion correct with set size. **(B)** Proportion correct with the min T-D difference difference. The box indicates the summary statistic that is the most diagnostic of performance. **(C)** Proportion correct with the T-D mean. **(D)** Proportion correct as a function of circular variance of the distractors. **(E)** Frequency of occurrence of chosen (incorrect) item by rank of similarity to the target.

**Figure A4:**
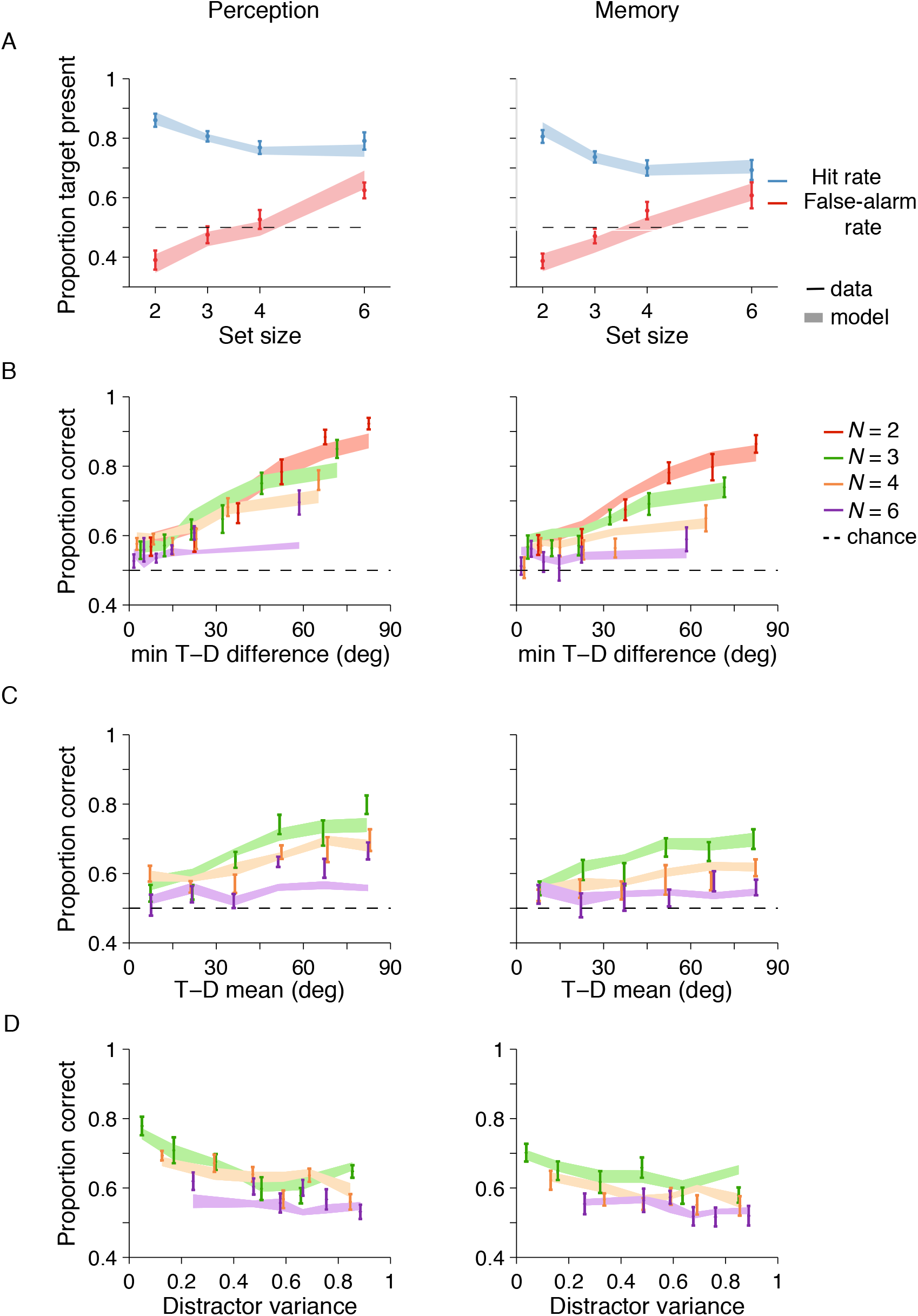
Target detection condition in Experiment 1: data (mean sem) and joint model fits. The joint model captures the data relatively well. **(A)** Proportion correct as a function of set size. **(B)** Proportion correct as a function of the min T-D difference. **(C)** Proportion correct with T-D mean. **(D)** Proportion correct as a function of the circular variance of the distractors.

## F Appendix 4: Results of Experiment 2 (small spacing)

### F.1 Experiment 2: Target localization

In target localization, the results from experiment 2 mostly replicate the patterns of results from experiment 1. We see this both qualitatively in Figure A5 as well as in the logistic regression factor importance analysis (Figure A6): the min T-D difference is the most useful regressor, though neither regressor is necessary.

**Figure A5:**
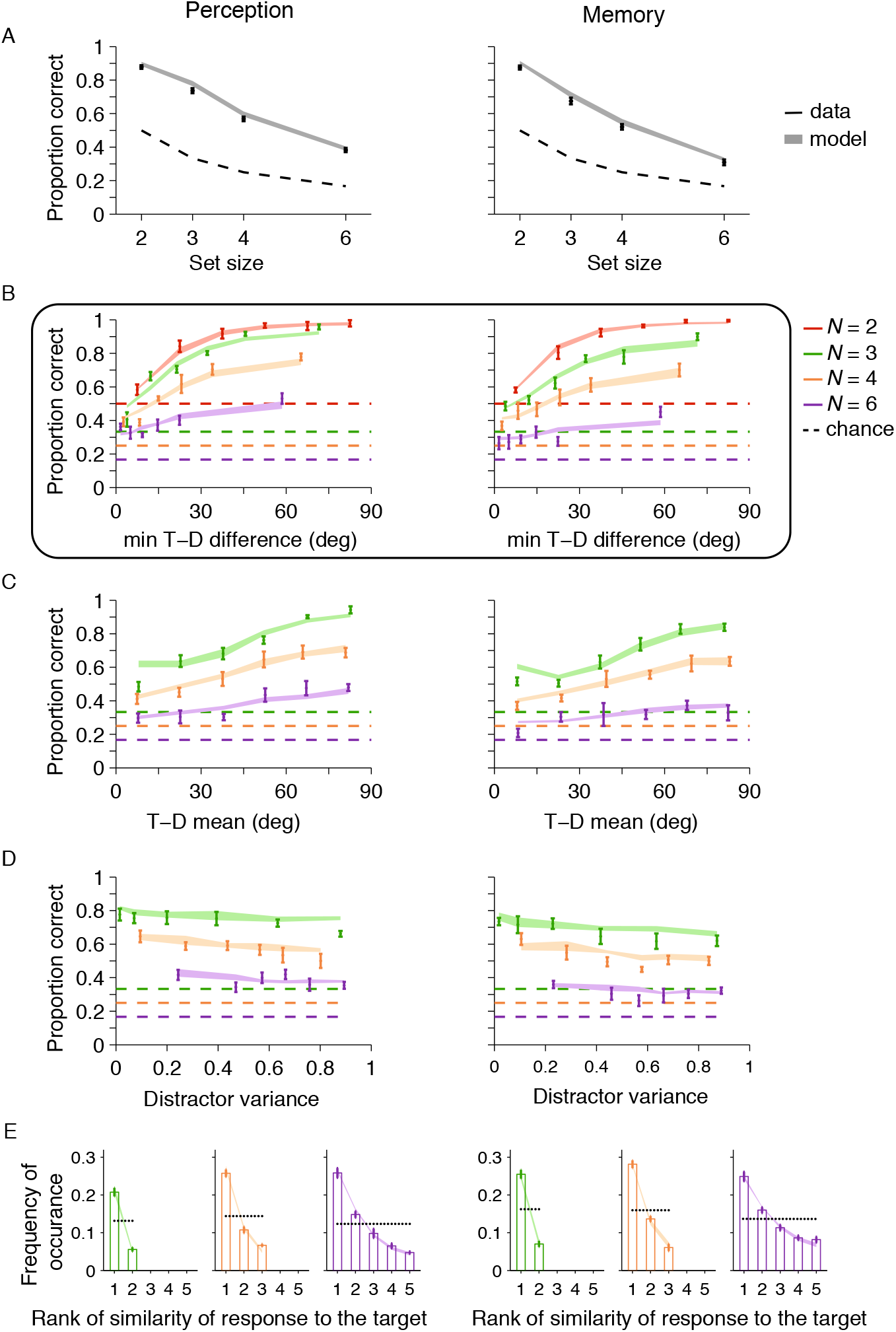
Results from the localization condition in Experiment 2: data and model 1 fits. As in experiment 1, proportion correct decreases with set size and increases most saliently with min T-D difference. **(A)** Proportion correct with set size. **(b)** Proportion correct with min T-D difference. **(C)** Proportion correct with T-D mean. **(D)** Histogram of responses by rank of similarity to target. **(E)** Proportion correct does not seem to vary with the increasing heterogeneity of the distractors, as quantified by their circular variance. **(Left)**: Perception and **(Right)**: Memory.

**Figure A6:**
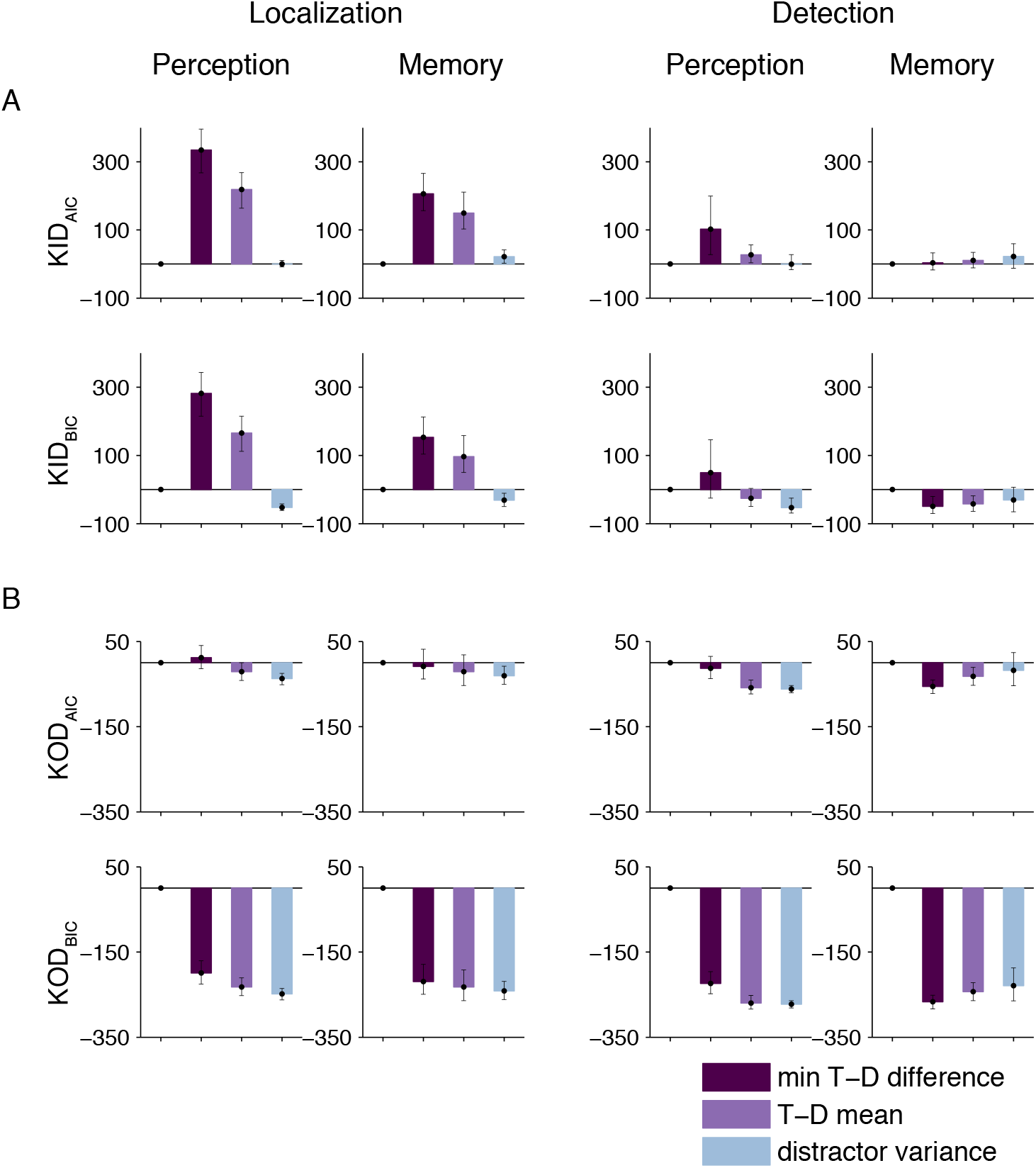
Experiment 2: Factor importance for the regressors: min T-D difference, T-D mean and distractor variance. min T-D difference was the most useful factor in 3/4 conditions (vs 4/4 in Experiment 1); distractor variance was the most useful factor in Detection - Memory. Neither factor was necessary. **A)** Factor usefulness : sum across subjects and bootstrapped confidence intervals of the knock-in difference (KID) of the model with the factor from the base model, for both AIC (top) and BIC (bottom). **B)** Factor necessity: sum across subjects and bootstrapped confidence intervals of the knock-out difference (KOD) of the full model from the model without the factor, for both AIC (top) and BIC (bottom).

### F.2 Experiment 2: Target detection

In experiment 2 detection, the influence of the summary statistics on performance also replicated to a good extent the pattern from experiment 1. However, we note an exception in the case of Detection - Memory, where distractor variance may be the most useful regressor (Figure A6).

**Figure A7:**
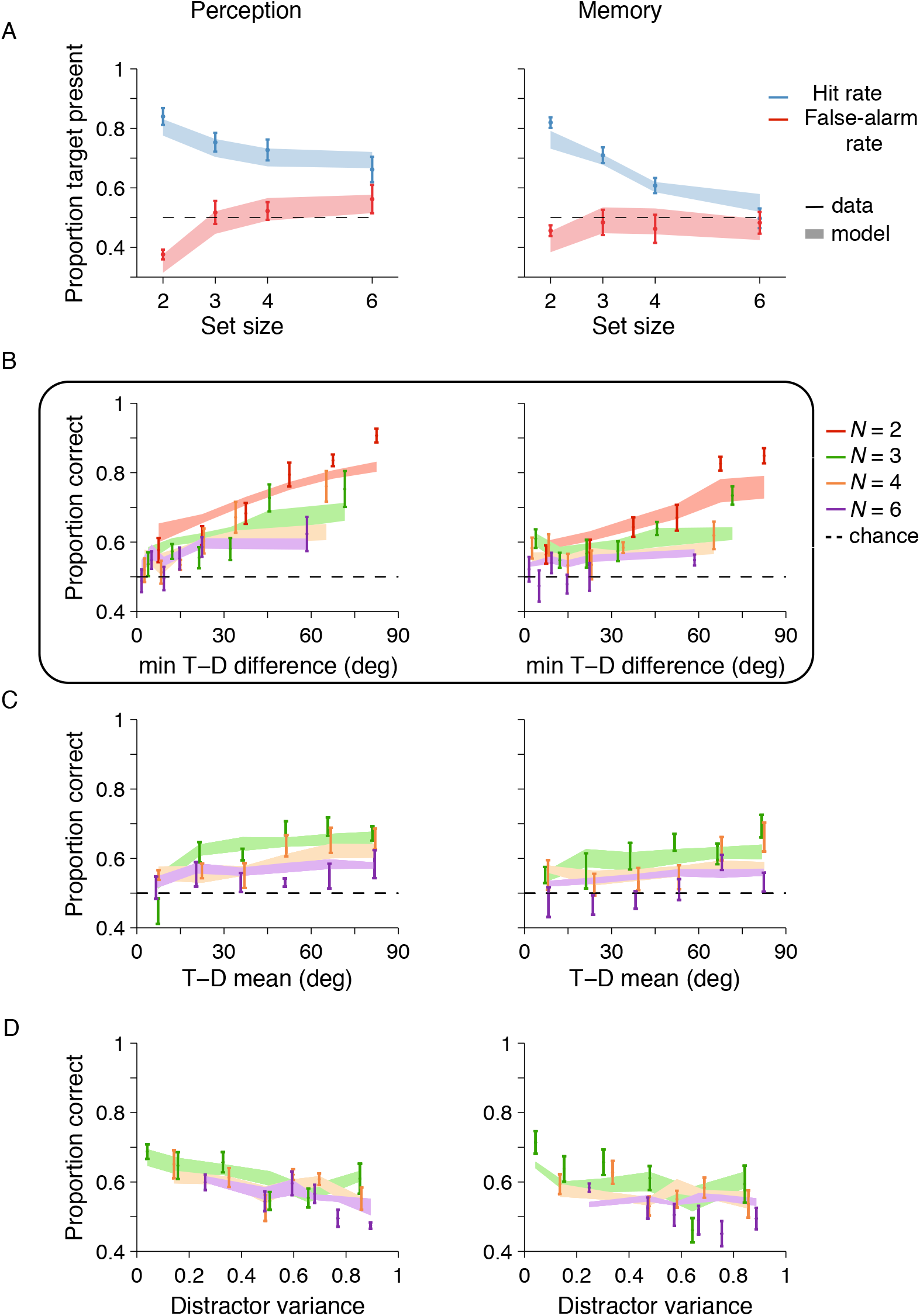
Results from the detection condition in Experiment 2: data and model 1 fits. As in experiment 1, proportion correct increases most saliently with min T-D difference. **(a)** Proportion correct with set size. **(b)** Proportion correct with the min T-D difference.

### F.3 Experiment 2: Model 2 with decision noise

We again replicate the pattern of results from experiment 1, as decision noise is not necessary in localization, but in the case of detection it improves the model fits.

**Figure A8:**
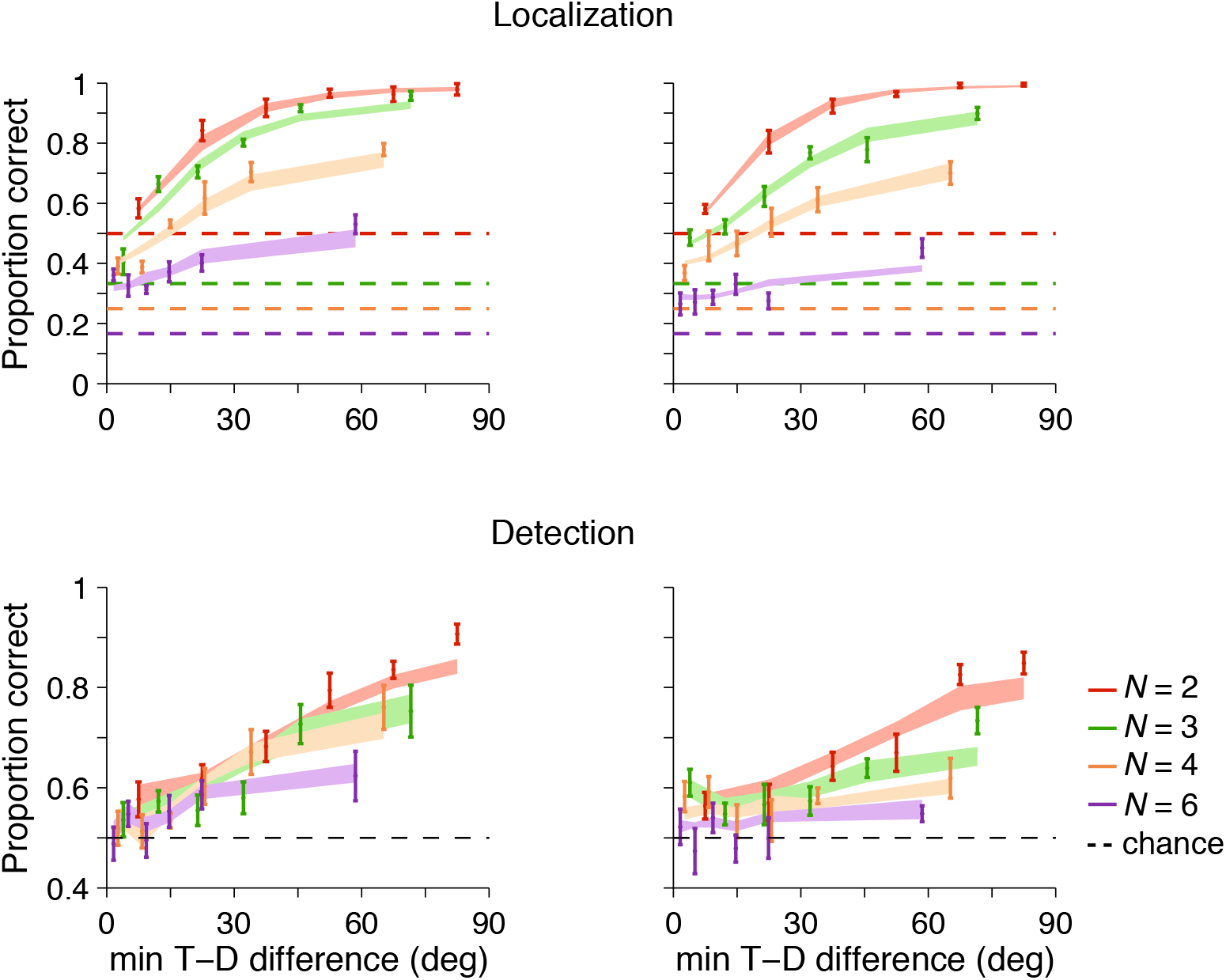
Experiment 2: addition of decision noise independently to localization and then to detection: data and model 2 fits. As in experiment 1, model 2 which includes decision noise provides a better fit to the detection data.

### F.4 Experiment 2: Target localization and target detection, joint fitting with decision noise

As we see in Figure 9B, the joint model fits also capture the data well in Experiment 2. Here we add decision noise to the joint fitting of the localization and detection data from Experiment 2.

### F.5 Joint fitting: model 2 fits

**Figure A9:**
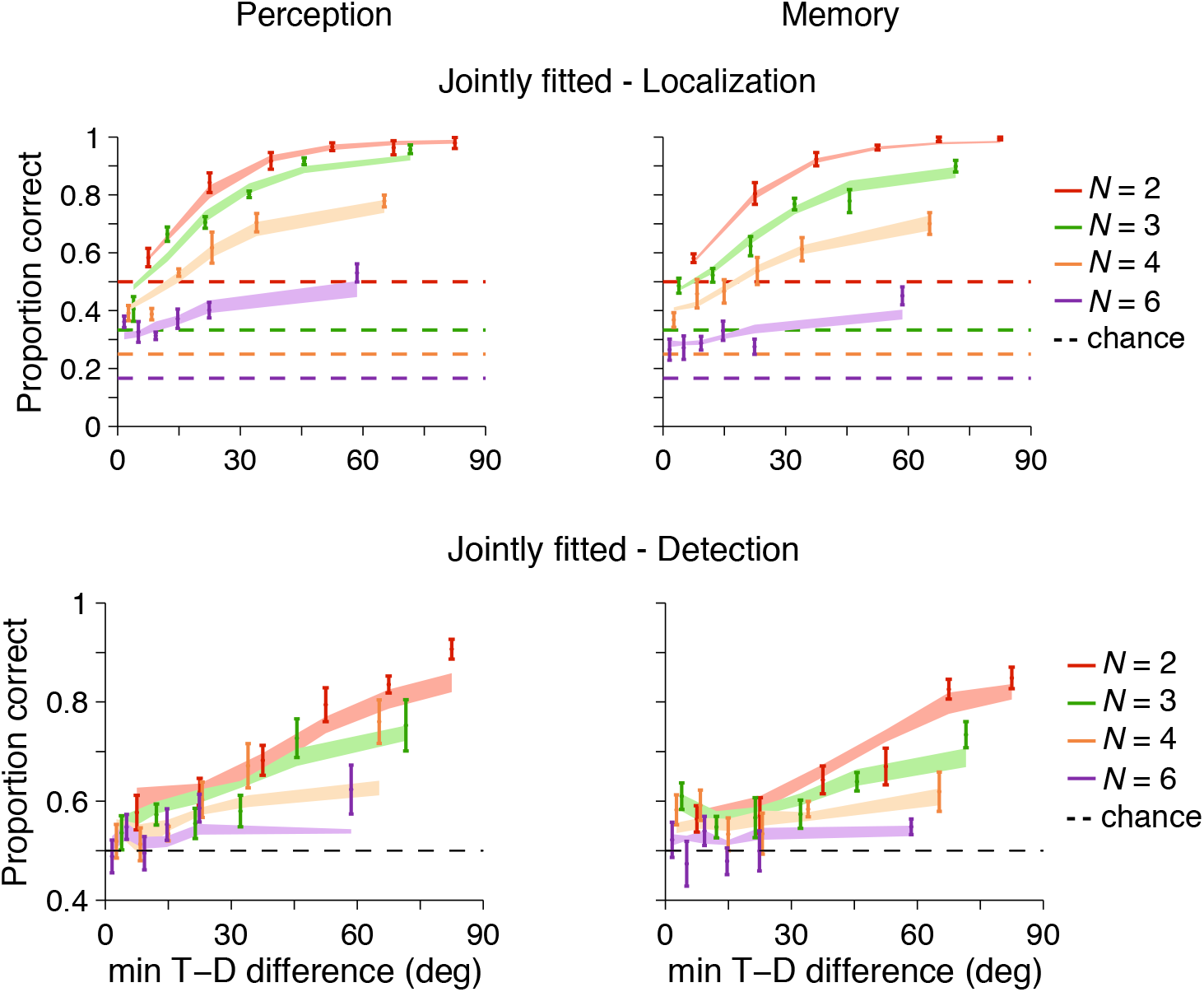
Experiment 2: part of the data and joint fits of model 2. As in experiment 1, the joint model captures the data relatively well, especially for localization.

## G Appendix: Model comparison and model parameters from model 2

### G.1 Model comparison

**Figure A10:**
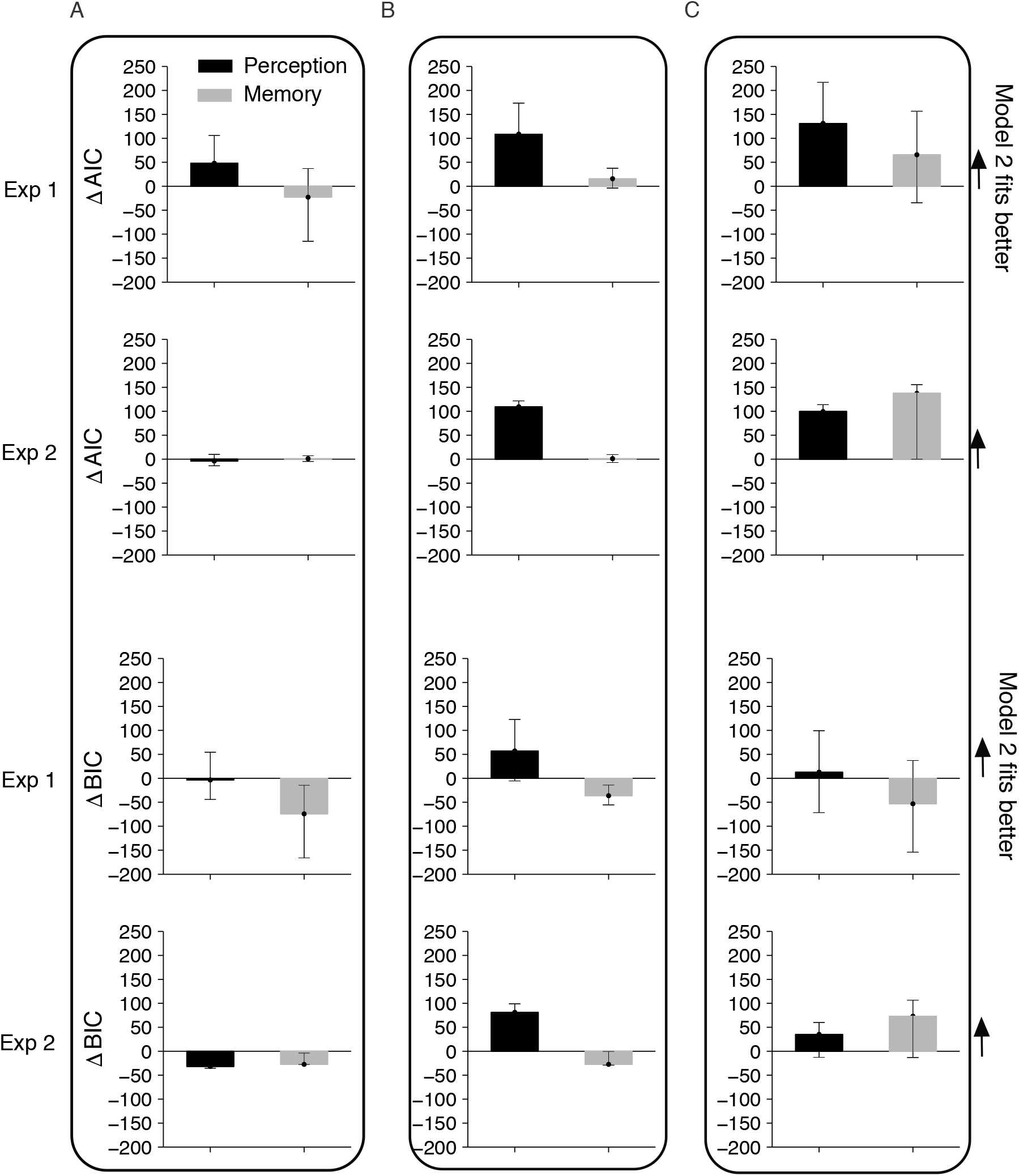
Model comparison shows that model 2 with decision noise is not necessary for localization, but improves the model fits for Detection - Perception in both experiments and for the joint fits in Experiment 2. **(A)** Localization. **(B)** Detection. **(C)** Joint localization and detection. The y-axis shows the ΔAIC = AIC_MI_ – AIC_M2_ (top) and similarly ΔBIC (bottom), specifically their sum across subjects with the bootstrapped 95 % confidence intervals.

### G.2 Model 2 parameters

#### G.2.1 Precision parameters

The precision parameters from model 2 follow similar patterns as the precision parameters from model 1 depicted in main (Figure 10).

**Figure A11:**
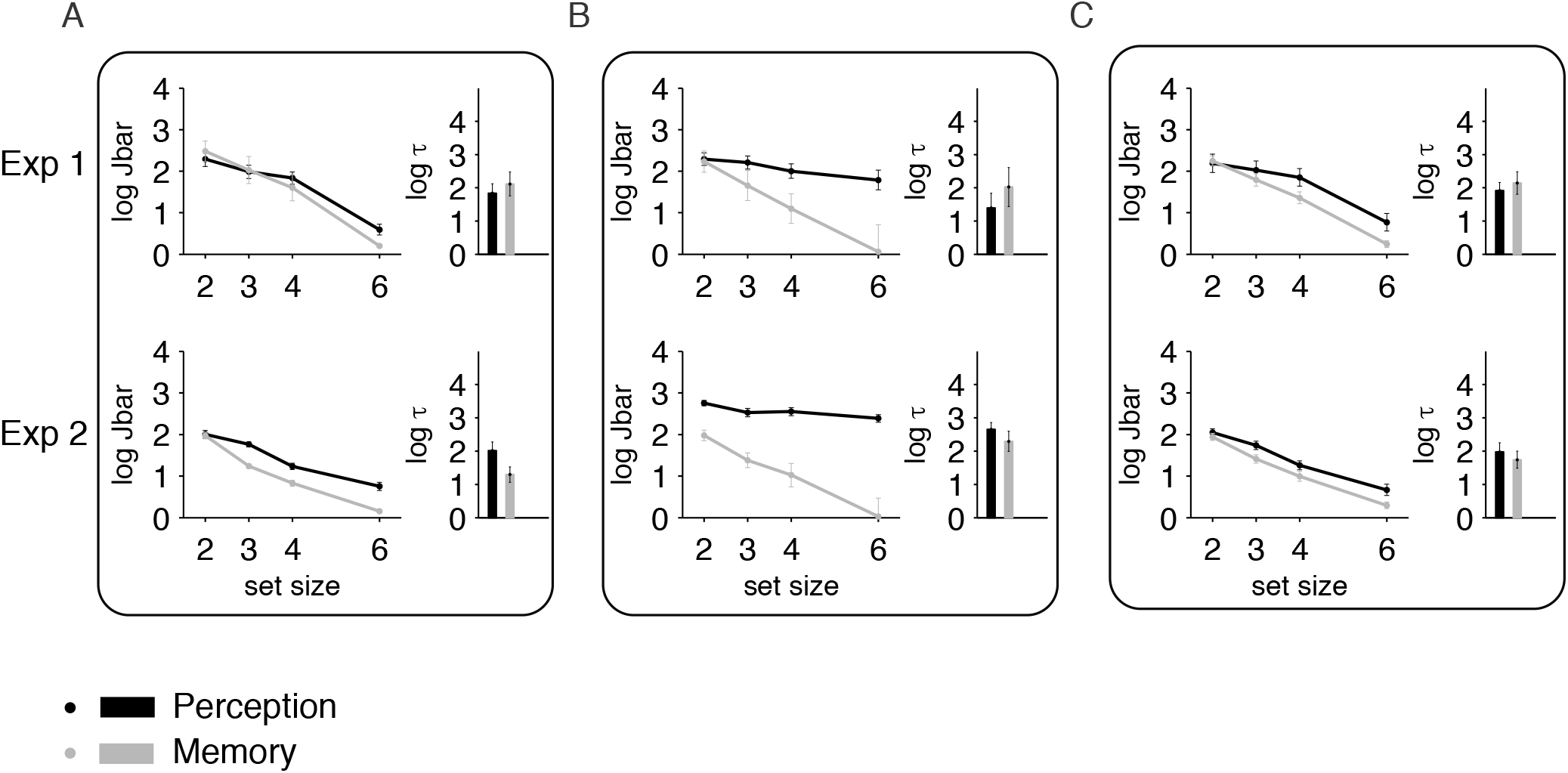
Estimates of precision parameters in model 2 across conditions and experiments: as for model 1, mean precision decreases with set size and is higher in perception than memory. **(A)** Localization. **(B)** Detection. **(C)** Joint localization and detection.

#### G.2.2 Decision noise

**Figure A12:**
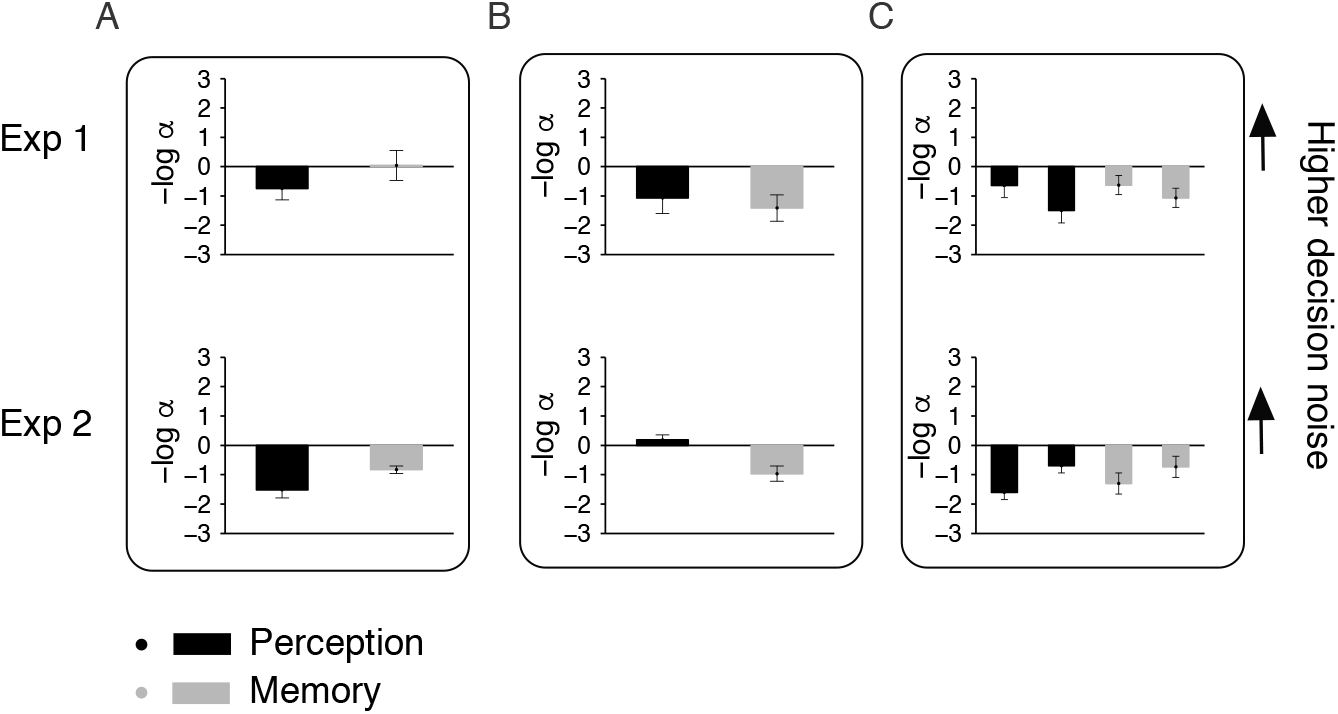
Decision noise parameters from model 2. **(A)** Localization. **(B)** Detection. **(C)** Joint localization and detection, within each Perception/Memory the first bar is *α*_Loc_ and the second is *α*_Det_.

## H Appendix: Reaction times

### H.1 Experiment 1

**Figure A13:**
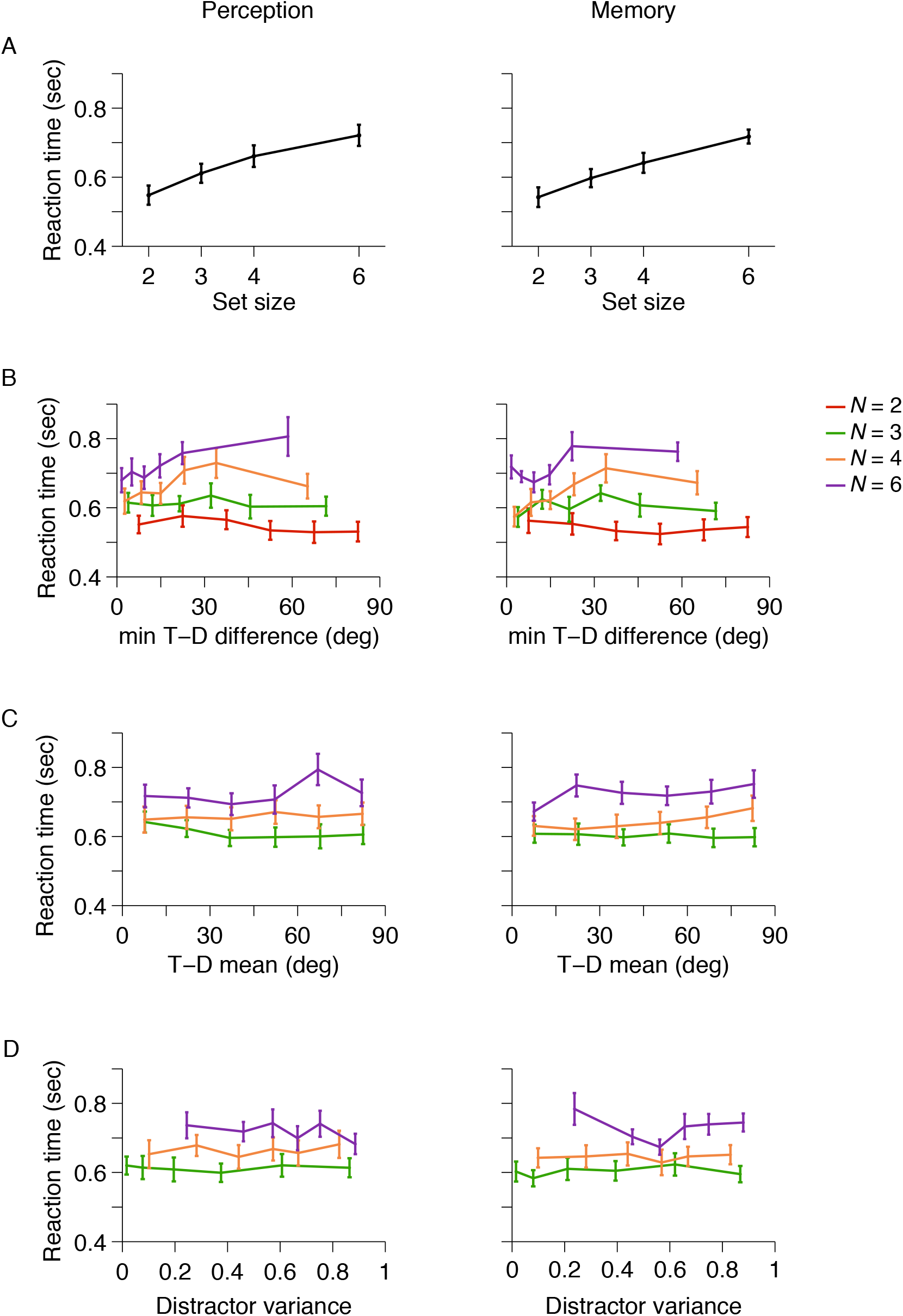
Reaction times for localization in Experiment 1. Reaction times increase with set size (A).

**Figure A14:**
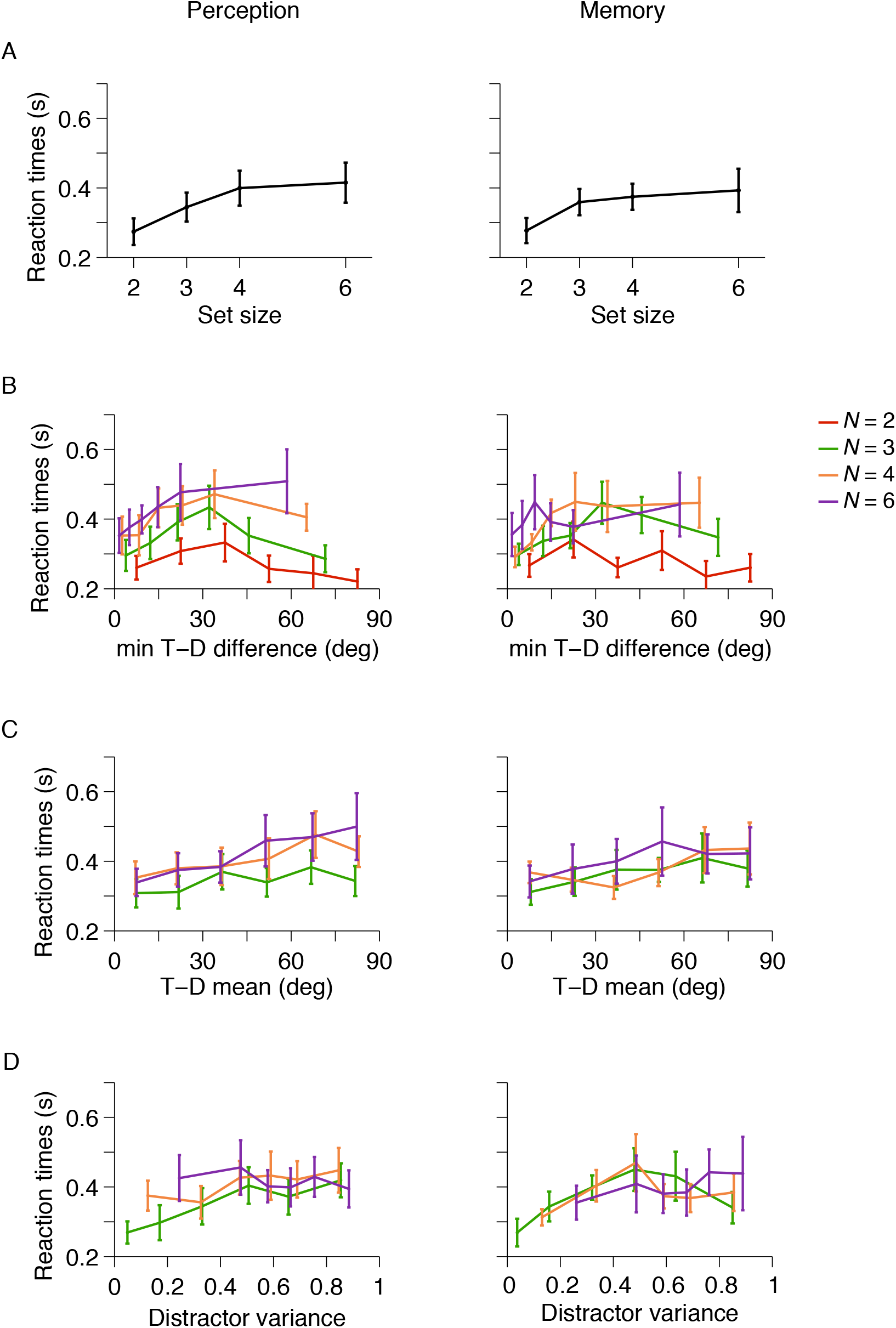
Reaction times for detection in Experiment 1. Reaction times increase with set size (A).

### H.2 Experiment 2

**Figure A15:**
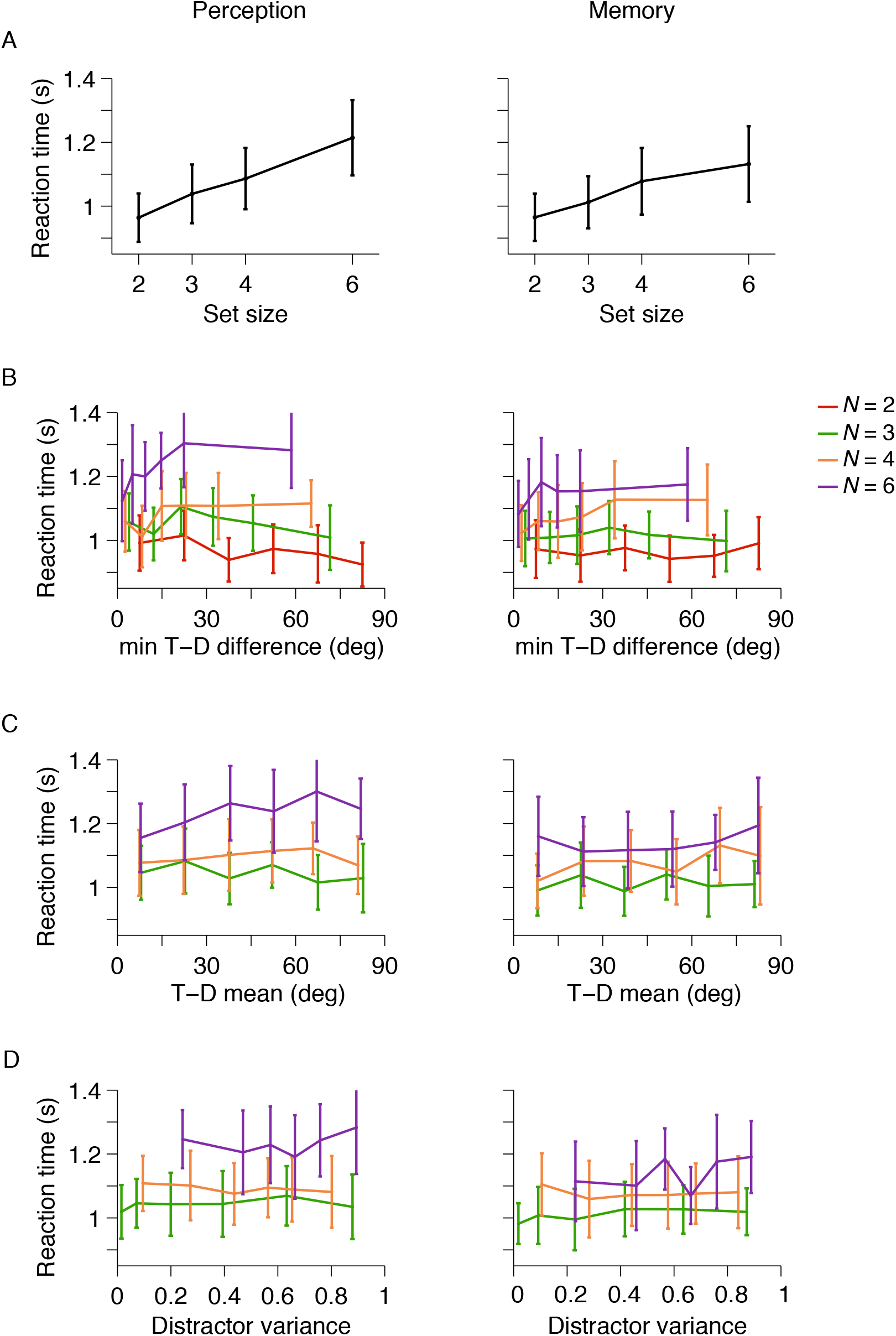
Reaction times for localization in Experiment 2. Reaction times increase with set size (A).

**Figure A16:**
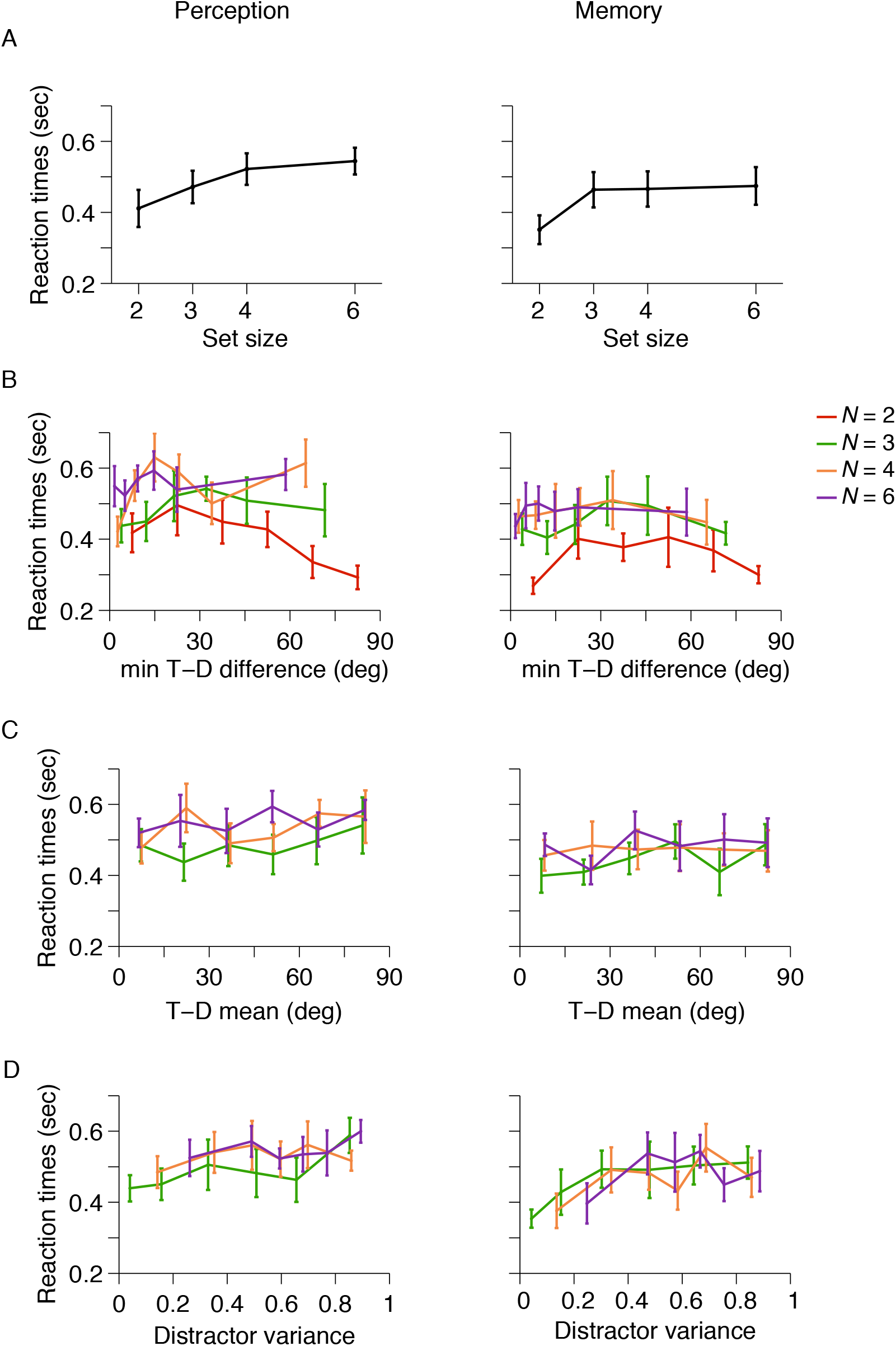
Reaction times for detection in Experiment 2. Reaction times increase with set size (A).

## Footnotes

1 Navalpakkam and Itti (2007); Vincent et al. (2009) are not listed here because they did not test the Bayes-optimal model; for details, see Ma et al. (2015)

2 To capture behavior across multiple set sizes, as in the present paper, one typically does need to assume that precision depends on set size; that, however, makes sense in a resource-constrained system (van den Berg & Ma, 2018)

## References

Acerbi, L., & Ma, W. J. (2017). Practical bayesian optimization for model fitting with bayesian adaptive direct search. Advances in Neural Information Processing Systems (30), 1834–1844.

Acerbi, L., Vijayakumar, S., & Wolpert, D. M. (2014). On the origins of suboptimality in human probabilistic inference. PLoS Computational Biology, 10(6), e1003661. doi: 10.1371/journal.pcbi.1003661

Ahmad, J., Swan, G., Bowman, H., Wyble, B., Nobre, A. C., Shapiro, K. L., & McNab, F. (2017). Competitive interactions affect working memory performance for both simultaneous and sequential stimulus presentation. Scientific Reports, 7(1). doi: 10.1038/s41598-017-05011-x

Akaike, H. (1974, December). A new look at the statistical model identification. IEEE Transactions on Automatic Control, 19(6), 716–723. Retrieved from https://doi.org/10.1109/tac.1974.1100705 doi: 10.1109/tac.1974.1100705

Avraham, T., Yeshurun, Y., & Lindenbaum, M. (2008). Predicting visual search performance by quantifying stimuli similarities. Journal of Vision, 8(4), 9. doi: 10.1167/8.4.9

Baddeley, A. D., & Hitch, G. (1974). Working memory. In Psychology of learning and motivation (pp. 47–89). Elsevier. doi: 10.1016/s0079-7421(08)60452-1

Baker, C. L., Saxe, R., & Tenenbaum, J. B. (2009, December). Action understanding as inverse planning. Cognition, 113(3), 329–349. Retrieved from https://doi.org/10.1016/j.cognition.2009.07.005 doi: 10.1016/j.cognition.2009.07.005

Battaglia, P. W., Jacobs, R. A., & Aslin, R. N. (2003, July). Bayesian integration of visual and auditory signals for spatial localization. Journal of the Optical Society of America A, 20(7), 1391. Retrieved from https://doi.org/10.1364/josaa.20.001391 doi: 10.1364/josaa.20.001391

Bays, P. M. (2016). Evaluating and excluding swap errors in analogue tests of working memory. Scientific Reports, 6(1). doi: 10.1038/srep19203

Beck, J. M., Ma, W. J., Pitkow, X., Latham, P. E., & Pouget, A. (2012, April). Not noisy, just wrong: The role of suboptimal inference in behavioral variability. Neuron, 74(1), 30–39. doi: 10.1016/j.neuron.2012.03.016

Berens, P. (2009). CircStat: A MATLAB Toolbox for circular statistics. Journal of Statistical Software, 31 (10). Retrieved from https://doi.org/10.18637/jss.v031.i10 doi: 10.18637/jss.v031.i10

Bergen, J. R., & Julesz, B. (1983). Rapid discrimination of visual patterns. IEEE Transactions on Systems, Man, and Cybernetics, SMC-13(5), 857–863. doi: 10.1109/tsmc.1983.6313080

Bhardwaj, M., van den Berg, R., Ma, W. J., & Josie, K. (2016). Do people take stimulus correlations into account in visual search? PLOS ONE, 11 (3), e0149402. doi: 10.1371/journal.pone.0149402

Biggs, A. T., & Mitroff, S. R. (2014). Improving the efficacy of security screening tasks: A review of visual search challenges and ways to mitigate their adverse effects. Applied Cognitive Psychology, 29(1), 142–148. doi: 10.1002/acp.3083

Boettcher, S. E. P., Draschkow, D., Dienhart, E., & Võ, M. L.-H. (2018, December). Anchoring visual search in scenes: Assessing the role of anchor objects on eye movements during visual search. Journal of Vision, 18(13), 11. Retrieved from https://doi.org/10.1167/18.13.11 doi: 10.1167/18.13.11

Bouma, H. (1970). Interaction effects in parafoveal letter recognition. Nature, 226(5241), 177–178. doi: 10.1038/226177a0

Bowers, J. S., & Davis, C. J. (2012). Bayesian just-so stories in psychology and neuroscience. Psychological Bulletin, 138(3), 389–414. Retrieved from https://doi.org/10.1037/a0026450 doi: 10.1037/a0026450

Brainard, D. H. (1997). The psychophysics toolbox. Spatial Vision, 10(4), 433–436.

Bravo, M., & Nakayama, K. (1992). The role of attention in different visual search tasks. Perception and psychophysics, 51, 465–72.

Busey, T., & Palmer, J. (2008). Set-size effects for identification versus localization depend on the visual search task. Journal of Experimental Psychology: Human Perception and Performance, 34(4), 790–810. doi: 10.1037/0096-1523.34.4.790

Calder-Travis, J., & Ma, W. J. (2020, January). Explaining the effects of distractor statistics in visual search. bioRxiv. Retrieved from https://doi.org/10.1101/2020.01.03.893057 doi: 10.1101/2020.01.03.893057

Cameron, L. E., Tai, J., Eckstein, M., & Carrasco, M. (2004). Signal detection theory applied to three visual search tasks — identification, yes/no detection and localization. Spatial Vision, 17(4), 295–325. doi: 10.1163/1568568041920212

Daw, N. D., O’Doherty, J. P., Dayan, P., Seymour, B., & Dolan, R. J. (2006). Cortical substrates for exploratory decisions in humans. Nature, 441(7095), 876–879. doi: 10.1038/nature04766

Drugowitsch, J., Wyart, V., Devauchelle, A.-D., & Koechlin, E. (2016). Computational precision of mental inference as critical source of human choice suboptimality. Neuron, 92(6), 1398–1411.

Dukewich, K. R., & Klein, R. M. (2009). Finding the target in search tasks using detection, localization, and identification responses. Canadian Journal of Experimental Psychology/Revue canadienne de psychologie experimentale, 63(1), 1–7. doi: 10.1037/a0012780

Duncan, J. (1989). Boundary conditions on parallel processing in human vision. Perception, 18(4), 457–469. doi: 10.1068/p180457

Duncan, J., & Humphreys, G. W. (1989). Visual search and stimulus similarity. Psychological Review, 96(3), 433–458. doi: 10.1037/0033-295x.96.3.433

Eckstein, M. P. (2011). Visual search: A retrospective. Journal of Vision, 11(5), 14–14. doi: 10.1167/11.5.14

Eckstein, M. P., Thomas, J. P., Palmer, J., & Shimozaki, S. S. (2000, 01). A signal detection model predicts the effects of set size on visual search accuracy for feature, conjunction, triple conjunction, and disjunction displays. Perception and Psychophysics, 62(3), 425–451. doi: 10.3758/BF03212096

Farmer, E. W., & Taylor, R. M. (1980). Visual search through color displays: Effects of target-background similarity and background uniformity. Perception & Psychophysics, 27(3), 267–272. doi: 10.3758/bf03204265

Fitch, W. T. (2014, September). Toward a computational framework for cognitive biology: Unifying approaches from cognitive neuroscience and comparative cognition. Physics of Life Reviews, 11(3), 329–364. Retrieved from https://doi.org/10.1016/j.plrev.2014.04.005 doi: 10.1016/j.plrev.2014.04.005

Frank, M. C., & Goodman, N. D. (2012, May). Predicting pragmatic reasoning in language games. Science, 336(6084), 998–998. Retrieved from https://doi.org/10.1126/science.1218633 doi: 10.1126/science.1218633

Geisler, W. S. (2011, April). Contributions of ideal observer theory to vision research. Vision Research, 51(7), 771–781. Retrieved from https://doi.org/10.1016/j.visres.2010.09.027 doi: 10.1016/j.visres.2010.09.027

Geisler, W. S., & Ringach, D. (2009, January). Natural systems analysis. Visual Neuroscience, 26(1), 1–3. Retrieved from https://doi.org/10.1017/s0952523808081005 doi: 10.1017/s0952523808081005

Geng, J. J., DiQuattro, N. E., & Helm, J. (2017). Distractor probability changes the shape of the attentional template. Journal of Experimental Psychology: Human Perception and Performance, 43(12), 1993–2007. doi: 10.1037/xhp0000430

Geng, J. J., & Witkowski, P. (2019, October). Template-to-distractor distinctiveness regulates visual search efficiency. Current Opinion in Psychology, 29, 119–125. Retrieved from https://doi.org/10.1016/j.copsyc.2019.01.003 doi: 10.1016/j.copsyc.2019.01.003

Goldreich, D. (2007, March). A bayesian perceptual model replicates the cutaneous rabbit and other tactile spatiotemporal illusions. PLoS ONE, 2(3), e333. Retrieved from https://doi.org/10.1371/journal.pone.0000333 doi: 10.1371/journal.pone.0000333

Goodale, M. A., & Milner, A. (1992). Separate visual pathways for perception and action. Trends in Neurosciences, 15(1), 20–25. doi: 10.1016/0166-2236(92)90344-8

Gordon, I. E. (1968). Interactions between items in visual search. Journal of Experimental Psychology, 76(3, Pt.1), 348–355. doi: 10.1037/h0025482

Gordon, I. E., Dulewicz, V., & Winwood, M. (1971). Irrelevant item variety and visual search. Journal of Experimental Psychology, 88(2), 295–296. doi: 10.1037/h0030888

Green, D., & Swets, J. (1966). Signal detection theory and psychophysics. John Wiley and Sons.

Harrison, W. J., & Bays, P. M. (2018, February). Visual working memory is independent of the cortical spacing between memoranda. The Journal of Neuroscience, 38(12), 3116–3123. Retrieved from https://doi.org/10.1523/jneurosci.2645-17.2017 doi: 10.1523/jneurosci.2645-17.2017

Howe, C. Q., & Purves, D. (2005, January). The muller-lyer illusion explained by the statistics of imagesource relationships. Proceedings of the National Academy of Sciences, 102(4), 1234–1239. Retrieved from https://doi.org/10.1073/pnas.0409314102 doi: 10.1073/pnas.0409314102

Jones, M., & Love, B. C. (2011, August). Bayesian fundamentalism or enlightenment? on the explanatory status and theoretical contributions of bayesian models of cognition. Behavioral and Brain Sciences, 34(4), 169–188. Retrieved from https://doi.org/10.1017/s0140525x10003134 doi: 10.1017/s0140525x10003134

Kiyonaga, A., & Egner, T. (2012). Working memory as internal attention: Toward an integrative account of internal and external selection processes. Psychonomic Bulletin & Review, 20(2), 228–242. doi: 10.3758/s13423-012-0359-y

Kleiner, M., Brainard, D., Pelli, D., Ingling, A., Murray, R., & Broussard, C. (2007). What’s new in psychtoolbox-3. Perception, 36(14), 1–16.

Kong, G., & Fougnie, D. (2019). Visual search within working memory. Journal of Experimental Psychology: General. doi: 10.1037/xge0000555

Kuo, B.-C., Rao, A., Lepsien, J., & Nobre, A. C. (2009). Searching for targets within the spatial layout of visual short-term memory. Journal of Neuroscience, 29(25), 8032–8038. doi: 10.1523/jneurosci.0952-09.2009

Levenstein, D., Alvarez, V. A., Amarasingham, A., Azab, H., Gerkin, R. C., Hasenstaub, A., … Redish, A. D. (2020). On the role of theory and modeling in neuroscience. arXiv. doi: arXiv:200313825

Lewandowsky, S., & Farrell, S. (2011). Computational modeling in cognition: Principles and practice. doi: 10.4135/9781483349428

Liu, G., Healey, C. G., & Enns, J. T. (2003). Target detection and localization in visual search: A dual systems perspective. Perception & Psychophysics, 65(5), 678–694.

Ma, W. J. (2012). Organizing probabilistic models of perception. Trends in Cognitive Sciences, 16(10), 511–518. doi: 10.1016/j.tics.2012.08.010

Ma, W. J. (2019, October). Bayesian decision models: A primer. Neuron, 104(1), 164–175. doi: 10.1016/j.neuron.2019.09.037

Ma, W. J., Beck, J. M., Latham, P. E., & Pouget, A. (2006, October). Bayesian inference with probabilistic population codes. Nature Neuroscience, 9(11), 1432–1438. Retrieved from https://doi.org/10.1038/nn1790 doi: 10.1038/nn1790

Ma, W. J., Navalpakkam, V., Beck, J. M., van den Berg, R., & Pouget, A. (2011). Behavior and neural basis of near-optimal visual search. Nature Neuroscience, 14(6), 783–790. doi: 10.1038/nn.2814

Ma, W. J., Shen, S., Dziugaite, G., & van den Berg, R. (2015). Requiem for the max rule? Vision Research, 116, 179–193. doi: 10.1016/j.visres.2014.12.019

Mazyar, H., van den Berg, R., & Ma, W. J. (2012). Does precision decrease with set size? Journal of Vision, 12(6), 10–10. doi: 10.1167/12.6.10

Mazyar, H., van den Berg, R., Seilheimer, R., & Ma, W. J. (2013). Independence is elusive: Set size effects on encoding precision in visual search. Journal of Vision, 13(5), 8–8. doi: 10.1167/13.5.8

Mueller, S. T., & Weidemann, C. T. (2008). Decision noise: An explanation for observed violations of signal detection theory. Psychonomic Bulletin & Review, 15(3), 465–494. doi: 10.3758/pbr.15.3.465

Nagy, A. L., Neriani, K. E., & Young, T. L. (2005). Effects of target and distractor heterogeneity on search for a color target. Vision Research, 45(14), 1885–1899. doi: 10.1016/j.visres.2005.01.007

Nagy, A. L., & Thomas, G. (2003). Distractor heterogeneity, attention, and color in visual search. Vision Research, 43(14), 1541–1552. doi: 10.1016/s0042-6989(03)00234-7

Najemnik, J., & Geisler, W. S. (2005, March). Optimal eye movement strategies in visual search. Nature, 434(7031), 387–391. doi: 10.1038/nature03390

Navalpakkam, V., & Itti, L. (2007, February). Search goal tunes visual features optimally. Neuron, 53(4), 605–617. Retrieved from https://doi.org/10.1016/j.neuron.2007.01.018 doi: 10.1016/j.neuron.2007.01.018

Nolte, L. W., & Jaarsma, D. (1967). More on the detection of one of m orthogonal signals. The Journal of the Acoustical Society of America, 41(2), 497–505. doi: 10.1121/1.1910360

Palmer, J. (1990). Attentional limits on the perception and memory of visual information. Journal of Experimental Psychology: Human perception and performance, 16(2), 332–350.

Palmer, J. (1994). Set-size effects in visual search: The effect of attention is independent of the stimulus for simple tasks. Vision Research, 34(13), 1703–1721. doi: 10.1016/0042-6989(94)90128-7

Palmer, J. (1995). Attention in visual search: Distinguishing four causes of a set-size effect. Current Directions in Psychological Science, 4(4), 118–123. doi: 10.1111/1467-8721.ep10772534

Palmer, J., Ames, C. T., & Lindsey, D. T. (1993). Measuring the effect of attention on simple visual search. Journal of Experimental Psychology: Human Perception and Performance, 19(1), 108–130. doi: 10.1037/0096-1523.19.1.108

Palmer, J., Verghese, P., & Pavel, M. (2000). The psychophysics of visual search. Vision Research, 40(10-12), 1227–1268. doi: 10.1016/s0042-6989(99)00244-8

Pelli, D. G. (1997). The VideoToolbox software for visual psychophysics: transforming numbers into movies. Spatial Vision, 10(4), 437–442.

Pertzov, Y., & Husain, M. (2013, September). The privileged role of location in visual working memory. Attention, Perception, & Psychophysics, 76(7), 1914–1924. Retrieved from https://doi.org/10.3758/s13414-013-0541-y doi: 10.3758/s13414-013-0541-y

Peterson, W., Birdsall, T., & Fox, W. (1954). The theory of signal detectability. Transactions of the IRE Professional Group on Information Theory, 4(4), 171–212. doi: 10.1109/tit.1954.1057460

Pratte, M. S. (2018). Swap errors in spatial working memory are guesses. Psychonomic Bulletin & Review. doi: 10.3758/s13423-018-1524-8

Rahnev, D., & Denison, R. N. (2018). Suboptimality in perceptual decision making. Behavioral and Brain Sciences, 41. Retrieved from https://doi.org/10.1017/s0140525x18000936 doi: 10.1017/s0140525x18000936

Rosenholtz, R. (1999, October). A simple saliency model predicts a number of motion popout phenomena. Vision Research, 39(19), 3157–3163. Retrieved from https://doi.org/10.1016/s0042-6989(99)00077-2 doi: 10.1016/s0042-6989(99)00077-2

Rosenholtz, R. (2001). Visual search for orientation among heterogeneous distractors: Experimental results and implications for signal-detection theory models of search. Journal of Experimental Psychology: Human Perception and Performance, 27(4), 985–999. doi: 10.1037/0096-1523.27.4.985

Rothkegel, L. O. M., Schütt, H. H., Trukenbrod, H. A., Wichmann, F. A., & Engbert, R. (2019, February). Searchers adjust their eye-movement dynamics to target characteristics in natural scenes. Scientific Reports, 9(1). Retrieved fromhttps://doi.org/10.1038/s41598-018-37548-w doi: 10.1038/s41598-018-37548-w

Schwarz, G. (1978). Estimating the dimension of a model. The Annals of Statistics, 6(2), 461–464.

Shaw, M. L. (1982). Attending to multiple sources of information: I. the integration of information in decisionmaking. Cognitive Psychology, 14(3), 353–409. doi: 10.1016/0010-0285(82)90014-7

Shen, S., & Ma, W. J. (2016). A detailed comparison of optimality and simplicity in perceptual decision making. Psychological Review, 123(4), 452–480. doi: 10.1037/rev0000028

Shen, S., & Ma, W. J. (2019). Variable precision in visual perception. Psychological Review. doi: 10.1101/153650

Stengård, E., & van den Berg, R. (2019). Imperfect bayesian inference in visual perception. PLOS Computational Biology, 15(4), e1006465. doi: 10.1371/journal.pcbi.1006465

Tamber-Rosenau, B. J., Fintzi, A. R., & Marois, R. (2015). Crowding in visual working memory reveals its spatial resolution and the nature of its representations. Psychological Science, 26(9), 1511–1521. doi: 10.1177/0956797615592394

Treisman, A., & Gelade, G. (1980). A feature-integration theory of attention. Cognitive Psychology, 12(1), 97–136. doi: 10.1016/0010-0285(80)90005-5

Treisman, A., & Gormican, S. (1988). Feature analysis in early vision: Evidence from search asymmetries. Psychological Review, 95(1), 15–48. doi: 10.1037/0033-295x.95.1.15

van den Berg, R., Shin, H., Chou, W.-C., George, R., & Ma, W. J. (2012). Variability in encoding precision accounts for visual short-term memory limitations. Proceedings of the National Academy of Sciences, 109(22), 8780–8785. doi: 10.1073/pnas.1117465109

van den Berg, R., & Ma, W. J. (2018, August). A resource-rational theory of set size effects in human visual working memory. eLife, 7. Retrieved from https://doi.org/10.7554/elife.34963 doi: 10.7554/elife.34963

van Opheusden, B., Acerbi, L., & Ma, W. J. (2020). Unbiased and efficient log-likelihood estimation with inverse binomial sampling. arXiv. doi: arXiv:2001.03985

Verghese, P. (2001). Visual search and attention. Neuron, 31(4), 523–535. doi: 10.1016/s0896-6273(01)00392-0

Verghese, P., & Nakayama, K. (1994). Stimulus discriminability in visual search. Vision Research, 34(18), 2453–2467. doi: 10.1016/0042-6989(94)90289-5

Verghese, P., & Stone, L. S. (1995). Combining speed information across space. Vision Research, 35(20), 2811–2823. doi: 10.1016/0042-6989(95)00038-2

Vincent, B., Baddeley, R., Troscianko, T., & Gilchrist, I. (2009). Optimal feature integration in visual search. Journal of Vision, 9(5), 15–15. doi: 10.1167/9.5.15

Weiss, Y., Simoncelli, E. P., & Adelson, E. H. (2002, May). Motion illusions as optimal percepts. Nature Neuroscience, 5(6), 598–604. Retrieved from https://doi.org/10.1038/nn0602-858 doi: 10.1038/nn0602-858

Wigner, E. (1960, April). The unreasonable effectiveness of mathematics in the natural sciences. Communications of pure and applied mathematics, 13, 001–14.

Wolfe, J. M. (1994). Visual search in continuous, naturalistic stimuli. Vision Research, 34(9), 1187–1195. doi: 10.1016/0042-6989(94)90300-x

Wolfe, J. M. (2010, April). Visual search. Current Biology, 20(8), R346–R349. Retrieved from https://doi.org/10.1016/j.cub.2010.02.016 doi: 10.1016/j.cub.2010.02.016

Wulff, M., Stainton, A., & Rotshtein, P. (2016). Effects of paired-object affordance in search tasks across the adult lifespan. Brain and Cognition, 105, 22–33. doi: 10.1016/j.bandc.2016.03.009

Yang, S. C.-H., Lengyel, M., & Wolpert, D. M. (2016). Active sensing in the categorization of visual patterns. eLife, 5. doi: 10.7554/elife.12215

